# Bird population trend analyses for a monitoring scheme with a highly structured sampling design

**DOI:** 10.1101/2024.06.30.601382

**Authors:** Mirjam R. Rieger, Christoph Grüneberg, Michael Oberhaus, Sven Trautmann, Madalin Parepa, Nils Anthes

## Abstract

Population trends derived from systematic monitoring programmes are essential to identify species of conservation concern and to evaluate conservation measures. However, monitoring data pose several challenges for statistical analysis, including spatial bias due to an unbalanced sampling of natural regions or habitats, variation in observer expertise, frequent observer changes, and overdispersion or zero-inflation in the raw data. An additional challenge arises from so-called ‘rolling’ survey designs, where each site is only visited once within each multi-year rotation cycle. We developed a GAMM-based workflow that addresses these challenges and exemplify its application with the highly structured data from the Ecological Area Sampling (EAS) in the German federal state North Rhine-Westphalia (NRW). First, we derive a routine that allows informed decisions about the most appropriate combination of distribution family (Poisson or negative binomial), model covariates (e.g., habitat characteristics), and zero-inflation formulations to reflect species-specific data distributions. Second, we develop a correction factor that buffers population trend estimates for variation in observer expertise as reflected in variation in total bird abundance. Third, we integrate region-specific trends that adjust for between-year variation in the representation of habitat or natural regions within the yearly subset of sampled sites. In a consistency check, we found good match between our GAMM-based EAS trends and TRIM-based trends from the German Common Bird Monitoring scheme. The study provides a template script for *R* statistical software so the workflow can be adapted to other monitoring programmes with comparable survey designs and data structures.

## 1 Introduction

Bird populations decline worldwide (Klvaňová et al. 2009, Sauer et al. 2017, Burns et al. 2021a, Burns et al. 2021b), with the European Union alone facing an approximate loss of 560–620 million bird individuals (17–19%) within 40 years (Burns et al. 2021a, Burns et al. 2021b). Such knowledge of population sizes and their trends heavily relies on standardised monitoring programmes to provide reliable population density estimates. The resultant population trend estimates allow allocating conservation resources to species of highest conservation concern (Niemelä 2000, Buckland and Johnston 2017), provide feedback on the efficiency of conservation efforts (Johnston et al. 2015), and raise awareness for the value and state of biodiversity among decision makers and the public (Jennings 2021).

However, many monitoring programmes pose substantial challenges for statistical inference that can reduce the reliability and robustness of the estimated trends if not treated with care (Buckland and Johnston 2017). Our study focuses on a monitoring scheme with a rolling (or ‘rotating’) survey design that generates multi-year intervals between repetitive surveys per site (Buckland and Johnston 2017). Rolling surveys can cover more study sites and thus a wider range of habitats or natural regions across years than monitoring schemes with a yearly coverage of all sites but result in a large fraction of sites with missing values each year and frequent observer changes. In this context, we develop an analytical protocol to cover three core aspects. First, rolling monitoring programmes rest on spatially structured sampling designs, often coupled with between-year variation in the representation of habitat types, which needs integration into the analysis (van Strien et al. 2004, Buckland and Johnston 2017). Spatial bias arises when relevant natural regions or habitats are not represented according to their spatial coverage in the monitoring sample (van Strien et al. 2004). As a result, when population trends vary between natural regions or habitats, the estimated overall population trend can be biased towards overrepresented natural regions, or reflect changes in sample composition rather than a true change in abundance (Buckland and Johnston 2017, Bowler et al. 2022).

Second, detection probabilities and survey quality can vary strongly among and (over time) within observers, potentially introducing random and systematic trend estimation errors when ignored (Sauer et al. 1994, Link and Sauer 1998, Kéry et al. 2005, Jiguet 2009, Farmer et al. 2014). When average observer expertise remains constant, between-observer variation primarily increases (random) variation in abundance estimates between sites and years. (Sauer et al. 1994, Johnston et al. 2018). In contrast, within-observer learning and systematic changes in average observer expertise introduce (systematic) biases in trend estimates. For example, initial familiarisation with a new study site and the specificities of the monitoring programme usually cause underestimated bird abundances in early survey years (Link and Sauer 1998, Jiguet 2009). Moreover, observer expertise usually increases with time after active engagement in bird territory surveys, while species detectability declines with age if hearing impairment reduces the detection of high-pitch bird vocalisations (Farmer et al. 2014). In rolling survey design, these general issues typically go along with a high turnover in observer identities between successive surveys of the same site, which adds further variation.

Finally, count response data as typical for bird census require models to reflect distribution properties such as overdispersion or zero inflation (Blasco-Moreno et al. 2019, Campbell 2021). Overdispersion arises when the variance of count data exceeds their mean value. It is typically resolved by modelling trends with a negative binomial error distribution instead of a Poisson distribution (Blasco-Moreno et al. 2019, Campbell 2021). Zero-inflation occurs when datasets contain so-called structural zeros. This implies a disproportionally large fraction of unoccupied sites that cannot be explained solely by sampling variation among sites considered suitable, but rather arises from survey sites that are unsuitable. Where present, structural zeros are modelled separately from the count distribution in zero-inflated model components (Korner-Nievergelt et al. 2015, Blasco-Moreno et al. 2019, Campbell 2021, Tirozzi et al. 2021).

The software TRIM (Pannekoek and van Strien 2001) represents a common analytical tool for the analysis of monitoring data and is used, for example, in the Pan European Common Bird Monitoring Scheme (Vorisek et al. 2008) and the national bird monitoring programmes of Sweden (Jiguet et al. 2013), Finland (Pöysä et al. 2013), or Germany (Kamp et al. 2021). TRIM computes trends and annual population indices from loglinear Poisson regressions. It corrects for overdispersion, serial correlation and missing values (van Strien et al. 2004) and allows weights to account for spatial bias (van Strien et al. 2004). Yet, designed for monitoring schemes with almost yearly surveys per site by the same observer, TRIM estimates are considered robust unless the turnover in survey sites between years leads to a ≥ 60 % fraction of missing values (van Strien et al. 2001, Bogaart et al. 2020, Dakki et al. 2021). Moreover, TRIM is restricted to categorical covariates, requiring climate or landscape composition covariates to be transformed into categories (Bogaart et al. 2020). Finally, long-term trend analyses beyond yearly index estimates are restricted to linear trends and breakpoint analyses, while more recent developments favour the integration of trend smoothers, e.g. with general additive (mixed) models (GA(M)Ms) (e.g., Fewster et al. (2000), Knape (2016)). Using a smoothing function, GA(M)Ms can also capture non-linear short- and long-term trends while allowing to identify periods of strongly increasing or decreasing trends and breakpoints of (linear) trend direction (Fewster et al. 2000, Zuur 2012, Knape 2016, Wood 2021).

We develop a user-friendly tool that integrates high observer turnover, long time gaps between successive surveys, spatial bias, as well as overdispersion and zero-inflation into an analysis of bird abundance trends from rolling surveys. It is exemplified with data from the Ecological Area Sampling (EAS), a monitoring programme of the German federal state North Rhine-Westphalia (NRW) with a six-year rolling sampling scheme. As a quality check, we compared the predicted population trends with West German trends derived from the German Common Bird Monitoring scheme (MhB) with yearly surveys per site. The statistical approach is provided as an *R* script (Rieger et al. 2025) and package (Rieger 2024) that can be adjusted to other datasets with a comparable structure.

## 2 Material and Methods

### 2.1 The Ecological area sampling (EAS) dataset

The Ecological Area Sampling (EAS) is a long-term biodiversity monitoring programme of the State Agency for Nature, Environment and Climate in North Rhine-Westphalia (LANUK, former LANUV) with 170 study sites that represent the average landscape of North Rhine-Westphalia (NRW) (LANUV 2016b). Sites are distributed across six natural regions and two biogeographic regions (atlantic lowlands, continental highlands) (Fig. 1) proportional to their spatial coverage. Another 21 metropolitan sites were added in 2011 to represent habitat characteristics of the Rhine-Ruhr metropolis region (Weiss and Schulze-Hagen 2014). The program targets an alternating six-year cycle for successive surveys of a given site but could not yet strictly impose it given restructuring in the yearly balance of sites among natural and metropolitan regions (see section 2.3). The dataset for the survey period 2002-2020 typically covered 25 to 36 (max. 47) sites per year, two to four replicated surveys per site, and 614 surveys in total. Each site is visited at least nine times per sampling year (two to eight hours per date) between February and July with complete coverage of its 1 km^2^ area (LANUV 2016a). The repetitive surveys are used to derive the territory count per species and km^2^, which represents the response variable *abundance* in all statistical analyses (LANUV 2016a). Survey effort (survey number and duration) constitute relevant model covariates but the respective information is not available for the current dataset (Further detail on EAS methodology in appendix section 1).

**Figure 1:**
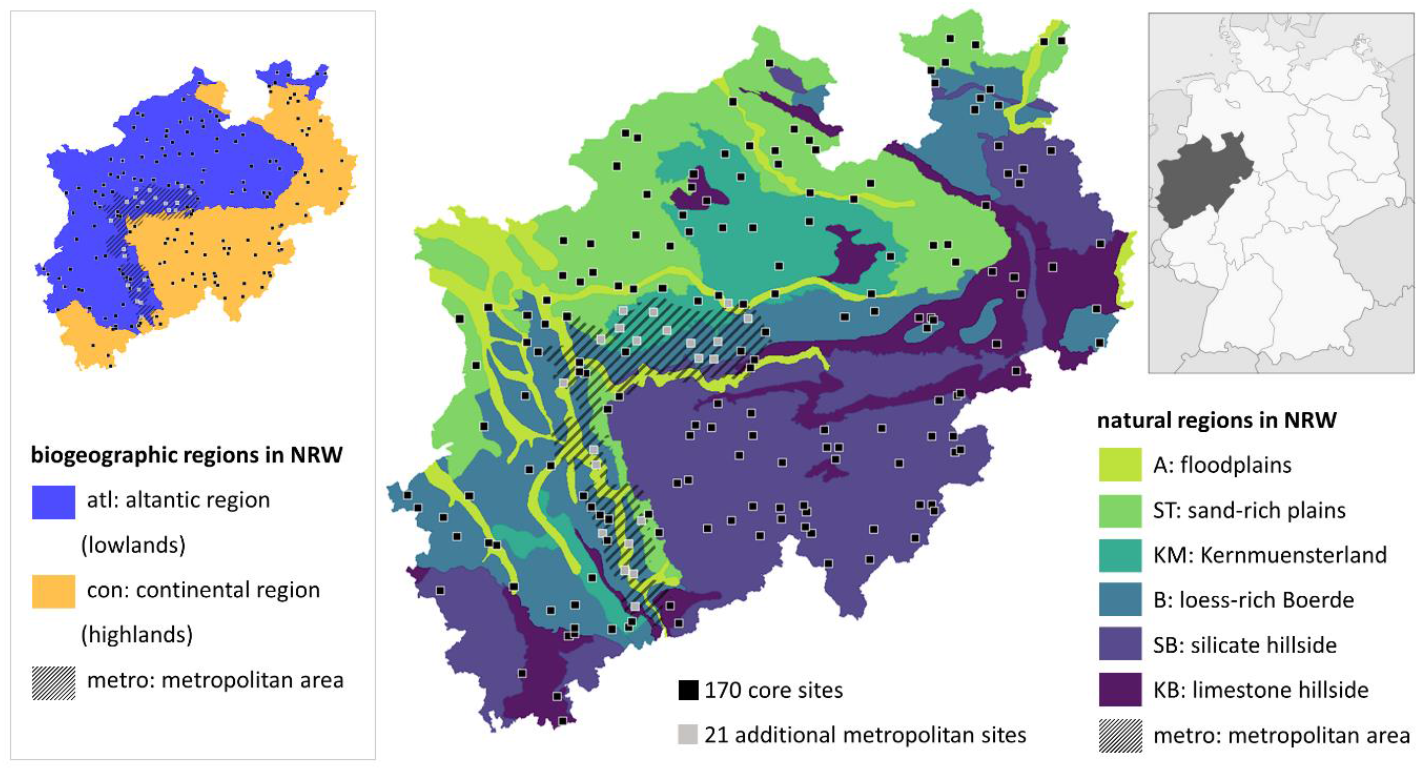
Sites and study area of EAS coloured by six natural regions (middle) and two biogeographic regions (left), both including the Rhine-Ruhr metropolis, which was implemented as seventh natural region. Map adapted from LANUV (2016b).

### 2.2 Site characterisation

Sites were distributed into six natural regions and whether they were located in a metropolitan area or not (Fig. 1). We combined these two layers of information to produce a new *natural regions* variable with seven levels, all sites within the metropolitan area being classified as “metropolitan area” while others were classified according to the natural region they belong to. Another site-specific categorical variable was *biogeographic region* (atlantic, continental). Natural regions showed near-complete separation between biogeographical regions, where the continental region encompasses the natural regions ‘silicate hillside’ and ‘limestone hillside’ and the Atlantic region the remaining five natural regions including the ‘metropolitan area’ (Fig. 2 right). We therefore only considered *biogeographic region* in the post-modelling process (see section ‘Population trends’).

**Figure 2:**
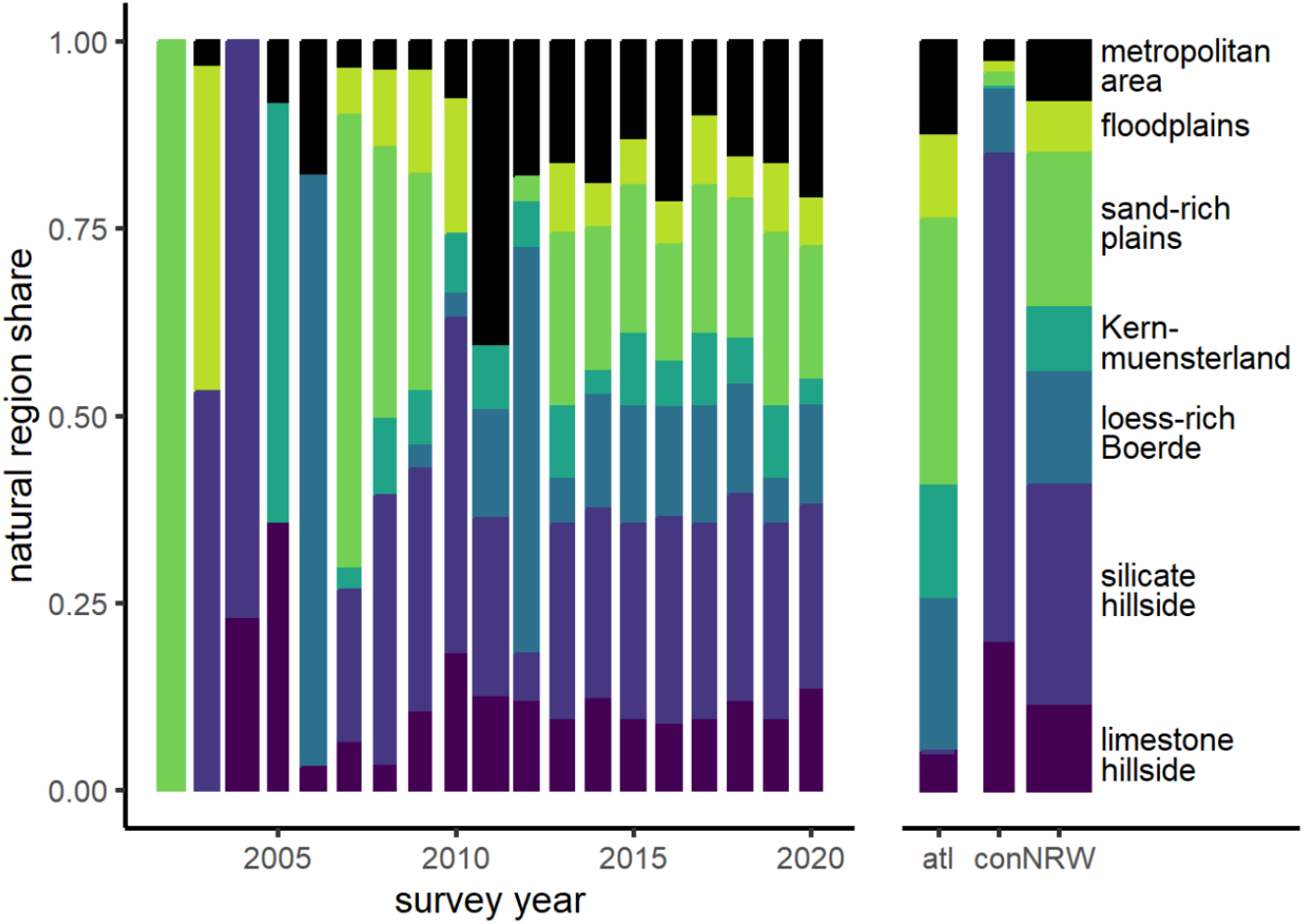
Natural region shares in annual EAS samples (main panel) and their spatial coverage per biogeographic region (atlantic - atl, continental - con) and in NRW (right panel). Bar widths are proportional to the number of surveyed sites per year.

Quantitative site characteristics included *altitude* above sea level (ATKIS 2013), ten parameters of *climate* and three of *landscape compositions* to account for variation in mean environmental attributes between the site subsets surveyed per year. Climate parameters included long-term spring and winter averages between 1981 and 2010 (CDC 2010) of minimum, mean, and maximum temperatures, precipitation, and sunshine duration. Landscape composition variables were the coverages of forest, arable land, and settlements (ATKIS 2013). Given substantial collinearity between these 14 attributes, we integrated their principal components (PC) with eigenvalues exceeding 1 into the statistical models (three PCs in our case; for details see Appendix section 2).

### 2.3 Spatial bias

The initial sample of 170 EAS monitoring sites almost perfectly represented the proportional coverage of natural regions in NRW, but sampling bias arose from two sources. First, the monitoring programme added 21 metropolitan sites overrepresenting the metropolitan area to the other six natural regions in 2011. Second, the yearly subsampling of sites was highly unbalanced with respect to natural regions in the early phase of the EAS programme (Fig. 2), with improvements since 2007 and almost perfect balance since 2013.

To account for the uneven allocation in early programme years and for the overrepresentation of metropolitan areas since 2011, we analysed annual trends per natural region.

### 2.4 Observer effects

The delineation of breeding territories is prone to under- and overestimation arising from variation in observer expertise in species identification and detection, territory delineation, and general field survey quality (Südbeck et al. 2005, Johnston et al. 2018), even in programmes such as the EAS that hire skilled ornithologists (König 2020). Observer effects add further noise to trend estimates when observers change between successive surveys of a given site. Excluding data from first year surveys, an established standard in monitoring data with yearly site coverage (e.g., German Common Bird Monitoring (Kamp et al. 2021)), is unfeasible for rolling surveys: In the EAS dataset, 434 out of 614 breeding bird surveys between 2002 and 2020 (71 %) were such first-year combinations of observer and site. To characterise observer expertise and learning effects, we therefore categorised observers as (i) unfamiliar with the survey programme and site, (ii) familiar with survey programme but unfamiliar with the site, or (iii) familiar with both. Yet, we excluded this predictor from final models since it did not improve model fit, most likely because site familiarisation introduced no detectable bias into the EAS trend estimates given the high observer turnover rates among successive surveys.

Instead, we identified surveys with suspiciously high or low territory counts summed across all species. The procedure assumes that severe observer effects typically manifest in a general under- or overestimation of territory numbers (= abundance), so that total abundance (summed across species) for a given site and year stands out against average total abundance for that site. Based on this logic, we calculated

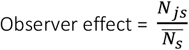

with *N*_*js*_ the total abundance across species on site *s* in survey year *j*, and 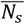 the mean total abundance across species on site *s* across all survey years (Rieger 2024). This observer effect is a ratio, so that a value of 1 indicates no deviation from the mean, and a value of 0.75 (1.25) a deviation of -25% (+25%) from the observed mean. We integrated the *observer effect* as a categorical predictor variable classified as ‘negative’ (or ‘positive’) when the abundance sum of a given survey was at least 25% smaller (or larger) than the mean per site, and as ‘none’ otherwise. As a sensitivity analysis, we also checked observer effect thresholds of 20% and 15%.

### 2.5 Distribution characteristics and the statistical analysis of EAS

We analysed species-specific abundance trends between 2002 and 2020 for all species with ≤ 90 % zero-sightings in both biogeographic regions (and thus at least 61 non-zero records) to allow plausible trend detection. The criterion was fulfilled in 61 (out of 148) species. For conciseness, this study focuses on a subset of 14 species that cover the observed range of abundance distributions and proportions of zero counts (Table 1).

**Table 1:**
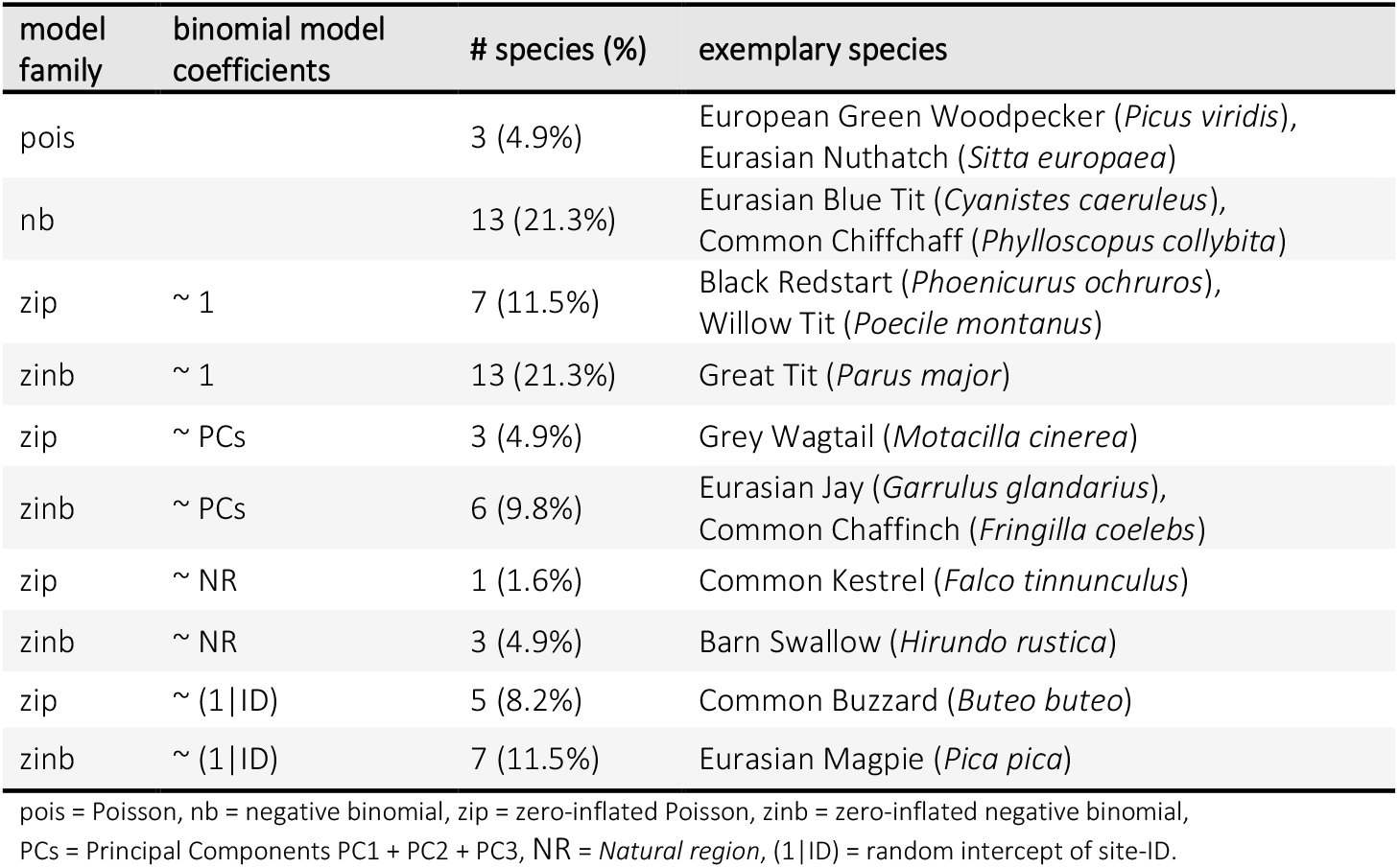
Overview of model types (main model family, coefficients in the optional zi-binomial model) and number of case study species per model type. Species names are given for the 14 exemplary species shown in this paper. Details for all species are in Appendix section 7. Models are ordered according to their parsimony (top = most parsimonious).

Trends in *Abundance*, i.e. the territory count per site and year, were analysed using generalised additive mixed models (GAMMs) within a Bayesian framework as implemented in the package *brms* (v.2.21.0 (Bürkner 2017)) for R (v.4.3.3 (R Core Team 2024)). The *brms* package fits Bayesian models using *Stan* via the package *cmdstanr* (v.0.7.1 (Gabry and Češnovar 2020)) and allows to more flexibly integrate the required error families and model structures. We implemented four different error structures to account for zero-inflation or overdispersion, namely a Poisson error structure (Pois), negative binomial (nb), zero-inflated Poisson (zip), or zero-inflated negative binomial (zinb), all using a log link (Table 2, Appendix section 3). Since the underlying survey method generates .5 abundances for peripheral territories, we multiplied *abundance* by two (so it would be number of territories for 2 km^2^) and added an *offset* of two to all models.

**Table 2:** Examples for model assessment and procedure to select species-specific models from six alternative model structures (for further species and models see Appendix section 3). Models differ in family and the presence of a zero-inflated model component (zinb and zip), the latter also in binomial model coefficients. Model assessment quantifies whether simulated data from the model is consistent with the observed response variable (abundance) with respect to its proportion of zeros (propZ), mean and maximum value, and standard deviation (SD). Bayesian p-values close to 0.5 indicate that the observed values are central within the posterior distribution of simulated raw datasets (8000 in total), indicating good model fit. Values close to 0 and 1 indicate substantial under- or overestimation and thus poor model fit. Final model selection is based on ELPD differences by selecting the most parsimonious model (bold, top to bottom) out of all models with an ELPD difference ± 1.96*SE of ELPD difference including 0 (bold values). Abbreviations see Table. 1.

| species | model family | binomial model coefficients | Bayesian p-value |  |  |  | ELPD |  |
| --- | --- | --- | --- | --- | --- | --- | --- | --- |
|  |  |  | propZ | mean | SD | max | diff. | se diff. |
| European Green Woodpecker<br>*Picus viridis* | pois |  | 0.01 | 0.49 | 0.79 | 0.65 | **-11.86** | **7.89** |
|  | nb |  | 0.01 | 0.42 | 0.62 | 0.68 | -204.94 | 47.02 |
|  | zip | ~ 1 | 0.12 | 0.44 | 0.87 | 0.72 | **-6.65** | **6.77** |
|  | zinb | ~ 1 | 0.13 | 0.46 | 0.91 | 0.80 | **-8.06** | **6.73** |
|  | zip | ~ PCs | 0.16 | 0.46 | 0.88 | 0.71 | **-3.66** | **6.92** |
|  | zinb | ~ PCs | 0.16 | 0.48 | 0.90 | 0.76 | -18.57 | 7.91 |
| Eurasian Blue Tit<br>*Cyanistes caeruleus* | pois |  | 0 | 0.50 | 0.01 | 0.02 | -510.17 | 69.13 |
|  | **nb** |  | 0 | 0.64 | 0.70 | 0.37 | **-11.2** | **5.77** |
|  | zip | ~ 1 | 0.10 | 0.34 | 0.01 | 0.02 | -498.39 | 66.54 |
|  | zinb | ~ 1 | 0.07 | 0.58 | 0.69 | 0.37 | -19.9 | 8.09 |
|  | zip | ~ PCs | 0.11 | 0.39 | 0.01 | 0.02 | -509.75 | 71.86 |
|  | zinb | ~ PCs | 0.07 | 0.60 | 0.70 | 0.38 | **-6.2** | **5.83** |
| Black Redstart<br>*Phoenicurus ochruros* | pois |  | 0 | 0.51 | 0.56 | 0.47 | -23.37 | 10.14 |
|  | nb |  | 0 | 0.59 | 0.82 | 0.59 | -3654.6 | 780.81 |
|  | **zip** | **~ 1** | 0 | 0.30 | 0.48 | 0.45 | **-8.51** | **9.42** |
|  | zinb | ~ 1 | 0 | 0.42 | 0.76 | 0.56 | **-4.01** | **7.52** |
|  | zip | ~ PCs | 0 | 0.42 | 0.54 | 0.48 | **-12.87** | **9.19** |
|  | zinb | ~ PCs | 0 | 0.51 | 0.79 | 0.51 | **0** | **0** |
| Eurasian Jay<br>*Garullus glandarius* | pois |  | 0 | 0.50 | 0.76 | 0.77 | -37.68 | 11.47 |
|  | nb |  | 0 | 0.69 | 0.89 | 0.70 | -117.44 | 22.38 |
|  | zip | ~ 1 | 0 | 0.31 | 0.78 | 0.76 | -27.5 | 8.37 |
|  | zinb | ~ 1 | 0 | 0.34 | 0.83 | 0.81 | -28.52 | 8.11 |
|  | zip | ~ PCs | 0.19 | 0.36 | 0.77 | 0.67 | -23.57 | 10.83 |
|  | **zinb** | **~ PCs** | 0.19 | 0.39 | 0.81 | 0.72 | **-15.08** | **10.25** |

The main model was constructed as follows, and we explain each model component below:

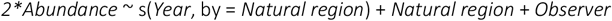

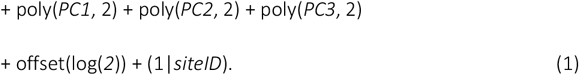

Survey *Year* as the main continuous predictor was modelled with a smoothing function (thin plate regression splines with an automatic selection of the degree of smoothing and k = 10 default basis dimension, (Wood 2021)) to account for non-linear changes in abundance over time while allowing adjacent years to be related to each other (Fewster et al. 2000, Zuur 2012, Knape 2016, Wood 2021). Thus, smoothers buffer against the property of rolling surveys that sampled site subsets differ between successive years, so that year-to-year changes in abundance cannot differentiate among-site variation from true short-term change in population size (Zuur 2012). Smoothers were modelled separately per *Natural region* to allow detecting regional differences in population trends and – for the overall trend estimates – accounting for spatial bias that may arise from unbalanced sampling design among regions (see above) in the post-modelling process (Fig. 2).

We added *Natural region* as a categorical main effect (Wood 2021) to contrast population densities and trends between the seven natural regions. Since natural regions showed near-complete separation between biogeographical regions (see Fig. 2 right and section ‘Site characterisation’), we therefore refrained from adding *biogeographic region* as a separate predictor, but extract population trends per biogeographic region by combining the respective natural region trends (see section ‘Population trends’).

*Observer* effects as outlined in section 2.4 were added as a categorical covariate. For numeric covariates, we included linear and quadratic terms of site-specific environmental attributes captured in the first *three principal components PC1* to *PC3* as outlined in section 2.2 and Appendix section 2. Finally, we added *site-ID* as a random intercept to reflect repeated measures per site and to model variation in mean abundance between the different site subsets per year.

The zero-inflation model component, where needed (Table 1), represents a binomial (Bernoulli) model with a logit link to estimate the additional occurrence of structural zeros. We kept this component as simple as possible (formulation 2.1) and estimated separate zero-inflation parameters per *Natural region*, environmental *PC*, or study *site-ID* (formulations 2.2–2.4) only where needed to improve model convergence and better reflect the observed distribution of zero counts (Fig. 3, Table 2):

**Figure 3:**
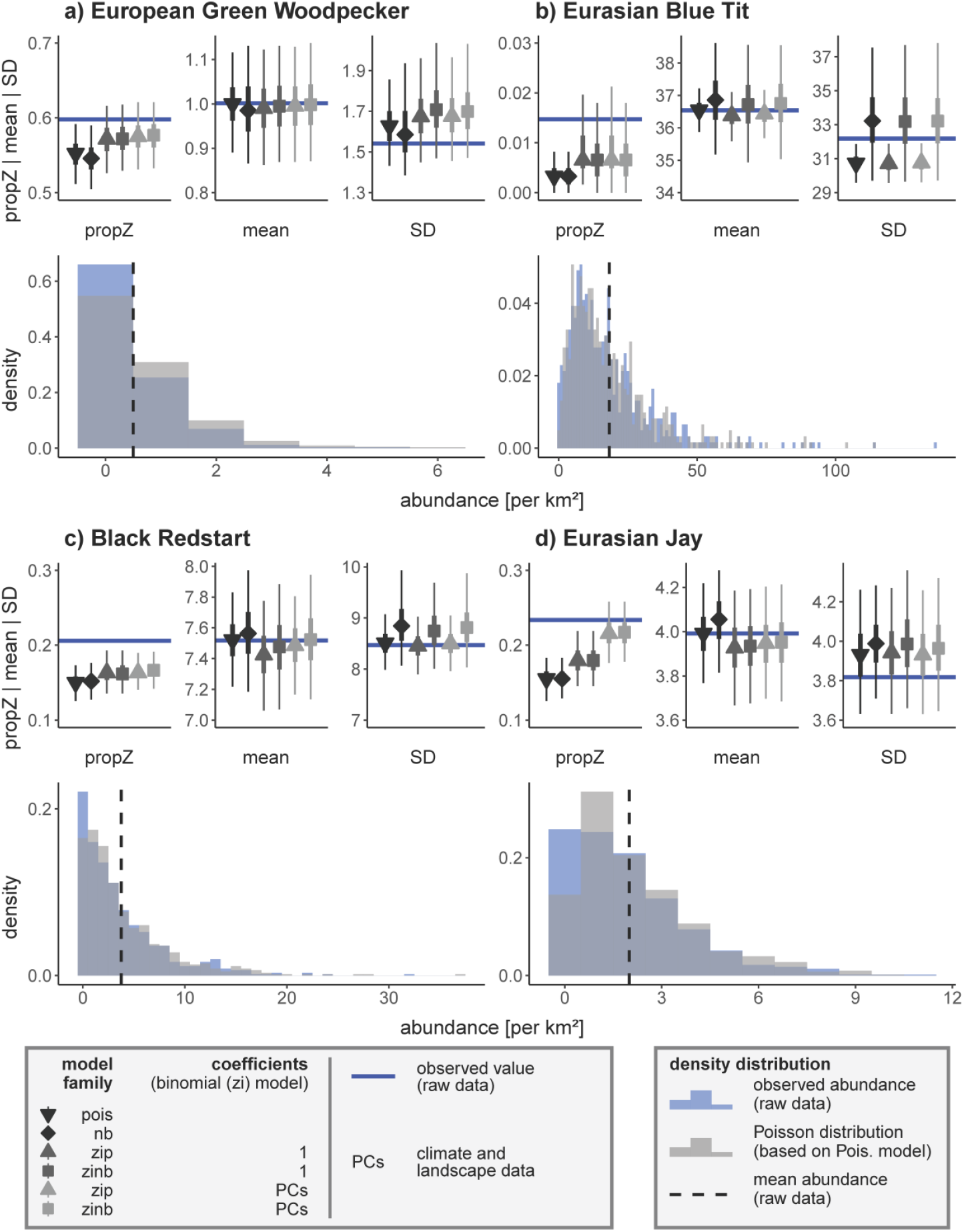
Reasoning for selecting appropriate model structures for four species. Upper panels: Model performance with respect to the proportion of zero counts (propZ), mean, and standard deviation (SD) of the response variable ‘abundance’. Symbols with error bars display the median ± 50% CrI (thick bars) and 95% CrI (thin bars) for 8000 posterior raw datasets simulated from the selected models as given in the legend. Optimal models have a close match between observed and model-predicted data. Lower panels: Density distributions of observed abundances (blue, dashed line = mean) and Poisson distributions (grey) based on simulated responses derived from the respective Poisson model. Y-axes differ in scale.

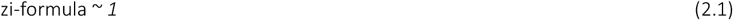

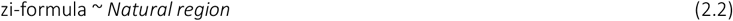

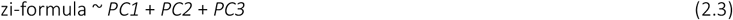

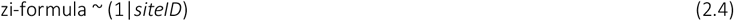

We did not integrate temporal autoregressive structures (van Strien et al. 2004, Korner-Nievergelt et al. 2015) because variograms plotted with the *variog* function of the package *geoR* (v.1.9-3 (Ribeiro Jr and Diggle 2020)) did not reveal issues with temporal autocorrelation. This may result from the difficulty of detecting autocorrelation with just 2-4 observations per site as well as from the six-year gaps between repeated measures (usually accompanied by a change in observer), which may add more variation to model residuals than expected from the remaining temporal pattern.

To fit Bayesian models, we used weakly informative prior distributions for coefficients (normal with mean = 0, SD = 2.5 for categorical coefficients, mean = 0, SD = 10 for numeric coefficients) and a maximum tree depth of 10 for the Hamiltonian Monte Carlo algorithm in Stan. Each model ran four Markov chain Monte Carlo chains with 1,000 warm-up and 2,000 post-warm-up samples per chain. We used the 8,000 post-warm-up samples for posterior predictive checks of the model’s ability to simulate the observed *abundance* distribution. Criteria included distributional characteristics (mean and standard deviation) as well as the model’s ability to reflect extreme values (maximum value) and potential zero inflation (proportion of zero counts, propZ) in the data. In all these cases, we calculated the proportion of simulations exceeding the observed raw data value (Bayesian p-value, Korner-Nievergelt et al. 2015). Bayesian p close to 0.5 imply a good match because the observed values are roughly central within the distribution of simulated data, while values approaching 0 or 1 flag models that clearly under- or overestimated the observed value.

We further checked several conversion indicators of the coefficient estimates: R_hat_-values ≤ 1.1 (observed max = 1.01) (Brooks and Gelman 1998), effective posterior samples ≥ 10% of total samples (observed min = 9.4%) and Monte Carlo standard errors ≤ 10% of the standard deviation (observed max = 4.3%) (Korner-Nievergelt et al. 2015). To select the model with the best predictive performance per species, we did a k-fold-cross-validation using 16 folds resulting in a theoretical expected log pointwise predictive density (ELPD) per model, as well as ELPD differences (model with best ELPD compared to others) and the respective standard error of the difference (se ELPD difference) (Bürkner 2017). When several models resulted in near-identical performance (ELPD difference ± 1.96*se ELPD difference included 0), we continued with the most parsimonious model, i.e., the model with the lowest number of covariates or model components (see Table 1), that fulfilled our conversion indicators. Note that cross-validation leads to a high computational cost: Runtime per model and species without cross-validation span between just one and five minutes, whereas cross-validation (16 additional models) took 15 to 60 minutes per species on a machine with 16 threads.

From the post-warm-up samples we also derived mean covariate coefficient estimates with their 95% credible intervals (CrI) and the posterior probability that the estimate exceeds 0, P(β > 0). Posterior probabilities approaching 0 or 1 indicate increasingly strong evidence for a directional coefficient.

#### Population trends

Model predictions ± 95% CrI were derived via brms.fitted (Bürkner 2017) to display trends graphically. When estimating trends over years and differences between natural regions, values of the remaining model covariates were set to their natural region-specific sample mean (continuous covariates), to the level ‘none’ (for ‘observer effect’) (Korner-Nievergelt et al. 2015), and group-level effects (‘site ID’) were generalised beyond the specific grouping level by using the argument re_formula = NA. We derived trends across biogeographic regions by combining these natural region-specific trends according to the natural region’s share of the biogeographic region (Atlantic (continental): A = 11.7% (2.1%), B = 20.9% (9.2%), KB = 5.1% (20.1%), KM = 15.1% (0.4%), SB = 0.0% (64.7%), ST = 35.1% (1.2%), metro = 12.1% (2.3%)). Overall trends were derived alike by combining the region-specific trends according to their share of NRW (55.5% Atlantic region and 44.5% continental region). Based on pairwise differences of predicted abundances, we estimated trends as changes in abundance per km^2^ between the start and end date of any desired time period as

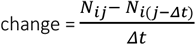

with *N* the predicted abundance per km^2^ for the *i*th posterior draw at times *j* and *j* − *Δt*, where *Δt* is the desired time period. If the 95% CrI of these changes in abundance excluded 0, the trend estimate was considered robust. We specifically produced pairwise differences to estimate annual changes (*Δt* = 1) and a mean long-term trend between 2008 and 2019 (*Δt* = 12, *j* = 2019).

#### Population index

For comparison with other monitoring programmes, we integrated the option to transform predicted mean abundances per km^2^ into an index relative to a user-selected baseline year (here: 2006). Even though the first few years of the monitoring programme suffer from non-representative sampling (see section 2.3), weights and smoothing-function enabled accurate estimates in these years without distorting successive estimates (Buckland and Johnston 2017). Abundance indices were calculated as

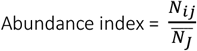

with *N*_*ij*_ the predicted abundance of the *i*th posterior draw in year *j*, and 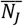 the mean predicted abundance of all simulations in the baseline year *J* (here *J* = 2006). Based on 10,000 simulations, we calculated the mean index and its 95% CrI for each time step *j*. Note that this computation calculates an uncertainty also for the baseline year *J*, which implies that confidence intervals display the uncertainty of the calculated index value and not of the change in index relative to the reference year.

Users who prefer a fixed baseline year index without uncertainty need to replace the mean abundance denominator,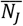 in the formula by *N*_*iJ*_, i.e. the *i*th simulation in the baseline year.

#### Data exclusion

Based on an internal validation process of the LANUK, some abundance data were classified as implausible and therefore excluded from the analysis. This process was species-dependent, leading to sample sizes of 608 to 614 per species (full dataset: 614, Appendix section 7).

### 2.6 Consistency check

We checked our EAS trends for NRW for consistency with population trends in all West Germany according to the German Common Bird Monitoring scheme (MhB), coordinated by the German association for field ornithologists. MhB study sites are ideally surveyed each year by the same observer (four surveys per site between 10 March and 20 June; full programme details in Kamp et al. (2021)).

The consistency analysis included 59 widely distributed species spanning a broad range of habitat associations and breeding strategies for which we had trend estimates available from the MhB and from the EAS. It is obviously difficult to compare trend estimates between programmes that rest on different methodologies and survey efforts (nine vs. four surveys per year), have different spatial coverage (NRW vs. West Germany) with just a partial match in study sites, and employ fundamentally different analytical approaches (GAM smoothers in EAS vs. yearly point estimates in MhB). Nevertheless, largely consistent estimates from EAS and MhB would reassure that the analytical approach developed for rolling sampling strategies produces reliable trends that are robust to the underlying survey design. With these limitations in mind, we expected that yearly samples in the MhB versus 6-yearly samples in the EAS should generate largely consistent long-term trend estimates, except for cases where a species’ biology would objectively favour one survey and analysis routine over the other (see Discussion).

To assess trend consistency we calculated two parameters for each species: (i) a Pearson ‘correlation’ coefficient between the annual TRIM-based MhB indices and the annual EAS-indices extracted from model smoothers between 2005 and 2020, and (ii) the Median Symmetric Accuracy (Morley et al. 2018), which estimates the median percentage error between both annual indices *I*_*j*_:

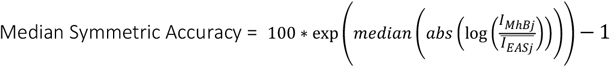

Note that the baseline year 2006 (both indices = 1) was excluded for both methods.

## 3 Results

Breeding bird trends were analysed for 61 out of 148 bird species (full detail in Appendix section 6, 7), but we focus the description of model assessment results on 14 exemplary species (tab. 1). Exemplified data for four species, script, and code are available from Github (https://github.com/m-rieger/EAS_bird) (Rieger et al. 2025) as well as a package with helper functions (https://github.com/m-rieger/EAS) (Rieger 2024).

### 3.1 Distribution characteristics: Species-specific optimisation

Model selection via k-fold cross validation resulted in different best-performing model families and a species-specific selection of coefficients added to the binomial part of zero-inflated models (Table 1, species-specific overview in Appendix section 7). We illustrate below the selection of model structure on four species that strongly vary in overall mean abundance, abundance dispersion, and the proportion of sites with zero counts: European Green Woodpecker, Eurasian Blue Tit, Black Redstart, and Eurasian Jay. The four measures that inform abundance distribution (propZ, mean, SD, max) can – but do not necessarily need to – vary with each other, so that an over- or underestimation of zero counts often goes along with an under- or overestimation of the population mean, and an over- or underestimation of SD with an over- or underestimation of maximum abundance.

**European Green Woodpecker** (Table 2, Fig. 3a) represents species with typically low territory numbers and many zero counts. While all models underestimated the number of zero counts, a Poisson model sufficiently captured other distribution parameters, resulting in a good predictive performance.

**Eurasian Blue Tit** (Table 2, Fig. 3b) is an abundant and widespread species that occurs in nearly all study sites. Here, abundance varied clearly more within sites and/or between year than reflected by a Poisson distribution. This issue of an underestimated SD was solved by using a negative binomial model family, even though leading to a slight overestimated SD.

**Black Redstart** (Table 2, Fig. 3c) is widespread at typically medium densities, resulting in a larger fraction of unoccupied sites than expected under a Poisson distribution. All models failed to capture this zero-bias, but models with a zero-inflation component were closer to the observed value. Since zero-inflated models mostly resulted in comparable ELPDs, we used the most parsimonious model (zero-inflated Poisson with no additional binomial coefficient) for further analyses.

**Eurasian Jay** (Table 2, Fig. 3d) is a widespread species with low territory numbers. Using zero-inflated models with environmental PCs as binomial model coefficient sufficiently captured the observed number of zero counts, and the zero-inflated negative binomial model resulted in the best predictive performance.

### 3.2 Spatial bias

For many species, (long-term) trends varied between natural regions, confirming the need to account for spatial bias. Especially long-term trends in metropolitan areas differed considerably from trends in other natural regions, usually showing steeper decreases in abundance per km^2^ (except for a steeper increase in Black Redstart, Fig. 4). Ignoring the overrepresentation of metropolitan sites (or any other unbalance in natural region samples) in the EAS dataset would therefore result in biased (long-term) trends for biogeographic regions and overall. In case of Eurasian Magpie and Eurasian Nuthatch, trends would be more negative and for Black Redstart more positive.

**Figure 4:**
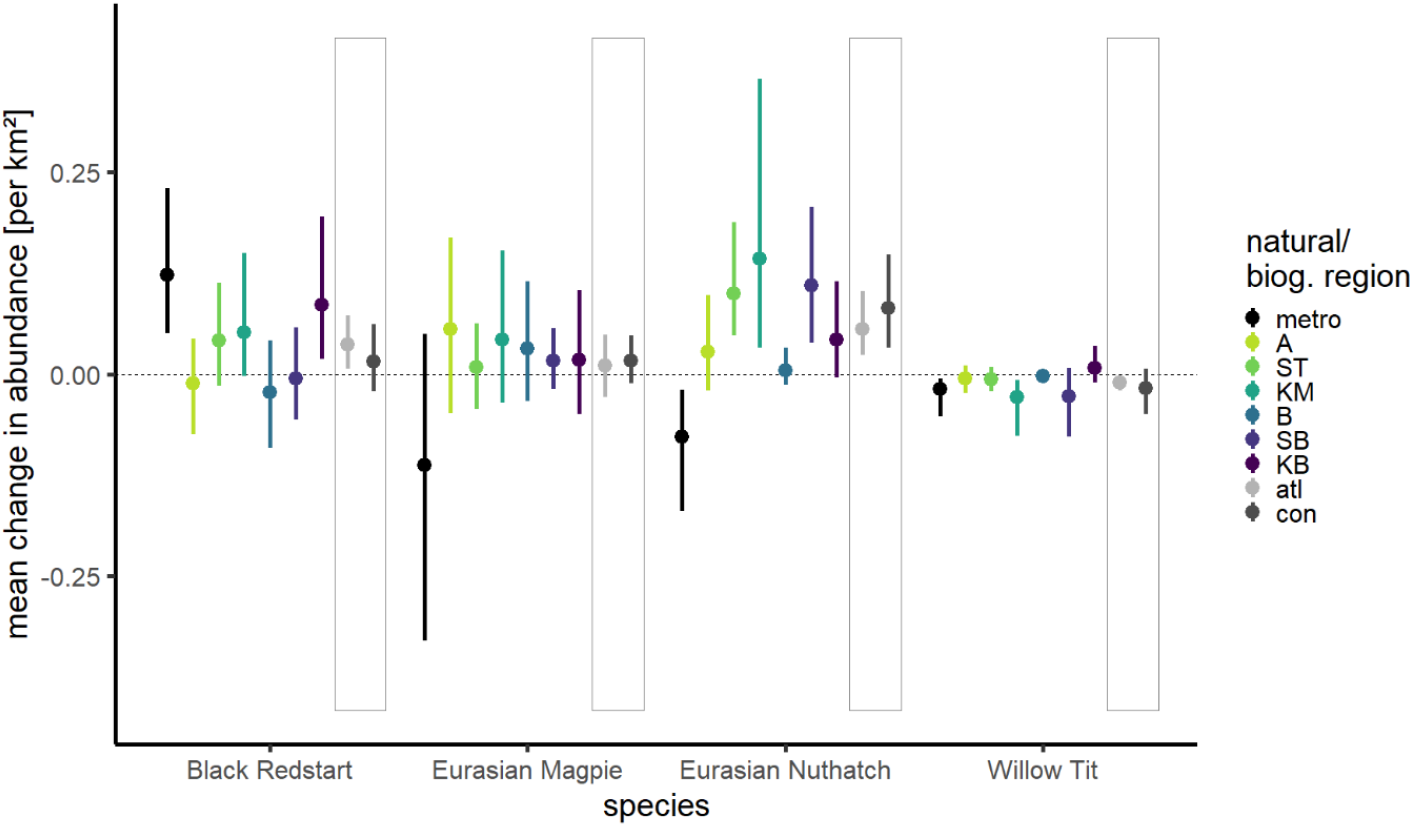
Mean annual change in abundance per km^2^ (long-term period 2008-2019) per natural region (See Fig. 1 for acronyms) and combined per overarching biogeographic region (atlantic - atl, and continental - con, grey boxes) for four example species.

### 3.3 Observer effects

We found a strikingly inhomogeneous distribution of observer effects across years, bearing substantial potential for biased population trend estimates when unaccounted for. High total abundances occurred disproportionally often since 2015, and strikingly low abundances before 2006 (Fig. 5, left), irrespective of the chosen threshold value (Appendix section 4). Models ignoring observer effects generally produced more positive long-term trend estimates, i.e., resulted in steeper increases or less steep declines than models with the covariate *observer effect* (Fig. 5, right). For Chaffinch, correcting for observer effects even reversed the increase in abundance in the non-corrected model to a decrease in the corrected model (Fig. 5, right). OE-corrected trend estimates were generally more consistent with West German trend estimates from the MhB programme (Appendix section 4) where consecutive surveys at a given site are always from the same observer.

**Figure 5:**
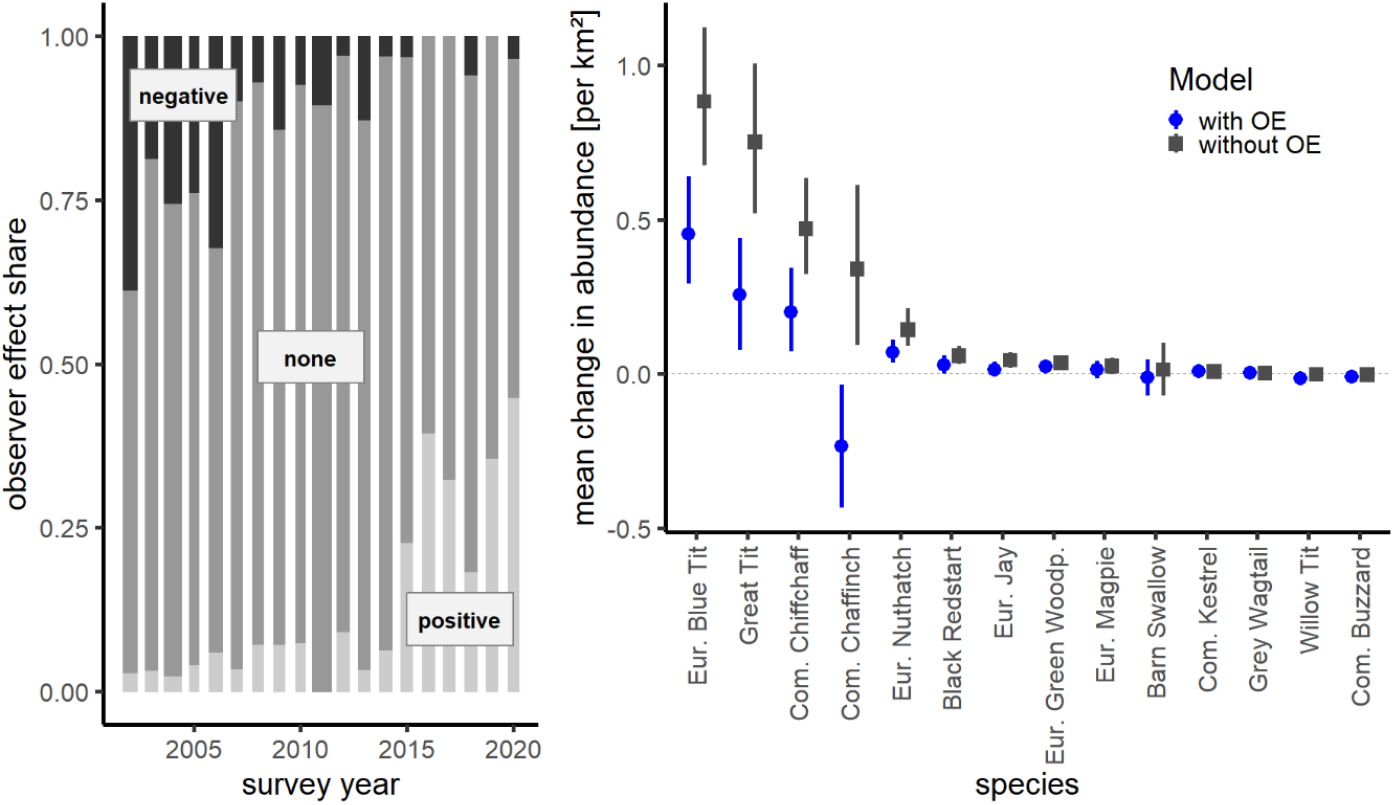
Left: Proportion of surveyed EAS sites per observer effect level per year. Bar widths are proportional to the number of surveyed sites per year. Right: Mean annual change in abundance per km^2^ (long-term period 2008-2019) estimated from models with and without the covariate observer effects for 14 species (example trend curves in Appendix section 4).

### 3.4 Consistency check

Correlation coefficients between our annual EAS indices and West German MhB indices revealed solid matches – i.e., with correlation coefficients and their 95% CI > 0 – for 34 out of 59 species (e.g., Nuthatch and Chiffchaff in Fig. 6a, b). Only five species (e.g., Eurasian Jay in Fig. 6d) tended to have negatively correlated yearly indices, but their 95% CI always included 0. Likewise, median symmetric accuracies (MSA) between index estimates were reasonably low, with 43 (or 26) out of 59 species deviating ≤ 20% (or ≤ 10%). Strong positive correlations do not necessarily go along with low MSA values, because the latter are rather sensitive to the chosen index baseline year as illustrated by the Chiffchaff data (Fig. 6b). Here, EAS- and MhB-analyses revealed rather consistent trend patterns for the period 2007–2020. Yet, the MSA-values were high, first because the exceptionally low MhB index in the baseline year 2006 shifted most other annual MhB-index values up so they exceeded the EAS values, but also because the smoothed trend did not reflect the magnitude of index differences between the early years 2005 + 2006 and later years. In contrast, Great Tit and Eurasian Jay exemplify how poor correlation between annual index values can go along with low MSA values (Fig. 6c, d), here because a rather stable vs. more fluctuating trend (Great Tit) that is identified by a weak correlation shows only minor deviations in trends (low MSA) or largely overlapping but slightly diverging trends (Eurasian Jay, especially due to year 2005) fail to correlate. Correlation coefficients, MSA values, and trend displays for all 59 species are in Appendix section 5.

**Figure 6:**
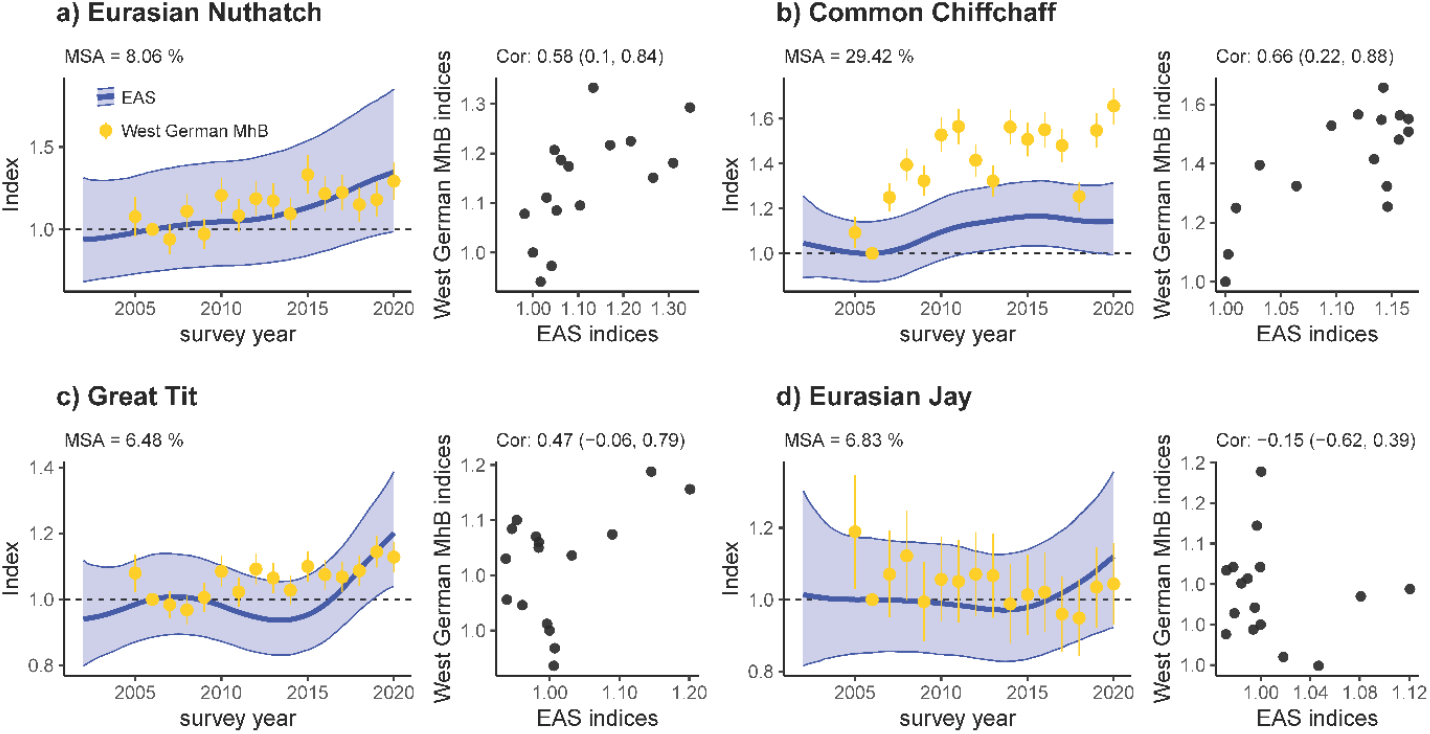
Trend comparison of EAS monitoring programme (this study, confidence intervals CI show uncertainty in the estimated index) and West German data from the German Common Bird Monitoring scheme (MhB, for comparison, CIs show uncertainty in index change) for four species. Left (per panel): Median Symmetric Accuracy (MSA) based on annual comparisons of indices. Right (per panel): correlation and correlation coefficients (± 95% CI) of annual indices from 2005 to 2020. Both approaches exclude the baseline year 2006.

## 4 Discussion

Count data are the raw unit for many population trend analyses, but their distribution poses several challenges during statistical analyses to account for overdispersion (42 out of 61 species in our current dataset) and/or zero inflation (45 out of 61 species). Moreover, many biodiversity monitoring programmes rest on ‘rolling’ sampling designs where each study site is visited only every so many years and observer identity frequently changes between years. We developed a workflow that takes these challenges into account (Fig. 7) (Rieger 2024, Rieger et al. 2025). First, we derive a routine that allows informed decisions about the most appropriate combination of residual families (Poisson or negative binomial), model covariates (e.g., habitat characteristics), and zero-inflation formulations to reflect species-specific data distributions. Second, we develop a correction factor that buffers population trend estimates for variation in observer expertise. Third, we integrate region-specific trends that adjust for between-year variation in the representation of habitat or natural regions within the yearly subset of sampled sites. Finally, by using generalised additive mixed models, we account for missing values due to rolling surveys. With these corrections in place, we find – within limits of the comparison – good consistency between the EAS trends and trends from the standard German Common Bird Monitoring scheme.

**Figure 7:**
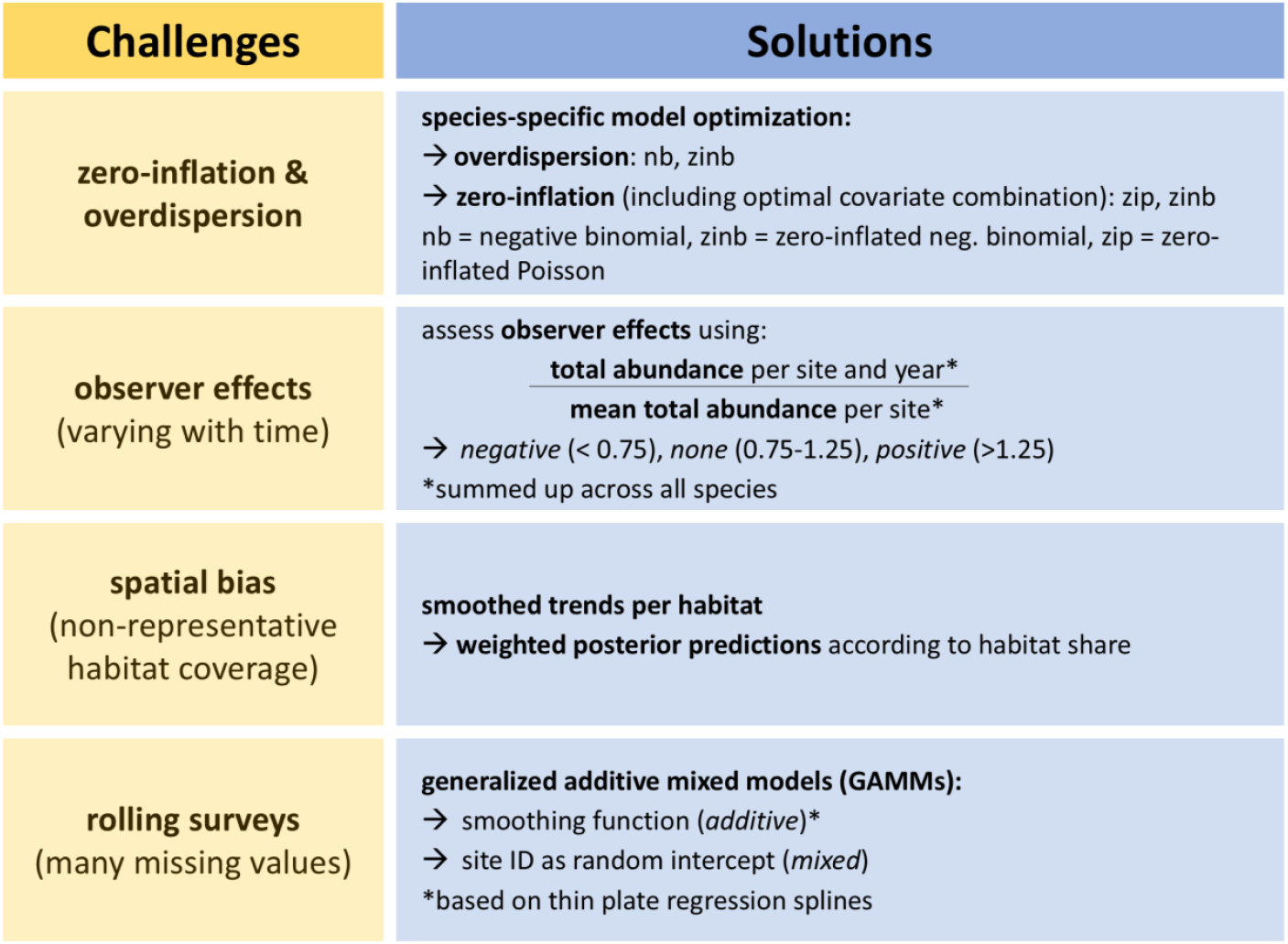
Workflow to address several challenges that arise when analysing breeding bird trends

### 4.1 Distribution characteristics: Species specific optimisation

We found substantial variation in model structures that best reflected the empirical abundance distributions and fraction of structural zeros across the 61 study species. Model formulations spanned from simple Poisson or negative binomial models (e.g., European Green Woodpecker, Eurasian Blue Tit; 26.2 % of all modelled species) to models with elaborate zero inflation formulae to account for variation in the fraction of structural zeros across natural regions, study sites, or environmental covariates (e.g., Great Spotted Woodpecker or Willow Tit; 73.8 % of all modelled species, tab. 1). This variation often tied in with breeding ecologies (cf. examples in section 3.1). Widespread and abundant species with intermediate territory counts in many study sites typically required rather simple count models but no correction for structural zeroes. In contrast, heterogeneously distributed species with a high fraction of unoccupied sites or colonial breeders with often large territory numbers at the few occupied sites almost exclusively required the integration of zero inflation formulae, possibly connected with controls for variation in zero frequencies between regions, sites or environments. This diversity in distribution and abundance patterns is typical for most (bird) monitoring programmes independent of the chosen modelling approach, with similar choices documented elsewhere (Etterson et al. 2009, Tirozzi et al. 2021, 2022, Hernández-Navarro et al. 2023).

### 4.2 Spatial bias

Modelling trends per natural region is a useful tool to reduce distorted trends due to spatial bias. This was exemplified by species such as Black Redstart, Eurasian Magpie, and Eurasian Nuthatch, showing striking diverging trends between metropolitan areas and other natural regions (Fig. 2, 4), which would otherwise bias regional or overall trends if discrepancies between natural region shares would be ignored. Similar distortions of trends or species distributions due to spatial bias often occur when using citizen science data since the sampled sites are selected by the observer and thus biased by its preferences (e.g., urban sites, sites of ecological interest (Johnston et al. 2020, Bowler et al. 2022))

Monitoring schemes should obviously aim to avoid the extreme disbalance in sampling that characterises the early phase of the EAS monitoring, where each year focused on a subset of natural regions. Such bias is difficult to correct *a posteriori* and bears the risk that compensating for missing values leads to a high level of uncertainty, masking possible trends. However, when the yearly sampling subsets are representative and include all present natural regions, as in the EAS programme since 2013 (Fig. 2), our approach can unfold its full capacity by accounting for minor bias in habitat coverage or the overrepresentation of metropolitan areas among study sites.

### 4.3 Observer effects

We found negative observer effects to occur disproportionally often during the early years of the EAS program but positive effects primarily in recent years. This likely arises from a combination of a temporal increase in expertise with ongoing involvement in the EAS program but also the new involvement of observers with individually higher population estimates during the later program years, with both effects visible in Fig. 4.3 of Appendix section 4.. This unfavourable combination of observer effects apparently caused substantial deviations in our abundance estimates compared to West German breeding bird trends, with a systematic shift towards more positive trends compared to MhB trends. In many cases, this bias was successfully mitigated by adding observer effect as a model covariate, now yielding trend estimates that were far more consistent with those of the West German Common Bird Monitoring programme (MhB, Appendix section 4).

Our approach to categorise observer effects from site-specific abundance data risks to mask drastic increases or declines in true bird abundance, e.g., after a substantial change in habitat quality or land-use. Such extreme shifts, however, will typically have diverging species-specific effects on abundance with winners and losers (Lemoine et al. 2007), and thus will massively affect total abundance across species just in exceptional extreme cases. Moreover, steady increases or declines in total abundance with habitat quality will also change mean total abundance across survey years, so that the threshold beyond which survey years are flagged as ‘extreme’ (positive or negative) also increases or decreases accordingly. Our categorisation fails, however, to identify cases of species-specific observer effects, such as insufficient expertise in discriminating ‘difficult’ species (e.g., songs of Garden Warbler vs. Blackcap) since total abundance is not affected by this, or hearing impairment for high pitched vocalisations (e.g., Goldcrest and Firecrest). For EAS, such cases are marked as implausible via an internal validating process and excluded from analysis (see section 2.5 ‘Data exclusion’).

Alternative indices may capture components of observer effort beyond our rather simplistic representation of expertise. A widespread approach integrates survey duration or transect length as model covariates, or combines both measures into survey duration per unit length (Kéry et al. 2005). Kéry et al. (2005) found higher survey effort to increase the detection rates of many bird species, resulting in higher abundance estimates. Others integrated observer age as a (quadratic) model covariate to correct for age-related changes in survey completeness (Farmer et al. 2014). Citizen Science data, e.g. those reported through online platforms such as ornitho (https://www.ornitho.de/) or ebird (https://ebird.org/), further allow estimating observer- and species-specific detection rates from species checklists, providing even more precise correction factors (Johnston et al. 2018). So far, the EAS monitoring programme cannot provide the data required to extract any of these covariates (i.e., effort, age, detection rates), but we highly recommend their integration to more precisely correct for possible observer effects in the future.

Some of the observer effects inherent to the EAS dataset likely arise from individual differences in the approach to delineate breeding territories. In future refinements of the program, this could be mitigated by using point observations per survey instead of territories as the basis for trend analyses, possibly in hierarchical models comparable to those used in other monitoring programmes such as the Swiss Breeding Bird survey (Kéry et al. 2005, Strebel et al. 2020). Alternatively, territory delineation could be automated from raw point observations, e.g., using the AutoTerri algorithm from TerriMap online developed by the Swiss Ornithological Institute (Wechsler 2018).

### 4.4 Consistency check

Population trend estimates for 59 species revealed a reasonably close match between the GAMM-based trend smoothers applied on the EAS dataset and TRIM-based yearly point estimates applied in the West German Common Bird Monitoring scheme. Despite substantial differences in the chosen modeling approach and spatial coverage, most trend estimates had solid positive correlations and/or a good Median Symmetric Accuracy between annual indices (Fig. 6, Appendix section 5). In many cases, smoothed GAMM-trends for the EAS data captured even short-term population fluctuations at a 3–7-year scale, similar to the – by definition – more fine-grained yearly MhB indices (e.g., Common Blackbird, Eurasian Wren, Firecrest, Appendix section 5). Neutral or even negative correlations between yearly EAS- and MhB-indices occurred primarily in two species subsets. The first concerns colonial birds (e.g., Common Swift, Eurasian Tree Sparrow, Common House Martin, Barn Swallow, Appendix section 5) where trend estimates are highly sensitive to the share of study sites with colonies present, the influence of few large colonies on the trend estimates, the precision of nest counts, and observer access to a given colony (Südbeck et al. 2005, LANUV 2016a). The second subset concerns species with pronounced affinity to urban areas as derived from low or high coefficient estimates for the linear effect of PC3 (Appendix section 6), where affinity can be positive (e.g., Eurasian Magpie, Dunnock, Common House Martin, Common Kestrel and Black Redstart) or negative (e.g., Common Chaffinch, Eurasian Jay, Marsh Tit or Crested Tit). Given an exceptionally high share and density of urban areas in NRW compared to other West German federal states, NRW trends are plausibly more positive than West German MhB trends for urban specialists (e.g., Eurasian Magpie, Dunnock, Common Kestrel, Black Redstart, Appendix section 5) and more negative for non-urban species (e.g. Common Chaffinch and Crested Tit, Appendix section 5).

## 5 Conclusion

We developed a modelling routine for bird population trend analyses that can handle several common problems in monitoring data. Our approach accounts for unbalanced sampling of natural regions, optimises species-specific model structure with respect to zero-inflation and overdispersion, partially accounts for variation in observer expertise, and integrates GAM-based smoothing to bridge even extensive gaps between replicated surveys in a ‘rolling’ sampling design. Our consistency check against trends from the Germany Common Bird Monitoring programme indicates that our routine produces reliable and robust trends for most species. The R script and package provided with this study can be adapted to other monitoring programmes with comparable survey designs and data structures.

## Acknowledgements

We thank the many observers for contributing to the EAS and MhB monitoring programmes, Matteo Santon for statistical and computational advice, and Jakob Katzenberger, Johannes Kamp, and Johannes

Wahl for comments on earlier versions of this workflow. We also thank Matthieu Paquet and two anonymous referees for their helpful feedback during the submission process.

## Data, scripts, code and supplementary information availability

A data example for four species, R script, and code are available from Github (https://github.com/m-rieger/EAS_bird) (Rieger et al. 2025) as well as a package with helper functions (https://github.com/m-rieger/EAS) (Rieger 2024). Further figures, model outputs and detailed descriptions are provided in the electronic supplementary material (‘EAS-bird_Appendix.html’).

