## Appendix for "Bird population trend analyses for a monitoring scheme with a highly structured sampling design": EASpaper_supplements.html


### 1 The ecological area sampling (EAS) dataset – additional information

The breeding bird territories per site (= abundance) derive from at least nine successive surveys per breeding season (LANUV 2016a). Each sighting of a bird is noted on a species-specific map using international atlas codes to indicate territorial, courtship or breeding behaviour. Based on these species-specific maps summed up across all surveys, the observer defines the number of breeding territories per site including peripheral territories which only count as .5 territories. A breeding territory is defined by the number and level of atlas codes which accumulate in a given area, as suggested by LANUV (2016a) and Südbeck et al. (2005). Observers record territories of all present bird species (from 2002-2020 148 species in total).

### 2 Site characterizations

#### 2.1 additional information

Several site-specific attributes were highly correlated, precluding the integration of all those variables as predictors in the statistical models, which assume predictor independence. Since these attributes are supposed to explain a lot of variation in abundance, we did a Principal Component Analysis (PCA) with these 14 numeric variables prior to the statistical analysis (tab. S2.1). Based on orthogonal linear transformation, a PCA summarizes the correlated variables in uncorrelated principal components (PC). Most variance of the entered dataset is depicted by the first PC of a PCA, whereas the subsequent orthogonal PCs depict as much of the left variance as possible (Crawley 2013). Therefore, only the first few PCs are considered for further analysis since they already explain a sufficient amount of variation (Crawley 2013). The amount of explained variation per entered covariate which is depicted by each PC is indicated by factor loadings.

Our PCA resulted in three meaningful PCs (eigenvalue > 1), with the first PC explaining 63% of the total variation, the second PC explaining 14%, and the third PC another 8%. To more clearly differentiate factor loadings (i.e., making PCs better interpretable with respect to the underlying environmental variables), we subjected the initial PCs an orthogonal rotation using the varimax-function as implemented in R (R Core Team 2020). As derived from factor loadings (tab. S2.1), the first rotated PC mainly covaried with climate data and altitude. PC1 scores increased with increasing altitude and precipitation and decreased with increasing temperatures and sunshine durations (tab. S2.1 PC1). Scores of the second rotated PC mainly increased with an increasing forest coverage and a decreasing arable land coverage and additionally covaried with increasing precipitation (tab. S2.1 PC2). PC3 scores mainly increased with an increasing settlement area coverage and decreased with an increasing arable land coverage (tab. S2.1 PC3).  

Table 2.1: Factor loadings after orthogonal rotation for the first three principal components of a PCA based on 14 site-specific attributes. Bold values indicate whether the site-specific attribute contributes to the meaning of the PC based on a threshold of ± 0.5.

| site-specific attributes | PC1 | PC2 | PC3 |
| --- | --- | --- | --- |
| altitude | 0.89 | 0.34 | -0.08 |
| mean temperature spring | -0.95 | -0.18 | 0.21 |
| mean temperature winter | -0.96 | -0.16 | 0.16 |
| minimum temperature spring | -0.93 | -0.09 | 0.24 |
| minimum temperature winter | -0.95 | -0.09 | 0.17 |
| maximum temperature spring | -0.93 | -0.2 | 0.17 |
| maximum temperature winter | -0.96 | -0.15 | 0.14 |
| precipitation spring | 0.63 | 0.6 | 0.1 |
| precipitation winter | 0.71 | 0.48 | 0.07 |
| sunshine duration spring | -0.69 | -0.38 | -0.26 |
| sunshine duration winter | -0.63 | 0.27 | -0.25 |
| arable land coverage | 0.06 | -0.76 | -0.49 |
| forest coverage | 0.23 | 0.81 | -0.31 |
| settlement area coverage | -0.2 | 0.02 | 0.93 |

#### 2.2 results

Coefficient estimates for the three principal components from a PCA on site-specific attributes were robust for many species (fig. S2.1, tab. S6.1). Additionally, they resulted in a better model fit when added as binomial model coefficient in 15% of all modelled species (tab. S6.1, tab. 1), underlining the importance of integrating environmental data into trend analyses.

Figure 2.1: Coefficient slope estimates (mean ± 95% CrI) for the three PCs (black = linear term, grey = quadratic term) from the respective final model for 14 species. Species are sorted by increasing linear PC1 estimates. If the 95% CrI does not include 0, the estimate is robust. Positive (negative) estimates of the linear term indicate a general increase (decrease) in abundance with increasing PC scores. Positive (negative) estimates of the quadratic term additionally indicate a u-formed (n-formed) relationship between abundance PC scores. PC1 scores increased with increasing altitude and precipitation and decreased with increasing temperatures and sunshine durations, PC2 scores increased with an increasing forest coverage and precipitation and decreases with increasing arable land coverage, PC3 scores increased with an increasing settlement area coverage and decreased with an increasing arable land coverage. For further details, see tab. S3.1 (factor loadings of PCs) & tab. S8.1 (coefficient estimates). Be aware that x-axes differ in scale and intercept.

Figure 2.2: Predicted mean abundance (including 95% CrI) per PC1, PC2 and PC3 score. PC1 scores increased with increasing altitude and precipitation and decreased with increasing temperatures and sunshine durations, PC2 scores increased with an increasing forest coverage and precipitation and decreases with increasing arable land coverage, PC3 scores increased with an increasing settlement area coverage and decreased with an increasing arable land coverage. For further details, see tab. S3.1 (factor loadings of PCs) & tab. S8.1 (coefficient estimates). Be aware that y-axes of different species differ in scale.

### 3 Distribution characteristics: Species-specific optimisation

Table 3.1: Examples for model assessment and the procedure to select final species-specific models from among 10 alternative model structures. Models differ in family and the presence of a zero-inflated model component (zinb and zip), the latter also in binomial model coefficients. Model assessment quantifies whether simulated data from the model is consistent with the observed response variable (abundance) with respect to its proportion of zeros (propZ), mean and maximum value, and standard deviation (SD). Bayesian p-values close to 0.5 indicate that the observed values are central within the posterior distribution of simulated raw datasets (8000 in total), indicating good model fit. Values close to 0 and 1 indicate substantial under- or overestimation and thus poor model fit. Final model selection is based on ELPD differences by selecting the most parsimonious model (bold, top to bottom) out of all models with an ELPD difference ± 1.96\*SE of ELPD difference including 0 (bold values). Due to convergence issues for Common Chaffinch and Great Tit with the most parsimonious model, the next best model was selected.

|  | | | | Bayesian p-value | | | | ELPD kfold | |
| --- | --- | --- | --- | --- | --- | --- | --- | --- | --- |
| Species |  | model family | binomial model coefficients | propZ | mean | SD | max | diff. | se diff. |
| Barn Swallow | Hirundo rustica | pois |  | 0.00 | 0.50 | 0.06 | 0.63 | -388.04 | 58.32 |
|  |  | nb |  | 0.00 | 0.93 | 1.00 | 1.00 | -53.85 | 15.66 |
|  |  | zip | ~ 1 | 0.03 | 0.04 | 0.04 | 0.64 | -257.79 | 45.68 |
|  |  | zinb | ~ 1 | 0.01 | 0.48 | 0.95 | 1.00 | -39.96 | 12.6 |
|  |  | zip | ~ PCs | 0.04 | 0.31 | 0.40 | 0.81 | -237.38 | 41.24 |
|  |  | zinb | ~ PCs | 0.02 | 0.68 | 0.98 | 1.00 | -28.74 | 13.3 |
|  |  | zip | ~ L | 0.02 | 0.08 | 0.10 | 0.68 | -260.74 | 44.02 |
|  |  | zinb | ~ L | 0.01 | 0.56 | 0.96 | 1.00 | -23.24 | 12.88 |
|  |  | zip | ~ (1\|ID) | 0.14 | 0.32 | 0.18 | 0.70 | -239.02 | 44.45 |
|  |  | zinb | ~ (1\|ID) | 0.12 | 0.63 | 0.93 | 0.99 | 0 | 0 |
| Black Redstart | Phoenicurus ochruros | pois |  | 0.00 | 0.51 | 0.56 | 0.47 | -23.37 | 10.14 |
|  |  | nb |  | 0.00 | 0.59 | 0.82 | 0.59 | -3654.6 | 780.81 |
|  |  | zip | ~ 1 | 0.00 | 0.30 | 0.48 | 0.45 | -8.51 | 9.42 |
|  |  | zinb | ~ 1 | 0.00 | 0.42 | 0.76 | 0.56 | -4.01 | 7.52 |
|  |  | zip | ~ PCs | 0.00 | 0.42 | 0.54 | 0.48 | -12.87 | 9.19 |
|  |  | zinb | ~ PCs | 0.00 | 0.51 | 0.79 | 0.57 | 0 | 0 |
|  |  | zip | ~ L | 0.00 | 0.34 | 0.50 | 0.46 | -18.86 | 9.43 |
|  |  | zinb | ~ L | 0.00 | 0.46 | 0.77 | 0.56 | -7.58 | 6.86 |
|  |  | zip | ~ (1\|ID) | 0.07 | 0.31 | 0.46 | 0.45 | -3.32 | 11.08 |
|  |  | zinb | ~ (1\|ID) | 0.08 | 0.40 | 0.69 | 0.52 | -2.98 | 8.83 |
| Common Buzzard | Buteo buteo | pois |  | 0.01 | 0.50 | 0.90 | 0.98 | -52.53 | 9.64 |
|  |  | zip | ~ 1 | 0.19 | 0.42 | 0.97 | 0.99 | -38.43 | 8.56 |
|  |  | zinb | ~ 1 | 0.18 | 0.45 | 0.97 | 0.99 | -39.51 | 8.55 |
|  |  | zip | ~ PCs | 0.19 | 0.46 | 0.97 | 0.99 | -44.67 | 9.21 |
|  |  | zinb | ~ PCs | 0.19 | 0.48 | 0.98 | 0.99 | -39.65 | 8.97 |
|  |  | zip | ~ L | 0.17 | 0.45 | 0.97 | 0.99 | -26.61 | 8.35 |
|  |  | zinb | ~ L | 0.18 | 0.47 | 0.98 | 1.00 | -27.23 | 8.26 |
|  |  | zip | ~ (1\|ID) | 0.47 | 0.48 | 0.97 | 0.95 | 0 | 0 |
|  |  | zinb | ~ (1\|ID) | 0.47 | 0.49 | 0.97 | 0.96 | -3.31 | 4.18 |
| Common Chaffinch | Fringilla coelebs | pois |  | 0.01 | 0.50 | 0.01 | 0.01 | -458.83 | 68.16 |
|  |  | nb |  | 0.02 | 0.65 | 0.90 | 0.55 | -4630933686 | 1481217405 |
|  |  | zip | ~ 1 | 0.61 | 0.36 | 0.01 | 0.01 | -447.2 | 70.26 |
|  |  | zinb | ~ 1 | 0.62 | 0.58 | 0.91 | 0.56 | -6.41 | 7.51 |
|  |  | zip | ~ PCs | 0.62 | 0.38 | 0.02 | 0.01 | -384.85 | 60.41 |
|  |  | zinb | ~ PCs | 0.63 | 0.57 | 0.90 | 0.55 | 0 | 0 |
|  |  | zip | ~ L | 0.61 | 0.36 | 0.01 | 0.01 | -428.04 | 60.84 |
|  |  | zinb | ~ L | 0.61 | 0.57 | 0.90 | 0.55 | -17.74 | 6.46 |
|  |  | zip | ~ (1\|ID) | 0.63 | 0.36 | 0.01 | 0.01 | -500.96 | 72.81 |
|  |  | zinb | ~ (1\|ID) | 0.62 | 0.57 | 0.90 | 0.56 | -4.37 | 7.18 |
| Common Chiffchaff | Phylloscopus collybita | pois |  | 0.00 | 0.49 | 0.03 | 0.25 | -194.75 | 31.45 |
|  |  | nb |  | 0.00 | 0.61 | 0.86 | 0.78 | -10.94 | 5.71 |
|  |  | zip | ~ 1 | 0.25 | 0.40 | 0.05 | 0.25 | -191.93 | 33.28 |
|  |  | zinb | ~ 1 | 0.23 | 0.54 | 0.87 | 0.79 | -24.14 | 6.9 |
|  |  | zip | ~ PCs | 0.26 | 0.40 | 0.05 | 0.24 | -200.61 | 33.21 |
|  |  | zinb | ~ PCs | 0.27 | 0.54 | 0.87 | 0.79 | 0 | 0 |
|  |  | zip | ~ L | 0.33 | 0.39 | 0.05 | 0.24 | -184.58 | 32.18 |
|  |  | zinb | ~ L | 0.31 | 0.54 | 0.87 | 0.79 | -18.1 | 5.57 |
|  |  | zip | ~ (1\|ID) | 0.25 | 0.40 | 0.05 | 0.25 | -189.33 | 32.2 |
|  |  | zinb | ~ (1\|ID) | 0.24 | 0.54 | 0.87 | 0.79 | -28.46 | 6.04 |
| Common Kestrel | Falco tinnunculus | pois |  | 0.00 | 0.49 | 0.62 | 1.00 | -63.19 | 10.82 |
|  |  | nb |  | 0.15 | 0.77 | 0.98 | 1.00 | -39.99 | 5.86 |
|  |  | zip | ~ 1 | 0.46 | 0.48 | 0.93 | 1.00 | -15.71 | 6.46 |
|  |  | zinb | ~ 1 | 0.46 | 0.50 | 0.93 | 1.00 | -17.52 | 6.37 |
|  |  | zip | ~ PCs | 0.43 | 0.49 | 0.92 | 1.00 | -17.99 | 6.36 |
|  |  | zinb | ~ PCs | 0.43 | 0.50 | 0.93 | 1.00 | -19.46 | 6.46 |
|  |  | zip | ~ L | 0.42 | 0.53 | 0.93 | 1.00 | -12.29 | 6.76 |
|  |  | zinb | ~ L | 0.42 | 0.52 | 0.94 | 1.00 | -17.72 | 6.8 |
|  |  | zip | ~ (1\|ID) | 0.45 | 0.47 | 0.92 | 1.00 | 0 | 0 |
|  |  | zinb | ~ (1\|ID) | 0.46 | 0.48 | 0.93 | 1.00 | -2.19 | 1.77 |
| Eurasian Blue Tit | Cyanistes caeruleus | pois |  | 0.00 | 0.50 | 0.01 | 0.02 | -510.17 | 69.13 |
|  |  | nb |  | 0.00 | 0.64 | 0.70 | 0.37 | -11.2 | 5.77 |
|  |  | zip | ~ 1 | 0.10 | 0.34 | 0.01 | 0.02 | -498.39 | 66.54 |
|  |  | zinb | ~ 1 | 0.07 | 0.58 | 0.69 | 0.37 | -19.9 | 8.09 |
|  |  | zip | ~ PCs | 0.11 | 0.39 | 0.01 | 0.02 | -509.75 | 71.86 |
|  |  | zinb | ~ PCs | 0.07 | 0.60 | 0.70 | 0.38 | -6.2 | 5.83 |
|  |  | zip | ~ L | 0.09 | 0.35 | 0.01 | 0.02 | -525.96 | 74.07 |
|  |  | zinb | ~ L | 0.06 | 0.59 | 0.70 | 0.38 | -3.48 | 6.89 |
|  |  | zip | ~ (1\|ID) | 0.11 | 0.33 | 0.01 | 0.02 | -499.92 | 71.67 |
|  |  | zinb | ~ (1\|ID) | 0.07 | 0.58 | 0.69 | 0.37 | 0 | 0 |
| Eurasian Jay | Garrulus glandarius | pois |  | 0.00 | 0.50 | 0.76 | 0.77 | -37.68 | 11.47 |
|  |  | nb |  | 0.00 | 0.69 | 0.89 | 0.70 | -117.44 | 22.38 |
|  |  | zip | ~ 1 | 0.00 | 0.31 | 0.78 | 0.76 | -27.5 | 8.37 |
|  |  | zinb | ~ 1 | 0.00 | 0.34 | 0.83 | 0.81 | -28.52 | 8.11 |
|  |  | zip | ~ PCs | 0.19 | 0.36 | 0.77 | 0.67 | -23.57 | 10.83 |
|  |  | zinb | ~ PCs | 0.19 | 0.39 | 0.81 | 0.72 | -15.08 | 10.25 |
|  |  | zip | ~ L | 0.01 | 0.29 | 0.76 | 0.75 | -36.03 | 8.93 |
|  |  | zinb | ~ L | 0.01 | 0.33 | 0.82 | 0.80 | -30.23 | 8.55 |
|  |  | zip | ~ (1\|ID) | 0.14 | 0.32 | 0.67 | 0.65 | -11.05 | 4.62 |
|  |  | zinb | ~ (1\|ID) | 0.14 | 0.34 | 0.74 | 0.71 | 0 | 0 |
| Eurasian Magpie | Pica pica | pois |  | 0.00 | 0.50 | 0.37 | 0.52 | -82.47 | 16.68 |
|  |  | nb |  | 0.25 | 0.78 | 0.90 | 0.90 | -37.17 | 8.73 |
|  |  | zip | ~ 1 | 0.00 | 0.34 | 0.31 | 0.50 | -92.15 | 17.84 |
|  |  | zinb | ~ 1 | 0.00 | 0.63 | 0.84 | 0.86 | -36.46 | 9.02 |
|  |  | zip | ~ PCs | 0.00 | 0.46 | 0.37 | 0.51 | -77.95 | 17.54 |
|  |  | zinb | ~ PCs | 0.00 | 0.65 | 0.83 | 0.84 | -33.61 | 9.22 |
|  |  | zip | ~ L | 0.00 | 0.36 | 0.35 | 0.50 | -89.69 | 17.5 |
|  |  | zinb | ~ L | 0.00 | 0.63 | 0.84 | 0.85 | -36.63 | 8.17 |
|  |  | zip | ~ (1\|ID) | 0.09 | 0.31 | 0.23 | 0.47 | -36.7 | 13.49 |
|  |  | zinb | ~ (1\|ID) | 0.13 | 0.43 | 0.63 | 0.77 | 0 | 0 |
| Eurasian Nuthatch | Sitta europaea | pois |  | 0.00 | 0.50 | 0.53 | 0.53 | -19.18 | 12.83 |
|  |  | nb |  | 0.00 | 0.59 | 0.79 | 0.69 | -5048.1 | 1228.04 |
|  |  | zip | ~ 1 | 0.00 | 0.37 | 0.49 | 0.53 | -32.99 | 12.56 |
|  |  | zinb | ~ 1 | 0.00 | 0.49 | 0.77 | 0.68 | -15.52 | 10.86 |
|  |  | zip | ~ PCs | 0.01 | 0.39 | 0.54 | 0.54 | -25.51 | 12.71 |
|  |  | zinb | ~ PCs | 0.01 | 0.49 | 0.77 | 0.67 | -20.14 | 10.96 |
|  |  | zip | ~ L | 0.00 | 0.34 | 0.47 | 0.52 | -18.19 | 11.48 |
|  |  | zinb | ~ L | 0.00 | 0.48 | 0.77 | 0.67 | -4.23 | 10.08 |
|  |  | zip | ~ (1\|ID) | 0.07 | 0.31 | 0.41 | 0.52 | -8.02 | 10.35 |
|  |  | zinb | ~ (1\|ID) | 0.07 | 0.37 | 0.64 | 0.64 | 0 | 0 |
| European Green Woodpecker | Picus viridis | pois |  | 0.01 | 0.49 | 0.79 | 0.65 | -11.86 | 7.89 |
|  |  | nb |  | 0.01 | 0.42 | 0.62 | 0.68 | -204.94 | 47.02 |
|  |  | zip | ~ 1 | 0.12 | 0.44 | 0.87 | 0.72 | -6.65 | 6.77 |
|  |  | zinb | ~ 1 | 0.13 | 0.46 | 0.91 | 0.80 | -8.06 | 6.73 |
|  |  | zip | ~ PCs | 0.16 | 0.46 | 0.88 | 0.71 | -3.66 | 6.92 |
|  |  | zinb | ~ PCs | 0.16 | 0.48 | 0.90 | 0.76 | -18.57 | 7.91 |
|  |  | zip | ~ L | 0.12 | 0.40 | 0.84 | 0.70 | -4.14 | 6.61 |
|  |  | zinb | ~ L | 0.12 | 0.45 | 0.88 | 0.77 | -4.76 | 6.49 |
|  |  | zip | ~ (1\|ID) | 0.28 | 0.40 | 0.78 | 0.55 | -0.21 | 3.49 |
|  |  | zinb | ~ (1\|ID) | 0.29 | 0.41 | 0.82 | 0.61 | 0 | 0 |
| Great Tit | Parus major | pois |  | 0.00 | 0.51 | 0.00 | 0.00 | -570.37 | 76.34 |
|  |  | nb |  | 0.00 | 0.64 | 0.65 | 0.39 | 0 | 0 |
|  |  | zip | ~ 1 | 0.43 | 0.29 | 0.00 | 0.00 | -587.05 | 74.03 |
|  |  | zinb | ~ 1 | 0.42 | 0.50 | 0.61 | 0.37 | -12.1 | 6.61 |
|  |  | zip | ~ PCs | 0.53 | 0.36 | 0.00 | 0.00 | -534.91 | 70.76 |
|  |  | zinb | ~ PCs | 0.52 | 0.55 | 0.61 | 0.36 | -2.84 | 6.81 |
|  |  | zip | ~ L | 0.45 | 0.31 | 0.00 | 0.01 | -531.65 | 71.08 |
|  |  | zinb | ~ L | 0.43 | 0.52 | 0.61 | 0.37 | -11.37 | 6.12 |
|  |  | zip | ~ (1\|ID) | 0.48 | 0.30 | 0.00 | 0.00 | -507.26 | 69.06 |
|  |  | zinb | ~ (1\|ID) | 0.45 | 0.50 | 0.61 | 0.37 | -10.51 | 6.86 |
| Grey Wagtail | Motacilla cinerea | pois |  | 0.06 | 0.50 | 0.78 | 0.56 | -13.28 | 5.46 |
|  |  | nb |  | 0.11 | 0.38 | 0.81 | 0.69 | -45.35 | 15.77 |
|  |  | zip | ~ 1 | 0.12 | 0.41 | 0.74 | 0.55 | -18.52 | 6.55 |
|  |  | zinb | ~ 1 | 0.13 | 0.46 | 0.79 | 0.62 | -19.31 | 6.47 |
|  |  | zip | ~ PCs | 0.15 | 0.50 | 0.82 | 0.64 | -3.91 | 5.04 |
|  |  | zinb | ~ PCs | 0.17 | 0.55 | 0.86 | 0.70 | -5.9 | 5.74 |
|  |  | zip | ~ L | 0.16 | 0.52 | 0.81 | 0.67 | -17.91 | 5.13 |
|  |  | zinb | ~ L | 0.19 | 0.53 | 0.84 | 0.71 | -18.11 | 4.99 |
|  |  | zip | ~ (1\|ID) | 0.37 | 0.47 | 0.75 | 0.45 | 0 | 0 |
|  |  | zinb | ~ (1\|ID) | 0.38 | 0.49 | 0.77 | 0.49 | -11.32 | 4.24 |
| Willow Tit | Poecile montanus | pois |  | 0.00 | 0.49 | 0.30 | 0.37 | -83.21 | 19.77 |
|  |  | nb |  | 0.00 | 0.82 | 0.99 | 0.99 | -33.13 | 10.74 |
|  |  | zip | ~ 1 | 0.07 | 0.25 | 0.54 | 0.45 | -16.03 | 9.31 |
|  |  | zinb | ~ 1 | 0.06 | 0.40 | 0.83 | 0.82 | -15.29 | 9.09 |
|  |  | zip | ~ PCs | 0.16 | 0.43 | 0.54 | 0.34 | -26.07 | 14.9 |
|  |  | zinb | ~ PCs | 0.07 | 0.62 | 0.89 | 0.88 | -6.95 | 12.37 |
|  |  | zip | ~ L | 0.15 | 0.22 | 0.46 | 0.41 | -6.6 | 10.28 |
|  |  | zinb | ~ L | 0.14 | 0.30 | 0.70 | 0.69 | -16.81 | 10.54 |
|  |  | zip | ~ (1\|ID) | 0.17 | 0.26 | 0.34 | 0.25 | -9.86 | 6.28 |
|  |  | zinb | ~ (1\|ID) | 0.17 | 0.31 | 0.56 | 0.53 | 0 | 0 |

### 4 Observer effects

Consistent with the expectation that striking deviation in total abundance across species goes along with similarly high (or low) abundance estimates for most individual species, we found that all three levels had different mean abundances in most species as indicated by posterior probabilities near 0 for ‘negative’ observer effects or 1 for ‘positive’ observer effects (tab. S6.1).

Visual interpretation of estimated trends for most species (Black Redstart, Common Buzzard, Common Chaffinch, Common Chiffchaff, Eurasian Blue Tit, Eurasian Jay, Eurasian Nuthatch, European Green Woodpecker, Great Tit, Willow Tit) confirmed that observer effects had a particularly strong impact on estimated abundance and thus population trends in the most recent years of the EAS program since about 2013, leading to a distinct separation of trend curves for those years (fig. S4.2).

Concerning the threshold of observer effects, a deviance of more than 30 %, 25 %, and 20 %, respectively resulted in similar estimated trends (fig. S4.2), although the number of ‘negative’ effects in earlier years and ‘positive’ effects in recent years increased with decreasing deviance threshold (fig. S4.1). We continued with the 25 % threshold.

Fig. S4.2 shows which observers are more prone to being classified as ‘positive’ or ‘negative’. Especially observers which were only once on a given site and had no previous experience in the programme (panel 1, yellow points) had deviating low or high total abundances, additionally showing the unfortunate pattern of more negative effects in the beginning and more positive effects in recent years. Familiarity with a site (panels 2 to 4) on average also resulted in higher abundances in consecutive surveys but these changes were usually less steep and within the 25 % threshold.

Figure 4.1: Proportion of surveyed EAS sites per observer effect level per year, comparing different thresholds to delineate negative and positive effects from neutral (= none): (a) ± 20 % deviation of the observed total abundance from the mean total abundance per site, b) ± 25 %, c) ± 30 %). Bar widths are proportional to the number of surveyed sites per year.

Figure 4.2: Estimated mean population indices (left) and estimated mean abundances (right) ± 95% CrI based on models without observer effects (OE) as covariate (grey), and models with observer effects as covariate for three different thresholds (25%, 20%, 15%, blue shades) for 14 species. For comparison, annual West German breeding bird indices from the MhB programme based on TRIM were added (mean ± 1.96\*SE, orange) in index plots. For population indices, the base year was 2006 with index = 1 (dashed horizontal line). Be aware that y-axes differ in scale and intercept.

Figure 4.3: Visualization of observer effects (ratio of total abundance per site and year and mean total abundance per site) across years. Points are colored by how many surveys within the EAS programme (irrespective of the site) an observer already did before the actual survey took place (a measure of experience within the programme) and lines connect surveys of the same site. The four panels depict, how many surveys of a given site were done by the observer (a measure of familiarity with the site), e.g. panel 1 shwos observers which surveyed a given site only once and panel 4 shows 4 different observers each doing 4 surveys on a given site. Their first surveys on the site are the leftmost points, their foruth surveys on the site the rightmost points. These repetitive surveys are connected with a line. Dashed lines display the thresholds to delineate observer effects as positive (above 1.25) or negative (below 0.75) and numbers give the share of observers per observer effect for a given panel.

### 5 Quality check

Figure 5.1: Trend comparison of EAS monitoring programme and West German MhB for 14 species. Left (per panel): Median Symmetric Accuracy (MSA) based on annual comparisons of indices. Right (per panel): correlation and correlation coefficients (± 95% CI) of annual indices from 2005 to 2020. Both approaches exclude the baseline year 2006. Be aware that axes differ in scale and intercept.

Figure 5.2: Left: Correlation coefficients of annual indices for EAS and West German MhB trends. Coefficients > 0 indicate a close fit between the two trend estimations, coefficients < 0 indicate diverging trend estimations. Right: Median symmetric accuracy between EAS and West German MhB trends. Values state the symmetric difference between trends in %.

### 6 Bayesian model coefficient estimates

Table 6.1: Mean Bayesian model coefficient estimates (*fit*) with 95% CrI (*lwr*, *upr*) for the final models of all 61 species including the posterior probability (*postP*) of the estimate to exceed 0. Posterior probabilities near 0 or 1 (bold values) indicate an increased explanatory power of the coefficient. Smooth and quadratic coefficient estimates indicate whether the smoothed or quadratic trend line differs from a linear regression. An estimate near 0 does not differ from a linear regression. Coefficient levels for observer effects were *none*, *negative*, and *positive*, for natural regions *A*, *B*, *KB*, *KM*, *metro*, *SB*, and *ST*. The column ‘model’ indicates whether the coefficient was derived from a Poisson (*pois*), a negative binomial (*nb*), or – in case of a zero-inflated distribution – from the binomial model (*binomial (zi)*).

|  | | | | coefficient estimate | | |  |
| --- | --- | --- | --- | --- | --- | --- | --- |
| species | scientific | model | coefficient | fit | lwr | upr | postP |
| Eurasian Nuthatch | Sitta europaea | pois | intercept [negative, A] | 0.278 | -0.394 | 0.984 | 0.797 |
|  |  |  | region [B] | -1.035 | -1.825 | -0.246 | 0.006 |
|  |  |  | region [KB] | -0.217 | -1.178 | 0.738 | 0.331 |
|  |  |  | region [KM] | 0.455 | -0.454 | 1.345 | 0.844 |
|  |  |  | region [metro] | 0.060 | -0.779 | 0.877 | 0.557 |
|  |  |  | region [SB] | -0.349 | -1.301 | 0.590 | 0.244 |
|  |  |  | region [ST] | 0.414 | -0.359 | 1.156 | 0.856 |
|  |  |  | observer effect [none] | 0.439 | 0.292 | 0.587 | 1 |
|  |  |  | observer effect [positive] | 0.864 | 0.704 | 1.026 | 1 |
|  |  |  | PC1 - linear | 9.265 | 1.727 | 16.783 | 0.993 |
|  |  |  | PC2 - linear | 16.529 | 10.416 | 22.444 | 1 |
|  |  |  | PC3 - linear | -15.953 | -20.839 | -11.069 | 0 |
|  |  |  | PC1 - quadratic | -5.914 | -10.608 | -1.285 | 0.008 |
|  |  |  | PC2 - quadratic | -1.202 | -6.295 | 3.737 | 0.319 |
|  |  |  | PC3 - quadratic | -7.141 | -12.196 | -1.814 | 0.004 |
|  |  |  | survey year:region [A] | 0.286 | -2.358 | 2.037 | 0.647 |
|  |  |  | survey year:region [B] | 0.321 | -2.924 | 3.847 | 0.624 |
|  |  |  | survey year:region [KB] | 0.673 | -1.872 | 2.956 | 0.775 |
|  |  |  | survey year:region [KM] | 0.938 | -2.482 | 4.338 | 0.742 |
|  |  |  | survey year:region [metro] | -0.797 | -3.505 | 2.751 | 0.283 |
|  |  |  | survey year:region [SB] | 0.846 | -0.993 | 2.827 | 0.905 |
|  |  |  | survey year:region [ST] | 1.008 | -0.773 | 2.726 | 0.928 |
|  |  |  | survey year:region [A] - smoother | 0.314 | 0.013 | 1.797 | 1 |
|  |  |  | survey year:region [B] - smoother | 0.656 | 0.026 | 3.594 | 1 |
|  |  |  | survey year:region [KB] - smoother | 0.362 | 0.014 | 1.873 | 1 |
|  |  |  | survey year:region [KM] - smoother | 0.918 | 0.154 | 3.319 | 1 |
|  |  |  | survey year:region [metro] - smoother | 0.630 | 0.051 | 2.557 | 1 |
|  |  |  | survey year:region [SB] - smoother | 0.287 | 0.011 | 1.725 | 1 |
|  |  |  | survey year:region [ST] - smoother | 0.274 | 0.012 | 1.597 | 1 |
|  |  |  | ID intercept | 1.113 | 0.971 | 1.283 | 1 |
| European Green Woodpecker | Picus viridis | pois | intercept [negative, A] | -1.081 | -1.899 | -0.324 | 0.002 |
|  |  |  | region [B] | -0.490 | -1.291 | 0.316 | 0.106 |
|  |  |  | region [KB] | -1.062 | -2.136 | -0.045 | 0.02 |
|  |  |  | region [KM] | -0.236 | -1.112 | 0.686 | 0.302 |
|  |  |  | region [metro] | -0.352 | -1.188 | 0.526 | 0.204 |
|  |  |  | region [SB] | -1.407 | -2.387 | -0.373 | 0.004 |
|  |  |  | region [ST] | -0.799 | -1.571 | -0.045 | 0.021 |
|  |  |  | observer effect [none] | 0.363 | -0.022 | 0.777 | 0.965 |
|  |  |  | observer effect [positive] | 0.917 | 0.500 | 1.356 | 1 |
|  |  |  | PC1 - linear | -11.621 | -20.910 | -2.553 | 0.006 |
|  |  |  | PC2 - linear | 10.886 | 4.008 | 17.432 | 0.999 |
|  |  |  | PC3 - linear | 0.584 | -4.878 | 6.002 | 0.578 |
|  |  |  | PC1 - quadratic | -9.666 | -16.599 | -3.305 | 0.002 |
|  |  |  | PC2 - quadratic | -3.151 | -8.544 | 2.296 | 0.127 |
|  |  |  | PC3 - quadratic | -7.791 | -13.884 | -1.833 | 0.005 |
|  |  |  | survey year:region [A] | 0.746 | -2.531 | 4.809 | 0.695 |
|  |  |  | survey year:region [B] | 1.213 | -3.445 | 5.532 | 0.708 |
|  |  |  | survey year:region [KB] | 1.081 | -2.366 | 4.437 | 0.76 |
|  |  |  | survey year:region [KM] | 0.800 | -3.468 | 4.999 | 0.658 |
|  |  |  | survey year:region [metro] | 0.491 | -2.737 | 3.675 | 0.642 |
|  |  |  | survey year:region [SB] | 2.084 | -1.389 | 4.797 | 0.909 |
|  |  |  | survey year:region [ST] | 1.472 | -1.254 | 4.218 | 0.897 |
|  |  |  | survey year:region [A] - smoother | 1.164 | 0.039 | 5.211 | 1 |
|  |  |  | survey year:region [B] - smoother | 2.264 | 0.639 | 6.785 | 1 |
|  |  |  | survey year:region [KB] - smoother | 0.678 | 0.024 | 3.603 | 1 |
|  |  |  | survey year:region [KM] - smoother | 1.743 | 0.245 | 5.439 | 1 |
|  |  |  | survey year:region [metro] - smoother | 0.625 | 0.031 | 2.824 | 1 |
|  |  |  | survey year:region [SB] - smoother | 0.553 | 0.024 | 2.972 | 1 |
|  |  |  | survey year:region [ST] - smoother | 0.513 | 0.021 | 2.679 | 1 |
|  |  |  | ID intercept | 1.038 | 0.850 | 1.274 | 1 |
| Stock Dove | Columba oenas | pois | intercept [negative, A] | -1.639 | -2.672 | -0.636 | 0.001 |
|  |  |  | region [B] | -0.753 | -1.991 | 0.412 | 0.107 |
|  |  |  | region [KB] | -1.665 | -3.260 | -0.159 | 0.017 |
|  |  |  | region [KM] | -0.912 | -2.309 | 0.470 | 0.098 |
|  |  |  | region [metro] | -0.487 | -1.780 | 0.722 | 0.215 |
|  |  |  | region [SB] | -1.095 | -2.470 | 0.246 | 0.062 |
|  |  |  | region [ST] | 0.459 | -0.642 | 1.545 | 0.794 |
|  |  |  | observer effect [none] | 0.312 | 0.021 | 0.616 | 0.982 |
|  |  |  | observer effect [positive] | 0.806 | 0.451 | 1.164 | 1 |
|  |  |  | PC1 - linear | -7.023 | -18.098 | 4.387 | 0.114 |
|  |  |  | PC2 - linear | 0.452 | -9.159 | 10.135 | 0.538 |
|  |  |  | PC3 - linear | -21.603 | -30.214 | -13.330 | 0 |
|  |  |  | PC1 - quadratic | 7.805 | 0.145 | 15.100 | 0.977 |
|  |  |  | PC2 - quadratic | -1.396 | -9.912 | 7.344 | 0.374 |
|  |  |  | PC3 - quadratic | -1.482 | -10.373 | 7.287 | 0.369 |
|  |  |  | survey year:region [A] | 0.863 | -1.664 | 3.767 | 0.802 |
|  |  |  | survey year:region [B] | 0.776 | -4.009 | 5.609 | 0.615 |
|  |  |  | survey year:region [KB] | 1.172 | -2.733 | 4.429 | 0.756 |
|  |  |  | survey year:region [KM] | 0.431 | -3.262 | 3.621 | 0.606 |
|  |  |  | survey year:region [metro] | 0.884 | -2.972 | 4.243 | 0.712 |
|  |  |  | survey year:region [SB] | 0.277 | -4.634 | 4.969 | 0.541 |
|  |  |  | survey year:region [ST] | 0.059 | -3.514 | 2.850 | 0.523 |
|  |  |  | survey year:region [A] - smoother | 0.486 | 0.027 | 2.582 | 1 |
|  |  |  | survey year:region [B] - smoother | 4.777 | 1.884 | 10.831 | 1 |
|  |  |  | survey year:region [KB] - smoother | 0.898 | 0.051 | 3.889 | 1 |
|  |  |  | survey year:region [KM] - smoother | 0.688 | 0.033 | 3.160 | 1 |
|  |  |  | survey year:region [metro] - smoother | 0.859 | 0.034 | 3.843 | 1 |
|  |  |  | survey year:region [SB] - smoother | 4.814 | 1.966 | 11.708 | 1 |
|  |  |  | survey year:region [ST] - smoother | 0.832 | 0.076 | 5.325 | 1 |
|  |  |  | ID intercept | 1.692 | 1.403 | 2.059 | 1 |
| Coal Tit | Periparus ater | nb | intercept [negative, A] | -2.009 | -3.107 | -1.004 | 0 |
|  |  |  | region [B] | -0.855 | -2.217 | 0.480 | 0.103 |
|  |  |  | region [KB] | 1.109 | -0.220 | 2.525 | 0.947 |
|  |  |  | region [KM] | 0.837 | -0.444 | 2.223 | 0.89 |
|  |  |  | region [metro] | -0.351 | -1.693 | 0.933 | 0.292 |
|  |  |  | region [SB] | 1.448 | 0.135 | 2.847 | 0.985 |
|  |  |  | region [ST] | 1.426 | 0.297 | 2.557 | 0.995 |
|  |  |  | observer effect [none] | 0.326 | 0.106 | 0.541 | 0.999 |
|  |  |  | observer effect [positive] | 0.769 | 0.502 | 1.032 | 1 |
|  |  |  | PC1 - linear | 34.802 | 24.157 | 45.150 | 1 |
|  |  |  | PC2 - linear | 20.135 | 11.555 | 28.569 | 1 |
|  |  |  | PC3 - linear | -17.911 | -24.549 | -11.171 | 0 |
|  |  |  | PC1 - quadratic | -9.138 | -15.904 | -2.360 | 0.005 |
|  |  |  | PC2 - quadratic | -4.224 | -11.505 | 3.075 | 0.135 |
|  |  |  | PC3 - quadratic | 0.480 | -6.409 | 7.386 | 0.553 |
|  |  |  | survey year:region [A] | -1.196 | -4.159 | 3.204 | 0.273 |
|  |  |  | survey year:region [B] | -1.742 | -5.402 | 2.925 | 0.224 |
|  |  |  | survey year:region [KB] | -0.713 | -3.571 | 2.499 | 0.288 |
|  |  |  | survey year:region [KM] | -0.829 | -4.570 | 2.855 | 0.297 |
|  |  |  | survey year:region [metro] | 0.157 | -3.107 | 3.636 | 0.547 |
|  |  |  | survey year:region [SB] | -1.137 | -3.943 | 1.028 | 0.092 |
|  |  |  | survey year:region [ST] | -0.040 | -2.035 | 3.745 | 0.487 |
|  |  |  | survey year:region [A] - smoother | 0.865 | 0.039 | 3.773 | 1 |
|  |  |  | survey year:region [B] - smoother | 1.067 | 0.051 | 4.548 | 1 |
|  |  |  | survey year:region [KB] - smoother | 0.619 | 0.036 | 2.684 | 1 |
|  |  |  | survey year:region [KM] - smoother | 0.891 | 0.042 | 4.014 | 1 |
|  |  |  | survey year:region [metro] - smoother | 0.618 | 0.029 | 3.217 | 1 |
|  |  |  | survey year:region [SB] - smoother | 0.454 | 0.019 | 2.527 | 1 |
|  |  |  | survey year:region [ST] - smoother | 0.645 | 0.028 | 2.797 | 1 |
|  |  |  | ID intercept | 1.487 | 1.246 | 1.798 | 1 |
| Common Chiffchaff | Phylloscopus collybita | nb | intercept [negative, A] | 2.709 | 2.425 | 3.016 | 1 |
|  |  |  | region [B] | -1.024 | -1.375 | -0.679 | 0 |
|  |  |  | region [KB] | -0.602 | -1.023 | -0.183 | 0.002 |
|  |  |  | region [KM] | -0.117 | -0.507 | 0.260 | 0.27 |
|  |  |  | region [metro] | -0.324 | -0.660 | 0.024 | 0.035 |
|  |  |  | region [SB] | -0.530 | -0.965 | -0.133 | 0.005 |
|  |  |  | region [ST] | -0.025 | -0.350 | 0.277 | 0.436 |
|  |  |  | observer effect [none] | 0.525 | 0.432 | 0.616 | 1 |
|  |  |  | observer effect [positive] | 0.773 | 0.668 | 0.879 | 1 |
|  |  |  | PC1 - linear | 0.819 | -2.595 | 4.373 | 0.676 |
|  |  |  | PC2 - linear | 4.827 | 2.217 | 7.557 | 1 |
|  |  |  | PC3 - linear | -1.103 | -3.175 | 0.930 | 0.141 |
|  |  |  | PC1 - quadratic | -3.885 | -5.972 | -1.781 | 0 |
|  |  |  | PC2 - quadratic | -2.638 | -4.860 | -0.333 | 0.012 |
|  |  |  | PC3 - quadratic | 0.254 | -1.942 | 2.432 | 0.59 |
|  |  |  | survey year:region [A] | 0.150 | -1.878 | 1.961 | 0.615 |
|  |  |  | survey year:region [B] | 1.133 | -2.576 | 4.531 | 0.753 |
|  |  |  | survey year:region [KB] | -0.208 | -4.384 | 1.996 | 0.452 |
|  |  |  | survey year:region [KM] | 0.434 | -2.109 | 2.562 | 0.707 |
|  |  |  | survey year:region [metro] | -0.879 | -3.157 | 0.806 | 0.099 |
|  |  |  | survey year:region [SB] | -0.075 | -2.897 | 1.768 | 0.464 |
|  |  |  | survey year:region [ST] | -0.134 | -1.921 | 0.848 | 0.363 |
|  |  |  | survey year:region [A] - smoother | 0.255 | 0.009 | 1.679 | 1 |
|  |  |  | survey year:region [B] - smoother | 0.963 | 0.347 | 3.501 | 1 |
|  |  |  | survey year:region [KB] - smoother | 0.759 | 0.031 | 2.866 | 1 |
|  |  |  | survey year:region [KM] - smoother | 0.339 | 0.016 | 1.827 | 1 |
|  |  |  | survey year:region [metro] - smoother | 0.298 | 0.015 | 1.531 | 1 |
|  |  |  | survey year:region [SB] - smoother | 0.425 | 0.013 | 2.632 | 1 |
|  |  |  | survey year:region [ST] - smoother | 0.216 | 0.009 | 1.351 | 1 |
|  |  |  | ID intercept | 0.480 | 0.426 | 0.542 | 1 |
| Common Pheasant | Phasianus colchicus | nb | intercept [negative, A] | -0.966 | -1.581 | -0.354 | 0.002 |
|  |  |  | region [B] | 0.057 | -0.544 | 0.640 | 0.58 |
|  |  |  | region [KB] | -1.232 | -2.149 | -0.384 | 0.002 |
|  |  |  | region [KM] | 0.488 | -0.218 | 1.197 | 0.911 |
|  |  |  | region [metro] | -0.618 | -1.534 | 0.150 | 0.06 |
|  |  |  | region [SB] | -1.948 | -2.908 | -1.083 | 0 |
|  |  |  | region [ST] | 0.207 | -0.378 | 0.790 | 0.761 |
|  |  |  | observer effect [none] | -0.151 | -0.406 | 0.095 | 0.112 |
|  |  |  | observer effect [positive] | 0.338 | 0.046 | 0.634 | 0.987 |
|  |  |  | PC1 - linear | -39.584 | -51.756 | -27.671 | 0 |
|  |  |  | PC2 - linear | -32.827 | -40.571 | -25.062 | 0 |
|  |  |  | PC3 - linear | 5.069 | -1.030 | 11.048 | 0.948 |
|  |  |  | PC1 - quadratic | -10.465 | -18.389 | -2.340 | 0.005 |
|  |  |  | PC2 - quadratic | -17.594 | -23.885 | -11.658 | 0 |
|  |  |  | PC3 - quadratic | -12.057 | -19.959 | -4.843 | 0.001 |
|  |  |  | survey year:region [A] | -0.971 | -3.558 | 2.425 | 0.231 |
|  |  |  | survey year:region [B] | -0.607 | -3.230 | 3.474 | 0.341 |
|  |  |  | survey year:region [KB] | 0.563 | -2.757 | 4.220 | 0.654 |
|  |  |  | survey year:region [KM] | -0.812 | -3.411 | 2.641 | 0.264 |
|  |  |  | survey year:region [metro] | 0.075 | -3.203 | 4.048 | 0.516 |
|  |  |  | survey year:region [SB] | -0.145 | -3.546 | 3.518 | 0.463 |
|  |  |  | survey year:region [ST] | 0.227 | -3.905 | 3.900 | 0.549 |
|  |  |  | survey year:region [A] - smoother | 0.561 | 0.026 | 2.934 | 1 |
|  |  |  | survey year:region [B] - smoother | 0.675 | 0.034 | 3.179 | 1 |
|  |  |  | survey year:region [KB] - smoother | 0.784 | 0.032 | 4.319 | 1 |
|  |  |  | survey year:region [KM] - smoother | 0.509 | 0.028 | 2.554 | 1 |
|  |  |  | survey year:region [metro] - smoother | 0.779 | 0.034 | 4.271 | 1 |
|  |  |  | survey year:region [SB] - smoother | 0.714 | 0.030 | 3.774 | 1 |
|  |  |  | survey year:region [ST] - smoother | 1.856 | 0.615 | 5.036 | 1 |
|  |  |  | ID intercept | 0.788 | 0.634 | 0.972 | 1 |
| Common Starling | Sturnus vulgaris | nb | intercept [negative, A] | 0.528 | -0.220 | 1.246 | 0.921 |
|  |  |  | region [B] | -0.097 | -0.893 | 0.746 | 0.412 |
|  |  |  | region [KB] | -0.027 | -1.035 | 0.976 | 0.477 |
|  |  |  | region [KM] | -0.195 | -1.130 | 0.755 | 0.347 |
|  |  |  | region [metro] | -0.456 | -1.321 | 0.442 | 0.154 |
|  |  |  | region [SB] | 0.385 | -0.560 | 1.411 | 0.784 |
|  |  |  | region [ST] | 0.533 | -0.244 | 1.336 | 0.913 |
|  |  |  | observer effect [none] | 0.432 | 0.205 | 0.651 | 1 |
|  |  |  | observer effect [positive] | 0.796 | 0.535 | 1.053 | 1 |
|  |  |  | PC1 - linear | -18.223 | -26.453 | -10.137 | 0 |
|  |  |  | PC2 - linear | -3.063 | -9.230 | 3.317 | 0.163 |
|  |  |  | PC3 - linear | 11.409 | 6.404 | 16.472 | 1 |
|  |  |  | PC1 - quadratic | -1.290 | -6.581 | 4.082 | 0.312 |
|  |  |  | PC2 - quadratic | -5.146 | -10.267 | 0.062 | 0.026 |
|  |  |  | PC3 - quadratic | -5.321 | -10.817 | 0.161 | 0.028 |
|  |  |  | survey year:region [A] | -0.013 | -2.696 | 2.609 | 0.495 |
|  |  |  | survey year:region [B] | -0.068 | -2.883 | 2.796 | 0.472 |
|  |  |  | survey year:region [KB] | -0.757 | -4.205 | 2.970 | 0.297 |
|  |  |  | survey year:region [KM] | -0.236 | -3.406 | 3.207 | 0.426 |
|  |  |  | survey year:region [metro] | -1.443 | -5.105 | 1.877 | 0.166 |
|  |  |  | survey year:region [SB] | -0.648 | -3.591 | 1.366 | 0.195 |
|  |  |  | survey year:region [ST] | -0.141 | -2.801 | 2.421 | 0.443 |
|  |  |  | survey year:region [A] - smoother | 0.412 | 0.018 | 2.323 | 1 |
|  |  |  | survey year:region [B] - smoother | 0.428 | 0.017 | 2.385 | 1 |
|  |  |  | survey year:region [KB] - smoother | 0.831 | 0.033 | 4.395 | 1 |
|  |  |  | survey year:region [KM] - smoother | 0.688 | 0.038 | 2.973 | 1 |
|  |  |  | survey year:region [metro] - smoother | 0.941 | 0.048 | 4.079 | 1 |
|  |  |  | survey year:region [SB] - smoother | 0.384 | 0.014 | 2.364 | 1 |
|  |  |  | survey year:region [ST] - smoother | 0.671 | 0.126 | 2.407 | 1 |
|  |  |  | ID intercept | 1.170 | 1.022 | 1.350 | 1 |
| Common Wood Pigeon | Columba palumbus | nb | intercept [negative, A] | 2.161 | 1.835 | 2.497 | 1 |
|  |  |  | region [B] | -0.145 | -0.505 | 0.202 | 0.204 |
|  |  |  | region [KB] | -0.157 | -0.616 | 0.304 | 0.257 |
|  |  |  | region [KM] | 0.228 | -0.194 | 0.631 | 0.855 |
|  |  |  | region [metro] | -0.238 | -0.619 | 0.164 | 0.116 |
|  |  |  | region [SB] | -0.356 | -0.845 | 0.108 | 0.066 |
|  |  |  | region [ST] | 0.368 | 0.011 | 0.717 | 0.979 |
|  |  |  | observer effect [none] | 0.503 | 0.375 | 0.631 | 1 |
|  |  |  | observer effect [positive] | 0.921 | 0.775 | 1.070 | 1 |
|  |  |  | PC1 - linear | -6.235 | -10.120 | -2.350 | 0.001 |
|  |  |  | PC2 - linear | 8.426 | 5.611 | 11.245 | 1 |
|  |  |  | PC3 - linear | 3.760 | 1.521 | 6.061 | 0.999 |
|  |  |  | PC1 - quadratic | 0.067 | -2.233 | 2.371 | 0.524 |
|  |  |  | PC2 - quadratic | -4.401 | -6.734 | -2.043 | 0 |
|  |  |  | PC3 - quadratic | -2.365 | -4.691 | -0.004 | 0.025 |
|  |  |  | survey year:region [A] | 0.311 | -1.599 | 3.220 | 0.673 |
|  |  |  | survey year:region [B] | 0.310 | -2.685 | 2.526 | 0.637 |
|  |  |  | survey year:region [KB] | -0.398 | -3.017 | 2.234 | 0.325 |
|  |  |  | survey year:region [KM] | 0.026 | -2.131 | 3.238 | 0.514 |
|  |  |  | survey year:region [metro] | 0.251 | -2.764 | 2.534 | 0.612 |
|  |  |  | survey year:region [SB] | 0.809 | -0.664 | 2.613 | 0.914 |
|  |  |  | survey year:region [ST] | 0.771 | -1.206 | 3.192 | 0.848 |
|  |  |  | survey year:region [A] - smoother | 0.379 | 0.015 | 2.033 | 1 |
|  |  |  | survey year:region [B] - smoother | 0.400 | 0.022 | 1.954 | 1 |
|  |  |  | survey year:region [KB] - smoother | 0.465 | 0.026 | 2.121 | 1 |
|  |  |  | survey year:region [KM] - smoother | 0.408 | 0.016 | 2.126 | 1 |
|  |  |  | survey year:region [metro] - smoother | 0.399 | 0.017 | 2.597 | 1 |
|  |  |  | survey year:region [SB] - smoother | 0.198 | 0.009 | 1.202 | 1 |
|  |  |  | survey year:region [ST] - smoother | 0.529 | 0.030 | 2.069 | 1 |
|  |  |  | ID intercept | 0.500 | 0.439 | 0.571 | 1 |
| Eurasian Blackcap | Sylvia atricapilla | nb | intercept [negative, A] | 2.728 | 2.424 | 3.006 | 1 |
|  |  |  | region [B] | -0.635 | -0.952 | -0.303 | 0 |
|  |  |  | region [KB] | -0.326 | -0.732 | 0.088 | 0.058 |
|  |  |  | region [KM] | 0.001 | -0.364 | 0.382 | 0.501 |
|  |  |  | region [metro] | 0.055 | -0.269 | 0.389 | 0.628 |
|  |  |  | region [SB] | -0.430 | -0.837 | -0.011 | 0.021 |
|  |  |  | region [ST] | -0.167 | -0.471 | 0.156 | 0.151 |
|  |  |  | observer effect [none] | 0.403 | 0.311 | 0.496 | 1 |
|  |  |  | observer effect [positive] | 0.639 | 0.532 | 0.745 | 1 |
|  |  |  | PC1 - linear | 3.519 | 0.095 | 6.915 | 0.978 |
|  |  |  | PC2 - linear | 7.353 | 4.962 | 9.894 | 1 |
|  |  |  | PC3 - linear | -2.853 | -4.848 | -0.892 | 0.003 |
|  |  |  | PC1 - quadratic | -2.182 | -4.176 | -0.103 | 0.019 |
|  |  |  | PC2 - quadratic | -2.266 | -4.316 | -0.204 | 0.015 |
|  |  |  | PC3 - quadratic | -1.377 | -3.465 | 0.774 | 0.102 |
|  |  |  | survey year:region [A] | 1.163 | -0.981 | 2.918 | 0.906 |
|  |  |  | survey year:region [B] | 1.748 | -1.706 | 4.858 | 0.893 |
|  |  |  | survey year:region [KB] | 1.265 | -0.797 | 2.852 | 0.927 |
|  |  |  | survey year:region [KM] | 0.724 | -1.988 | 2.810 | 0.78 |
|  |  |  | survey year:region [metro] | 0.508 | -1.028 | 2.646 | 0.826 |
|  |  |  | survey year:region [SB] | 1.443 | -0.877 | 3.177 | 0.926 |
|  |  |  | survey year:region [ST] | 1.100 | -0.498 | 2.328 | 0.943 |
|  |  |  | survey year:region [A] - smoother | 0.263 | 0.010 | 1.751 | 1 |
|  |  |  | survey year:region [B] - smoother | 0.705 | 0.038 | 2.749 | 1 |
|  |  |  | survey year:region [KB] - smoother | 0.218 | 0.008 | 1.432 | 1 |
|  |  |  | survey year:region [KM] - smoother | 0.333 | 0.014 | 1.689 | 1 |
|  |  |  | survey year:region [metro] - smoother | 0.211 | 0.010 | 1.232 | 1 |
|  |  |  | survey year:region [SB] - smoother | 0.348 | 0.012 | 1.713 | 1 |
|  |  |  | survey year:region [ST] - smoother | 0.248 | 0.016 | 1.235 | 1 |
|  |  |  | ID intercept | 0.455 | 0.404 | 0.512 | 1 |
| Eurasian Blue Tit | Cyanistes caeruleus | nb | intercept [negative, A] | 2.211 | 1.853 | 2.569 | 1 |
|  |  |  | region [B] | -0.414 | -0.795 | -0.038 | 0.017 |
|  |  |  | region [KB] | -0.202 | -0.665 | 0.287 | 0.204 |
|  |  |  | region [KM] | 0.261 | -0.173 | 0.698 | 0.877 |
|  |  |  | region [metro] | 0.029 | -0.386 | 0.439 | 0.552 |
|  |  |  | region [SB] | -0.055 | -0.532 | 0.443 | 0.412 |
|  |  |  | region [ST] | 0.036 | -0.323 | 0.419 | 0.578 |
|  |  |  | observer effect [none] | 0.485 | 0.366 | 0.607 | 1 |
|  |  |  | observer effect [positive] | 0.910 | 0.771 | 1.047 | 1 |
|  |  |  | PC1 - linear | -5.826 | -9.933 | -1.755 | 0.001 |
|  |  |  | PC2 - linear | 11.894 | 8.957 | 14.949 | 1 |
|  |  |  | PC3 - linear | 3.668 | 1.280 | 6.003 | 0.998 |
|  |  |  | PC1 - quadratic | -5.542 | -8.040 | -3.035 | 0 |
|  |  |  | PC2 - quadratic | -3.438 | -6.015 | -0.857 | 0.004 |
|  |  |  | PC3 - quadratic | -1.912 | -4.522 | 0.616 | 0.071 |
|  |  |  | survey year:region [A] | 0.824 | -1.912 | 2.739 | 0.81 |
|  |  |  | survey year:region [B] | 0.470 | -2.236 | 2.924 | 0.723 |
|  |  |  | survey year:region [KB] | 0.966 | -1.992 | 3.496 | 0.815 |
|  |  |  | survey year:region [KM] | 0.374 | -2.519 | 2.656 | 0.677 |
|  |  |  | survey year:region [metro] | 0.458 | -1.611 | 4.196 | 0.679 |
|  |  |  | survey year:region [SB] | 1.034 | -1.540 | 2.629 | 0.878 |
|  |  |  | survey year:region [ST] | 0.852 | -0.684 | 2.845 | 0.923 |
|  |  |  | survey year:region [A] - smoother | 0.337 | 0.015 | 1.802 | 1 |
|  |  |  | survey year:region [B] - smoother | 0.367 | 0.017 | 1.870 | 1 |
|  |  |  | survey year:region [KB] - smoother | 0.451 | 0.017 | 3.355 | 1 |
|  |  |  | survey year:region [KM] - smoother | 0.351 | 0.018 | 1.954 | 1 |
|  |  |  | survey year:region [metro] - smoother | 0.524 | 0.028 | 2.535 | 1 |
|  |  |  | survey year:region [SB] - smoother | 0.281 | 0.010 | 1.828 | 1 |
|  |  |  | survey year:region [ST] - smoother | 0.277 | 0.011 | 1.653 | 1 |
|  |  |  | ID intercept | 0.544 | 0.479 | 0.620 | 1 |
| Eurasian Wren | Troglodytes troglodytes | nb | intercept [negative, A] | 2.408 | 2.039 | 2.797 | 1 |
|  |  |  | region [B] | -0.750 | -1.164 | -0.314 | 0 |
|  |  |  | region [KB] | -0.568 | -1.093 | -0.054 | 0.015 |
|  |  |  | region [KM] | -0.028 | -0.501 | 0.457 | 0.455 |
|  |  |  | region [metro] | 0.202 | -0.239 | 0.641 | 0.814 |
|  |  |  | region [SB] | -0.576 | -1.101 | -0.059 | 0.014 |
|  |  |  | region [ST] | 0.147 | -0.253 | 0.558 | 0.76 |
|  |  |  | observer effect [none] | 0.475 | 0.354 | 0.592 | 1 |
|  |  |  | observer effect [positive] | 0.806 | 0.673 | 0.936 | 1 |
|  |  |  | PC1 - linear | 4.021 | -0.338 | 8.383 | 0.966 |
|  |  |  | PC2 - linear | 10.594 | 7.261 | 13.891 | 1 |
|  |  |  | PC3 - linear | -3.871 | -6.481 | -1.277 | 0.002 |
|  |  |  | PC1 - quadratic | -2.465 | -5.105 | 0.195 | 0.035 |
|  |  |  | PC2 - quadratic | -4.088 | -6.828 | -1.304 | 0.004 |
|  |  |  | PC3 - quadratic | -0.102 | -2.816 | 2.762 | 0.473 |
|  |  |  | survey year:region [A] | -0.987 | -5.033 | 3.167 | 0.304 |
|  |  |  | survey year:region [B] | 0.467 | -3.156 | 3.496 | 0.621 |
|  |  |  | survey year:region [KB] | 0.131 | -3.752 | 2.767 | 0.532 |
|  |  |  | survey year:region [KM] | -0.178 | -4.092 | 3.144 | 0.458 |
|  |  |  | survey year:region [metro] | -0.488 | -4.081 | 3.573 | 0.402 |
|  |  |  | survey year:region [SB] | 0.054 | -3.200 | 2.601 | 0.52 |
|  |  |  | survey year:region [ST] | 0.149 | -3.662 | 3.519 | 0.535 |
|  |  |  | survey year:region [A] - smoother | 2.316 | 0.450 | 5.180 | 1 |
|  |  |  | survey year:region [B] - smoother | 0.835 | 0.222 | 2.663 | 1 |
|  |  |  | survey year:region [KB] - smoother | 0.867 | 0.073 | 2.735 | 1 |
|  |  |  | survey year:region [KM] - smoother | 1.148 | 0.227 | 3.654 | 1 |
|  |  |  | survey year:region [metro] - smoother | 1.623 | 0.596 | 4.328 | 1 |
|  |  |  | survey year:region [SB] - smoother | 0.821 | 0.029 | 3.108 | 1 |
|  |  |  | survey year:region [ST] - smoother | 1.920 | 0.712 | 4.458 | 1 |
|  |  |  | ID intercept | 0.605 | 0.536 | 0.686 | 1 |
| Song Thrush | Turdus philomelos | nb | intercept [negative, A] | 1.544 | 1.173 | 1.907 | 1 |
|  |  |  | region [B] | -0.571 | -0.986 | -0.165 | 0.004 |
|  |  |  | region [KB] | -0.163 | -0.685 | 0.338 | 0.264 |
|  |  |  | region [KM] | 0.220 | -0.244 | 0.687 | 0.821 |
|  |  |  | region [metro] | 0.219 | -0.209 | 0.653 | 0.845 |
|  |  |  | region [SB] | -0.103 | -0.626 | 0.402 | 0.349 |
|  |  |  | region [ST] | 0.155 | -0.240 | 0.545 | 0.787 |
|  |  |  | observer effect [none] | 0.310 | 0.184 | 0.435 | 1 |
|  |  |  | observer effect [positive] | 0.616 | 0.470 | 0.769 | 1 |
|  |  |  | PC1 - linear | 7.001 | 2.797 | 11.295 | 0.999 |
|  |  |  | PC2 - linear | 4.912 | 1.751 | 8.156 | 0.998 |
|  |  |  | PC3 - linear | -4.914 | -7.429 | -2.343 | 0 |
|  |  |  | PC1 - quadratic | -3.105 | -5.669 | -0.584 | 0.007 |
|  |  |  | PC2 - quadratic | -3.044 | -5.720 | -0.414 | 0.013 |
|  |  |  | PC3 - quadratic | -0.958 | -3.582 | 1.749 | 0.251 |
|  |  |  | survey year:region [A] | 0.263 | -2.421 | 2.354 | 0.635 |
|  |  |  | survey year:region [B] | 0.306 | -3.127 | 2.923 | 0.615 |
|  |  |  | survey year:region [KB] | 0.346 | -1.931 | 3.357 | 0.673 |
|  |  |  | survey year:region [KM] | -0.172 | -2.523 | 2.718 | 0.423 |
|  |  |  | survey year:region [metro] | -0.123 | -2.016 | 2.682 | 0.437 |
|  |  |  | survey year:region [SB] | -0.006 | -2.038 | 1.156 | 0.494 |
|  |  |  | survey year:region [ST] | 0.391 | -1.238 | 2.766 | 0.763 |
|  |  |  | survey year:region [A] - smoother | 0.347 | 0.013 | 2.308 | 1 |
|  |  |  | survey year:region [B] - smoother | 0.510 | 0.023 | 2.377 | 1 |
|  |  |  | survey year:region [KB] - smoother | 0.434 | 0.018 | 2.235 | 1 |
|  |  |  | survey year:region [KM] - smoother | 0.410 | 0.020 | 2.197 | 1 |
|  |  |  | survey year:region [metro] - smoother | 0.318 | 0.014 | 1.829 | 1 |
|  |  |  | survey year:region [SB] - smoother | 0.185 | 0.009 | 1.224 | 1 |
|  |  |  | survey year:region [ST] - smoother | 0.380 | 0.016 | 1.954 | 1 |
|  |  |  | ID intercept | 0.568 | 0.498 | 0.652 | 1 |
| Tree Pipit | Anthus trivialis | nb | intercept [negative, A] | -4.218 | -5.839 | -2.710 | 0 |
|  |  |  | region [B] | -0.702 | -2.748 | 1.184 | 0.235 |
|  |  |  | region [KB] | 1.461 | -0.393 | 3.361 | 0.938 |
|  |  |  | region [KM] | 0.780 | -1.213 | 2.773 | 0.777 |
|  |  |  | region [metro] | -3.155 | -6.659 | -0.273 | 0.013 |
|  |  |  | region [SB] | 1.399 | -0.388 | 3.301 | 0.937 |
|  |  |  | region [ST] | 2.540 | 0.920 | 4.164 | 0.999 |
|  |  |  | observer effect [none] | 0.052 | -0.514 | 0.631 | 0.574 |
|  |  |  | observer effect [positive] | 0.589 | -0.128 | 1.342 | 0.943 |
|  |  |  | PC1 - linear | 18.930 | 4.491 | 32.415 | 0.995 |
|  |  |  | PC2 - linear | -3.958 | -16.819 | 8.848 | 0.262 |
|  |  |  | PC3 - linear | -23.176 | -34.998 | -11.911 | 0 |
|  |  |  | PC1 - quadratic | -3.968 | -14.073 | 6.174 | 0.218 |
|  |  |  | PC2 - quadratic | -2.155 | -13.808 | 8.653 | 0.353 |
|  |  |  | PC3 - quadratic | -3.740 | -15.311 | 7.339 | 0.261 |
|  |  |  | survey year:region [A] | -1.145 | -5.134 | 3.262 | 0.299 |
|  |  |  | survey year:region [B] | 0.163 | -3.910 | 4.219 | 0.534 |
|  |  |  | survey year:region [KB] | 1.078 | -2.486 | 4.508 | 0.756 |
|  |  |  | survey year:region [KM] | -1.098 | -4.923 | 3.195 | 0.298 |
|  |  |  | survey year:region [metro] | -0.245 | -5.154 | 4.633 | 0.46 |
|  |  |  | survey year:region [SB] | 0.144 | -3.431 | 3.228 | 0.544 |
|  |  |  | survey year:region [ST] | -1.094 | -4.656 | 1.933 | 0.212 |
|  |  |  | survey year:region [A] - smoother | 1.003 | 0.055 | 4.373 | 1 |
|  |  |  | survey year:region [B] - smoother | 0.890 | 0.036 | 4.319 | 1 |
|  |  |  | survey year:region [KB] - smoother | 0.724 | 0.032 | 3.733 | 1 |
|  |  |  | survey year:region [KM] - smoother | 0.917 | 0.038 | 4.577 | 1 |
|  |  |  | survey year:region [metro] - smoother | 1.538 | 0.059 | 7.066 | 1 |
|  |  |  | survey year:region [SB] - smoother | 0.745 | 0.037 | 3.308 | 1 |
|  |  |  | survey year:region [ST] - smoother | 0.774 | 0.035 | 3.852 | 1 |
|  |  |  | ID intercept | 2.213 | 1.779 | 2.759 | 1 |
| Willow Warbler | Phylloscopus trochilus | nb | intercept [negative, A] | 0.886 | 0.160 | 1.568 | 0.992 |
|  |  |  | region [B] | -1.226 | -2.037 | -0.422 | 0.002 |
|  |  |  | region [KB] | -1.043 | -1.997 | -0.056 | 0.019 |
|  |  |  | region [KM] | -0.492 | -1.400 | 0.475 | 0.156 |
|  |  |  | region [metro] | -0.281 | -1.110 | 0.559 | 0.264 |
|  |  |  | region [SB] | -0.843 | -1.805 | 0.173 | 0.05 |
|  |  |  | region [ST] | 0.552 | -0.162 | 1.304 | 0.934 |
|  |  |  | observer effect [none] | 0.345 | 0.103 | 0.578 | 0.997 |
|  |  |  | observer effect [positive] | 0.492 | 0.194 | 0.780 | 1 |
|  |  |  | PC1 - linear | 19.287 | 10.799 | 27.309 | 1 |
|  |  |  | PC2 - linear | 7.306 | 0.871 | 13.548 | 0.987 |
|  |  |  | PC3 - linear | -4.460 | -9.471 | 0.521 | 0.039 |
|  |  |  | PC1 - quadratic | -5.996 | -11.021 | -0.966 | 0.008 |
|  |  |  | PC2 - quadratic | -4.094 | -9.233 | 0.980 | 0.062 |
|  |  |  | PC3 - quadratic | 0.368 | -4.925 | 5.646 | 0.555 |
|  |  |  | survey year:region [A] | -1.127 | -4.102 | 1.908 | 0.176 |
|  |  |  | survey year:region [B] | 0.212 | -3.505 | 3.441 | 0.561 |
|  |  |  | survey year:region [KB] | 0.195 | -2.562 | 3.409 | 0.571 |
|  |  |  | survey year:region [KM] | -1.718 | -5.377 | 2.764 | 0.218 |
|  |  |  | survey year:region [metro] | -1.834 | -4.856 | 2.756 | 0.198 |
|  |  |  | survey year:region [SB] | -0.160 | -2.824 | 2.218 | 0.419 |
|  |  |  | survey year:region [ST] | -1.421 | -4.450 | 0.938 | 0.074 |
|  |  |  | survey year:region [A] - smoother | 0.571 | 0.024 | 3.198 | 1 |
|  |  |  | survey year:region [B] - smoother | 0.742 | 0.032 | 3.679 | 1 |
|  |  |  | survey year:region [KB] - smoother | 0.513 | 0.022 | 2.569 | 1 |
|  |  |  | survey year:region [KM] - smoother | 1.380 | 0.127 | 4.785 | 1 |
|  |  |  | survey year:region [metro] - smoother | 0.849 | 0.044 | 3.433 | 1 |
|  |  |  | survey year:region [SB] - smoother | 0.474 | 0.028 | 2.119 | 1 |
|  |  |  | survey year:region [ST] - smoother | 0.527 | 0.022 | 2.755 | 1 |
|  |  |  | ID intercept | 1.149 | 1.002 | 1.323 | 1 |
| Yellowhammer | Emberiza citrinella | nb | intercept [negative, A] | 0.011 | -0.654 | 0.654 | 0.514 |
|  |  |  | region [B] | 0.318 | -0.361 | 1.032 | 0.818 |
|  |  |  | region [KB] | 0.576 | -0.336 | 1.470 | 0.905 |
|  |  |  | region [KM] | 0.629 | -0.193 | 1.438 | 0.931 |
|  |  |  | region [metro] | -0.427 | -1.339 | 0.428 | 0.168 |
|  |  |  | region [SB] | 0.760 | -0.137 | 1.686 | 0.952 |
|  |  |  | region [ST] | 0.367 | -0.337 | 1.084 | 0.849 |
|  |  |  | observer effect [none] | 0.285 | 0.113 | 0.454 | 0.999 |
|  |  |  | observer effect [positive] | 0.395 | 0.193 | 0.599 | 1 |
|  |  |  | PC1 - linear | 2.416 | -5.057 | 9.995 | 0.735 |
|  |  |  | PC2 - linear | -20.786 | -26.922 | -15.038 | 0 |
|  |  |  | PC3 - linear | 1.943 | -3.012 | 6.936 | 0.778 |
|  |  |  | PC1 - quadratic | -12.875 | -17.983 | -8.052 | 0 |
|  |  |  | PC2 - quadratic | -5.571 | -10.821 | -0.437 | 0.016 |
|  |  |  | PC3 - quadratic | -9.997 | -15.502 | -4.551 | 0 |
|  |  |  | survey year:region [A] | 0.436 | -2.544 | 3.224 | 0.66 |
|  |  |  | survey year:region [B] | -0.891 | -3.462 | 2.357 | 0.226 |
|  |  |  | survey year:region [KB] | -0.098 | -2.127 | 2.195 | 0.438 |
|  |  |  | survey year:region [KM] | -0.763 | -3.855 | 2.126 | 0.224 |
|  |  |  | survey year:region [metro] | -1.452 | -4.627 | 2.953 | 0.233 |
|  |  |  | survey year:region [SB] | -0.145 | -3.616 | 1.544 | 0.415 |
|  |  |  | survey year:region [ST] | -3.104 | -6.537 | 0.411 | 0.039 |
|  |  |  | survey year:region [A] - smoother | 0.546 | 0.026 | 2.779 | 1 |
|  |  |  | survey year:region [B] - smoother | 0.495 | 0.030 | 2.229 | 1 |
|  |  |  | survey year:region [KB] - smoother | 0.321 | 0.015 | 1.743 | 1 |
|  |  |  | survey year:region [KM] - smoother | 0.514 | 0.023 | 2.741 | 1 |
|  |  |  | survey year:region [metro] - smoother | 0.817 | 0.032 | 3.827 | 1 |
|  |  |  | survey year:region [SB] - smoother | 0.371 | 0.014 | 2.487 | 1 |
|  |  |  | survey year:region [ST] - smoother | 1.419 | 0.384 | 4.329 | 1 |
|  |  |  | ID intercept | 1.036 | 0.900 | 1.205 | 1 |
| Black Redstart | Phoenicurus ochruros | pois | intercept [negative, A] | 0.285 | -0.273 | 0.839 | 0.839 |
|  |  |  | region [B] | 0.317 | -0.325 | 0.977 | 0.826 |
|  |  |  | region [KB] | 0.008 | -0.796 | 0.804 | 0.508 |
|  |  |  | region [KM] | -0.178 | -0.964 | 0.586 | 0.323 |
|  |  |  | region [metro] | -0.563 | -1.220 | 0.116 | 0.052 |
|  |  |  | region [SB] | 0.423 | -0.375 | 1.207 | 0.854 |
|  |  |  | region [ST] | 0.257 | -0.353 | 0.865 | 0.792 |
|  |  |  | observer effect [none] | 0.210 | 0.087 | 0.335 | 1 |
|  |  |  | observer effect [positive] | 0.508 | 0.366 | 0.647 | 1 |
|  |  |  | PC1 - linear | -5.448 | -11.909 | 1.045 | 0.047 |
|  |  |  | PC2 - linear | 6.319 | 1.445 | 11.452 | 0.994 |
|  |  |  | PC3 - linear | 26.026 | 21.580 | 30.788 | 1 |
|  |  |  | PC1 - quadratic | -3.075 | -7.603 | 1.259 | 0.083 |
|  |  |  | PC2 - quadratic | -4.475 | -8.727 | -0.384 | 0.018 |
|  |  |  | PC3 - quadratic | -9.464 | -14.233 | -5.063 | 0 |
|  |  |  | survey year:region [A] | -0.079 | -2.140 | 2.820 | 0.457 |
|  |  |  | survey year:region [B] | -0.104 | -2.932 | 2.661 | 0.449 |
|  |  |  | survey year:region [KB] | 1.027 | -2.291 | 3.229 | 0.795 |
|  |  |  | survey year:region [KM] | 0.692 | -2.695 | 3.556 | 0.72 |
|  |  |  | survey year:region [metro] | 1.146 | -1.238 | 3.261 | 0.901 |
|  |  |  | survey year:region [SB] | -1.065 | -4.595 | 1.210 | 0.197 |
|  |  |  | survey year:region [ST] | -0.104 | -2.125 | 1.883 | 0.44 |
|  |  |  | survey year:region [A] - smoother | 0.335 | 0.012 | 2.123 | 1 |
|  |  |  | survey year:region [B] - smoother | 0.482 | 0.033 | 2.435 | 1 |
|  |  |  | survey year:region [KB] - smoother | 0.438 | 0.021 | 2.157 | 1 |
|  |  |  | survey year:region [KM] - smoother | 0.582 | 0.022 | 3.339 | 1 |
|  |  |  | survey year:region [metro] - smoother | 0.271 | 0.012 | 1.578 | 1 |
|  |  |  | survey year:region [SB] - smoother | 0.778 | 0.224 | 2.628 | 1 |
|  |  |  | survey year:region [ST] - smoother | 0.457 | 0.069 | 1.811 | 1 |
|  |  |  | ID intercept | 0.922 | 0.803 | 1.067 | 1 |
|  |  | binomial (zi) | zi - intercept | -3.841 | -5.249 | -3.036 | 0 |
| Common Swift | Apus apus | pois | intercept [negative, A] | -3.823 | -5.721 | -2.185 | 0 |
|  |  |  | region [B] | 0.398 | -1.666 | 2.400 | 0.648 |
|  |  |  | region [KB] | -1.417 | -4.018 | 1.130 | 0.141 |
|  |  |  | region [KM] | -2.168 | -5.165 | 0.602 | 0.069 |
|  |  |  | region [metro] | 0.586 | -1.401 | 2.857 | 0.714 |
|  |  |  | region [SB] | 0.098 | -2.050 | 2.282 | 0.537 |
|  |  |  | region [ST] | -1.270 | -3.421 | 0.816 | 0.116 |
|  |  |  | observer effect [none] | 0.234 | 0.025 | 0.446 | 0.987 |
|  |  |  | observer effect [positive] | 1.091 | 0.908 | 1.283 | 1 |
|  |  |  | PC1 - linear | -7.336 | -23.726 | 8.262 | 0.176 |
|  |  |  | PC2 - linear | 24.701 | 10.751 | 38.082 | 1 |
|  |  |  | PC3 - linear | 34.949 | 21.555 | 47.746 | 1 |
|  |  |  | PC1 - quadratic | 5.693 | -6.867 | 17.713 | 0.812 |
|  |  |  | PC2 - quadratic | 0.436 | -12.323 | 13.868 | 0.528 |
|  |  |  | PC3 - quadratic | 7.167 | -5.571 | 20.120 | 0.865 |
|  |  |  | survey year:region [A] | 0.146 | -3.413 | 4.771 | 0.532 |
|  |  |  | survey year:region [B] | -1.016 | -4.207 | 2.848 | 0.249 |
|  |  |  | survey year:region [KB] | 0.217 | -4.326 | 4.873 | 0.539 |
|  |  |  | survey year:region [KM] | 1.567 | -2.645 | 4.188 | 0.807 |
|  |  |  | survey year:region [metro] | 0.036 | -4.215 | 4.417 | 0.507 |
|  |  |  | survey year:region [SB] | -1.270 | -5.908 | 3.310 | 0.285 |
|  |  |  | survey year:region [ST] | 0.917 | -3.732 | 5.304 | 0.647 |
|  |  |  | survey year:region [A] - smoother | 2.092 | 0.062 | 10.979 | 1 |
|  |  |  | survey year:region [B] - smoother | 0.631 | 0.028 | 3.799 | 1 |
|  |  |  | survey year:region [KB] - smoother | 3.431 | 0.657 | 11.994 | 1 |
|  |  |  | survey year:region [KM] - smoother | 0.648 | 0.030 | 3.475 | 1 |
|  |  |  | survey year:region [metro] - smoother | 2.817 | 0.724 | 6.490 | 1 |
|  |  |  | survey year:region [SB] - smoother | 6.423 | 1.246 | 13.789 | 1 |
|  |  |  | survey year:region [ST] - smoother | 4.656 | 1.048 | 13.606 | 1 |
|  |  |  | ID intercept | 3.097 | 2.422 | 3.955 | 1 |
|  |  | binomial (zi) | zi - intercept | -1.029 | -1.520 | -0.588 | 0 |
| Eurasian Woodcock | Scolopax rusticola | pois | intercept [negative, A] | -5.615 | -7.379 | -3.960 | 0 |
|  |  |  | region [B] | -1.728 | -4.405 | 0.410 | 0.059 |
|  |  |  | region [KB] | -1.222 | -3.476 | 0.886 | 0.13 |
|  |  |  | region [KM] | 1.031 | -0.904 | 2.936 | 0.85 |
|  |  |  | region [metro] | -0.436 | -2.632 | 1.534 | 0.335 |
|  |  |  | region [SB] | 0.536 | -1.181 | 2.327 | 0.728 |
|  |  |  | region [ST] | 1.930 | 0.449 | 3.479 | 0.996 |
|  |  |  | observer effect [none] | 1.271 | 0.384 | 2.265 | 0.999 |
|  |  |  | observer effect [positive] | 1.234 | 0.241 | 2.314 | 0.993 |
|  |  |  | PC1 - linear | 10.979 | -2.631 | 24.918 | 0.944 |
|  |  |  | PC2 - linear | 2.136 | -10.359 | 14.842 | 0.63 |
|  |  |  | PC3 - linear | -25.003 | -36.560 | -14.245 | 0 |
|  |  |  | PC1 - quadratic | 1.733 | -7.509 | 11.052 | 0.651 |
|  |  |  | PC2 - quadratic | -8.721 | -20.447 | 2.681 | 0.063 |
|  |  |  | PC3 - quadratic | 4.219 | -6.366 | 15.041 | 0.779 |
|  |  |  | survey year:region [A] | 0.322 | -3.552 | 3.995 | 0.567 |
|  |  |  | survey year:region [B] | 0.512 | -4.369 | 5.163 | 0.585 |
|  |  |  | survey year:region [KB] | 1.432 | -3.418 | 6.011 | 0.718 |
|  |  |  | survey year:region [KM] | 0.711 | -3.990 | 5.184 | 0.619 |
|  |  |  | survey year:region [metro] | -0.548 | -5.157 | 4.105 | 0.402 |
|  |  |  | survey year:region [SB] | 0.545 | -2.987 | 3.414 | 0.649 |
|  |  |  | survey year:region [ST] | 1.053 | -2.706 | 3.748 | 0.753 |
|  |  |  | survey year:region [A] - smoother | 0.823 | 0.036 | 3.886 | 1 |
|  |  |  | survey year:region [B] - smoother | 1.513 | 0.066 | 6.849 | 1 |
|  |  |  | survey year:region [KB] - smoother | 2.081 | 0.162 | 6.883 | 1 |
|  |  |  | survey year:region [KM] - smoother | 2.786 | 0.401 | 8.392 | 1 |
|  |  |  | survey year:region [metro] - smoother | 2.223 | 0.111 | 8.947 | 1 |
|  |  |  | survey year:region [SB] - smoother | 0.602 | 0.027 | 2.893 | 1 |
|  |  |  | survey year:region [ST] - smoother | 0.604 | 0.027 | 2.985 | 1 |
|  |  |  | ID intercept | 1.793 | 1.301 | 2.431 | 1 |
|  |  | binomial (zi) | zi - intercept | -1.870 | -3.701 | -0.878 | 0 |
| Lesser Whitethroat | Sylvia curruca | pois | intercept [negative, A] | -1.162 | -2.017 | -0.357 | 0.002 |
|  |  |  | region [B] | -0.011 | -0.847 | 0.820 | 0.489 |
|  |  |  | region [KB] | 0.498 | -0.533 | 1.505 | 0.835 |
|  |  |  | region [KM] | -0.007 | -1.019 | 0.992 | 0.493 |
|  |  |  | region [metro] | 0.288 | -0.601 | 1.189 | 0.733 |
|  |  |  | region [SB] | 0.652 | -0.365 | 1.690 | 0.898 |
|  |  |  | region [ST] | -0.642 | -1.490 | 0.212 | 0.066 |
|  |  |  | observer effect [none] | 0.281 | -0.104 | 0.659 | 0.926 |
|  |  |  | observer effect [positive] | 0.383 | -0.052 | 0.812 | 0.959 |
|  |  |  | PC1 - linear | -7.806 | -16.404 | 0.439 | 0.03 |
|  |  |  | PC2 - linear | -4.281 | -10.727 | 2.499 | 0.102 |
|  |  |  | PC3 - linear | 17.949 | 12.030 | 24.418 | 1 |
|  |  |  | PC1 - quadratic | -0.978 | -6.256 | 4.359 | 0.35 |
|  |  |  | PC2 - quadratic | -0.057 | -5.640 | 5.443 | 0.491 |
|  |  |  | PC3 - quadratic | -5.562 | -11.805 | 0.156 | 0.028 |
|  |  |  | survey year:region [A] | -0.035 | -3.190 | 3.348 | 0.49 |
|  |  |  | survey year:region [B] | 0.749 | -2.944 | 4.134 | 0.703 |
|  |  |  | survey year:region [KB] | 0.030 | -3.180 | 3.540 | 0.507 |
|  |  |  | survey year:region [KM] | -0.065 | -3.509 | 3.603 | 0.482 |
|  |  |  | survey year:region [metro] | 0.110 | -2.833 | 3.463 | 0.533 |
|  |  |  | survey year:region [SB] | 0.290 | -2.037 | 3.283 | 0.624 |
|  |  |  | survey year:region [ST] | 0.531 | -3.035 | 3.158 | 0.658 |
|  |  |  | survey year:region [A] - smoother | 0.632 | 0.029 | 3.621 | 1 |
|  |  |  | survey year:region [B] - smoother | 0.712 | 0.029 | 4.577 | 1 |
|  |  |  | survey year:region [KB] - smoother | 0.788 | 0.048 | 3.216 | 1 |
|  |  |  | survey year:region [KM] - smoother | 0.732 | 0.032 | 4.002 | 1 |
|  |  |  | survey year:region [metro] - smoother | 0.578 | 0.031 | 2.780 | 1 |
|  |  |  | survey year:region [SB] - smoother | 0.417 | 0.016 | 2.303 | 1 |
|  |  |  | survey year:region [ST] - smoother | 0.679 | 0.031 | 3.052 | 1 |
|  |  |  | ID intercept | 0.963 | 0.700 | 1.269 | 1 |
|  |  | binomial (zi) | zi - intercept | -0.661 | -1.115 | -0.260 | 0 |
| Mallard | Anas platyrhynchos | pois | intercept [negative, A] | 0.698 | -0.155 | 1.477 | 0.945 |
|  |  |  | region [B] | -2.288 | -3.244 | -1.345 | 0 |
|  |  |  | region [KB] | -3.627 | -4.991 | -2.280 | 0 |
|  |  |  | region [KM] | -0.754 | -1.816 | 0.365 | 0.084 |
|  |  |  | region [metro] | -1.683 | -2.688 | -0.684 | 0.001 |
|  |  |  | region [SB] | -1.783 | -2.969 | -0.562 | 0.001 |
|  |  |  | region [ST] | -1.660 | -2.568 | -0.751 | 0 |
|  |  |  | observer effect [none] | 0.113 | -0.176 | 0.409 | 0.773 |
|  |  |  | observer effect [positive] | 0.047 | -0.346 | 0.448 | 0.591 |
|  |  |  | PC1 - linear | -4.822 | -15.352 | 5.594 | 0.177 |
|  |  |  | PC2 - linear | -0.814 | -8.774 | 7.021 | 0.425 |
|  |  |  | PC3 - linear | 10.796 | 4.225 | 17.597 | 0.999 |
|  |  |  | PC1 - quadratic | -0.324 | -6.933 | 6.330 | 0.462 |
|  |  |  | PC2 - quadratic | -4.046 | -11.016 | 2.804 | 0.127 |
|  |  |  | PC3 - quadratic | -1.025 | -8.184 | 6.049 | 0.392 |
|  |  |  | survey year:region [A] | -0.271 | -3.335 | 3.000 | 0.414 |
|  |  |  | survey year:region [B] | -1.056 | -4.328 | 2.671 | 0.265 |
|  |  |  | survey year:region [KB] | 0.856 | -3.517 | 4.668 | 0.658 |
|  |  |  | survey year:region [KM] | -1.022 | -4.111 | 2.930 | 0.27 |
|  |  |  | survey year:region [metro] | -0.343 | -3.306 | 2.790 | 0.383 |
|  |  |  | survey year:region [SB] | 0.503 | -2.302 | 2.879 | 0.718 |
|  |  |  | survey year:region [ST] | -0.010 | -2.475 | 3.233 | 0.497 |
|  |  |  | survey year:region [A] - smoother | 0.796 | 0.131 | 2.946 | 1 |
|  |  |  | survey year:region [B] - smoother | 0.663 | 0.028 | 3.482 | 1 |
|  |  |  | survey year:region [KB] - smoother | 1.119 | 0.054 | 5.362 | 1 |
|  |  |  | survey year:region [KM] - smoother | 0.716 | 0.034 | 3.245 | 1 |
|  |  |  | survey year:region [metro] - smoother | 0.572 | 0.025 | 3.080 | 1 |
|  |  |  | survey year:region [SB] - smoother | 0.384 | 0.019 | 2.337 | 1 |
|  |  |  | survey year:region [ST] - smoother | 0.573 | 0.027 | 2.664 | 1 |
|  |  |  | ID intercept | 1.376 | 1.121 | 1.686 | 1 |
|  |  | binomial (zi) | zi - intercept | -1.320 | -1.814 | -0.895 | 0 |
| Mistle Thrush | Turdus viscivorus | pois | intercept [negative, A] | -0.805 | -1.495 | -0.168 | 0.005 |
|  |  |  | region [B] | 0.485 | -0.236 | 1.209 | 0.908 |
|  |  |  | region [KB] | 0.028 | -0.870 | 0.929 | 0.527 |
|  |  |  | region [KM] | 0.448 | -0.403 | 1.269 | 0.855 |
|  |  |  | region [metro] | 0.705 | -0.124 | 1.547 | 0.951 |
|  |  |  | region [SB] | 0.356 | -0.491 | 1.220 | 0.801 |
|  |  |  | region [ST] | 1.097 | 0.416 | 1.773 | 0.999 |
|  |  |  | observer effect [none] | 0.134 | -0.092 | 0.361 | 0.874 |
|  |  |  | observer effect [positive] | 0.531 | 0.247 | 0.804 | 1 |
|  |  |  | PC1 - linear | 15.048 | 7.998 | 22.005 | 1 |
|  |  |  | PC2 - linear | 4.610 | -0.841 | 10.120 | 0.948 |
|  |  |  | PC3 - linear | -11.662 | -16.212 | -7.314 | 0 |
|  |  |  | PC1 - quadratic | -4.973 | -9.052 | -0.891 | 0.01 |
|  |  |  | PC2 - quadratic | -3.659 | -8.266 | 0.978 | 0.06 |
|  |  |  | PC3 - quadratic | -3.974 | -8.709 | 0.818 | 0.053 |
|  |  |  | survey year:region [A] | -0.531 | -4.767 | 3.665 | 0.404 |
|  |  |  | survey year:region [B] | -0.858 | -3.589 | 2.634 | 0.264 |
|  |  |  | survey year:region [KB] | 0.354 | -3.931 | 2.680 | 0.583 |
|  |  |  | survey year:region [KM] | -0.226 | -3.595 | 3.238 | 0.436 |
|  |  |  | survey year:region [metro] | -0.757 | -5.024 | 3.402 | 0.366 |
|  |  |  | survey year:region [SB] | 0.464 | -3.151 | 4.016 | 0.636 |
|  |  |  | survey year:region [ST] | 0.397 | -2.100 | 3.266 | 0.669 |
|  |  |  | survey year:region [A] - smoother | 1.804 | 0.433 | 5.413 | 1 |
|  |  |  | survey year:region [B] - smoother | 0.513 | 0.028 | 2.737 | 1 |
|  |  |  | survey year:region [KB] - smoother | 0.666 | 0.026 | 3.691 | 1 |
|  |  |  | survey year:region [KM] - smoother | 0.771 | 0.037 | 3.161 | 1 |
|  |  |  | survey year:region [metro] - smoother | 1.844 | 0.403 | 6.157 | 1 |
|  |  |  | survey year:region [SB] - smoother | 1.481 | 0.083 | 3.976 | 1 |
|  |  |  | survey year:region [ST] - smoother | 0.578 | 0.038 | 3.084 | 1 |
|  |  |  | ID intercept | 0.854 | 0.701 | 1.035 | 1 |
|  |  | binomial (zi) | zi - intercept | -1.742 | -2.269 | -1.325 | 0 |
| Willow Tit | Poecile montanus | pois | intercept [negative, A] | -1.643 | -2.583 | -0.716 | 0 |
|  |  |  | region [B] | -1.677 | -2.882 | -0.553 | 0.002 |
|  |  |  | region [KB] | -0.995 | -2.157 | 0.197 | 0.05 |
|  |  |  | region [KM] | 0.075 | -0.994 | 1.149 | 0.553 |
|  |  |  | region [metro] | -0.643 | -1.746 | 0.417 | 0.116 |
|  |  |  | region [SB] | -0.411 | -1.506 | 0.747 | 0.235 |
|  |  |  | region [ST] | 0.246 | -0.646 | 1.130 | 0.711 |
|  |  |  | observer effect [none] | 1.006 | 0.558 | 1.466 | 1 |
|  |  |  | observer effect [positive] | 1.276 | 0.759 | 1.781 | 1 |
|  |  |  | PC1 - linear | 23.290 | 14.106 | 32.448 | 1 |
|  |  |  | PC2 - linear | 12.908 | 6.026 | 19.698 | 1 |
|  |  |  | PC3 - linear | -9.400 | -15.579 | -3.859 | 0 |
|  |  |  | PC1 - quadratic | -6.733 | -11.941 | -1.215 | 0.009 |
|  |  |  | PC2 - quadratic | -2.713 | -8.916 | 3.732 | 0.196 |
|  |  |  | PC3 - quadratic | -2.259 | -8.174 | 3.623 | 0.224 |
|  |  |  | survey year:region [A] | -1.298 | -4.404 | 2.086 | 0.18 |
|  |  |  | survey year:region [B] | -0.829 | -5.278 | 3.901 | 0.364 |
|  |  |  | survey year:region [KB] | 0.616 | -3.019 | 3.302 | 0.67 |
|  |  |  | survey year:region [KM] | -1.683 | -5.222 | 2.804 | 0.212 |
|  |  |  | survey year:region [metro] | -1.639 | -5.968 | 3.440 | 0.257 |
|  |  |  | survey year:region [SB] | -0.513 | -2.683 | 1.947 | 0.238 |
|  |  |  | survey year:region [ST] | -0.526 | -2.729 | 2.967 | 0.327 |
|  |  |  | survey year:region [A] - smoother | 0.668 | 0.026 | 3.385 | 1 |
|  |  |  | survey year:region [B] - smoother | 1.960 | 0.122 | 6.790 | 1 |
|  |  |  | survey year:region [KB] - smoother | 0.564 | 0.022 | 3.037 | 1 |
|  |  |  | survey year:region [KM] - smoother | 1.178 | 0.073 | 4.453 | 1 |
|  |  |  | survey year:region [metro] - smoother | 2.191 | 0.169 | 7.826 | 1 |
|  |  |  | survey year:region [SB] - smoother | 0.303 | 0.011 | 2.088 | 1 |
|  |  |  | survey year:region [ST] - smoother | 0.571 | 0.027 | 3.303 | 1 |
|  |  |  | ID intercept | 1.109 | 0.866 | 1.394 | 1 |
|  |  | binomial (zi) | zi - intercept | -1.257 | -1.721 | -0.836 | 0 |
| Common Kestrel | Falco tinnunculus | pois | intercept [negative, A] | -0.426 | -1.703 | 0.643 | 0.231 |
|  |  |  | region [B] | -0.320 | -1.338 | 0.834 | 0.286 |
|  |  |  | region [KB] | 0.221 | -1.033 | 1.552 | 0.636 |
|  |  |  | region [KM] | 0.096 | -0.991 | 1.299 | 0.561 |
|  |  |  | region [metro] | -0.645 | -1.841 | 0.642 | 0.161 |
|  |  |  | region [SB] | -0.275 | -1.498 | 1.087 | 0.331 |
|  |  |  | region [ST] | -0.063 | -1.085 | 1.091 | 0.457 |
|  |  |  | observer effect [none] | -0.215 | -0.730 | 0.317 | 0.21 |
|  |  |  | observer effect [positive] | 0.236 | -0.366 | 0.864 | 0.776 |
|  |  |  | PC1 - linear | -1.566 | -8.809 | 5.831 | 0.336 |
|  |  |  | PC2 - linear | -0.194 | -5.974 | 5.519 | 0.473 |
|  |  |  | PC3 - linear | 13.068 | 7.294 | 19.081 | 1 |
|  |  |  | PC1 - quadratic | -1.502 | -6.980 | 4.383 | 0.287 |
|  |  |  | PC2 - quadratic | -5.047 | -9.967 | -0.426 | 0.017 |
|  |  |  | PC3 - quadratic | -12.146 | -19.180 | -5.597 | 0 |
|  |  |  | survey year:region [A] | -0.406 | -4.162 | 3.544 | 0.421 |
|  |  |  | survey year:region [B] | 0.989 | -2.593 | 4.017 | 0.739 |
|  |  |  | survey year:region [KB] | 0.817 | -3.290 | 4.528 | 0.673 |
|  |  |  | survey year:region [KM] | -0.208 | -3.513 | 3.447 | 0.445 |
|  |  |  | survey year:region [metro] | -0.045 | -3.804 | 3.743 | 0.489 |
|  |  |  | survey year:region [SB] | 1.645 | -1.779 | 4.704 | 0.856 |
|  |  |  | survey year:region [ST] | -0.613 | -3.410 | 2.921 | 0.334 |
|  |  |  | survey year:region [A] - smoother | 0.877 | 0.041 | 3.986 | 1 |
|  |  |  | survey year:region [B] - smoother | 0.606 | 0.031 | 2.977 | 1 |
|  |  |  | survey year:region [KB] - smoother | 0.835 | 0.039 | 3.801 | 1 |
|  |  |  | survey year:region [KM] - smoother | 0.713 | 0.030 | 3.618 | 1 |
|  |  |  | survey year:region [metro] - smoother | 0.722 | 0.031 | 3.485 | 1 |
|  |  |  | survey year:region [SB] - smoother | 0.665 | 0.026 | 3.000 | 1 |
|  |  |  | survey year:region [ST] - smoother | 0.581 | 0.029 | 2.915 | 1 |
|  |  |  | ID intercept | 0.110 | 0.005 | 0.413 | 1 |
|  |  | binomial (zi) | zi - intercept [A] | 1.054 | -0.166 | 1.918 | 0.96 |
|  |  |  | zi - region [B] | -1.315 | -2.668 | 0.046 | 0.029 |
|  |  |  | zi - region [KB] | 0.358 | -0.822 | 1.849 | 0.723 |
|  |  |  | zi - region [KM] | -0.700 | -1.938 | 0.718 | 0.151 |
|  |  |  | zi - region [metro] | -0.216 | -1.799 | 1.213 | 0.375 |
|  |  |  | zi - region [SB] | -0.934 | -2.014 | 0.384 | 0.069 |
|  |  |  | zi - region [ST] | -0.051 | -1.097 | 1.272 | 0.461 |
| Eurasian Bullfinch | Pyrrhula pyrrhula | pois | intercept [negative, A] | -0.922 | -1.609 | -0.267 | 0.003 |
|  |  |  | region [B] | -0.090 | -0.989 | 0.747 | 0.413 |
|  |  |  | region [KB] | -0.277 | -1.077 | 0.511 | 0.245 |
|  |  |  | region [KM] | 0.624 | -0.150 | 1.406 | 0.945 |
|  |  |  | region [metro] | 0.715 | -0.004 | 1.414 | 0.974 |
|  |  |  | region [SB] | 0.030 | -0.756 | 0.802 | 0.529 |
|  |  |  | region [ST] | 0.182 | -0.511 | 0.873 | 0.704 |
|  |  |  | observer effect [none] | 0.910 | 0.630 | 1.216 | 1 |
|  |  |  | observer effect [positive] | 1.284 | 0.965 | 1.622 | 1 |
|  |  |  | PC1 - linear | 11.164 | 5.204 | 17.217 | 1 |
|  |  |  | PC2 - linear | 10.895 | 6.321 | 15.552 | 1 |
|  |  |  | PC3 - linear | 0.840 | -2.279 | 3.919 | 0.7 |
|  |  |  | PC1 - quadratic | -5.227 | -8.463 | -2.016 | 0.002 |
|  |  |  | PC2 - quadratic | 0.101 | -3.804 | 3.942 | 0.521 |
|  |  |  | PC3 - quadratic | 2.230 | -0.842 | 5.436 | 0.928 |
|  |  |  | survey year:region [A] | 1.127 | -2.445 | 4.502 | 0.761 |
|  |  |  | survey year:region [B] | 1.339 | -3.643 | 5.570 | 0.701 |
|  |  |  | survey year:region [KB] | -0.066 | -4.214 | 3.896 | 0.487 |
|  |  |  | survey year:region [KM] | 0.075 | -3.942 | 3.247 | 0.518 |
|  |  |  | survey year:region [metro] | 0.252 | -2.884 | 2.904 | 0.588 |
|  |  |  | survey year:region [SB] | 0.977 | -1.030 | 2.627 | 0.897 |
|  |  |  | survey year:region [ST] | 0.800 | -1.951 | 3.211 | 0.775 |
|  |  |  | survey year:region [A] - smoother | 0.726 | 0.032 | 3.793 | 1 |
|  |  |  | survey year:region [B] - smoother | 1.518 | 0.099 | 5.623 | 1 |
|  |  |  | survey year:region [KB] - smoother | 1.656 | 0.435 | 4.532 | 1 |
|  |  |  | survey year:region [KM] - smoother | 0.853 | 0.036 | 3.948 | 1 |
|  |  |  | survey year:region [metro] - smoother | 0.435 | 0.021 | 2.295 | 1 |
|  |  |  | survey year:region [SB] - smoother | 0.256 | 0.011 | 1.270 | 1 |
|  |  |  | survey year:region [ST] - smoother | 0.512 | 0.019 | 2.492 | 1 |
|  |  |  | ID intercept | 0.540 | 0.422 | 0.693 | 1 |
|  |  | binomial (zi) | zi - intercept | -1.031 | -1.759 | -0.548 | 0 |
|  |  |  | zi - PC1 - linear | -1.969 | -2.830 | -1.383 | 0 |
|  |  |  | zi - PC2 - linear | -1.430 | -1.878 | -1.065 | 0 |
|  |  |  | zi - PC3 - linear | 1.001 | 0.614 | 1.522 | 1 |
| Grey Wagtail | Motacilla cinerea | pois | intercept [negative, A] | -1.729 | -3.274 | -0.406 | 0.004 |
|  |  |  | region [B] | -2.552 | -4.221 | -0.944 | 0.001 |
|  |  |  | region [KB] | -2.043 | -3.952 | -0.154 | 0.016 |
|  |  |  | region [KM] | -3.234 | -5.917 | -0.956 | 0.001 |
|  |  |  | region [metro] | -4.144 | -6.719 | -2.023 | 0 |
|  |  |  | region [SB] | -0.955 | -2.545 | 0.672 | 0.122 |
|  |  |  | region [ST] | -1.421 | -2.871 | -0.031 | 0.023 |
|  |  |  | observer effect [none] | 0.559 | 0.007 | 1.168 | 0.976 |
|  |  |  | observer effect [positive] | 0.394 | -0.305 | 1.111 | 0.867 |
|  |  |  | PC1 - linear | 6.074 | -7.591 | 19.759 | 0.807 |
|  |  |  | PC2 - linear | 2.461 | -11.348 | 18.374 | 0.629 |
|  |  |  | PC3 - linear | 5.896 | -3.870 | 15.839 | 0.883 |
|  |  |  | PC1 - quadratic | -3.541 | -13.129 | 6.519 | 0.236 |
|  |  |  | PC2 - quadratic | 2.480 | -11.995 | 16.217 | 0.64 |
|  |  |  | PC3 - quadratic | -3.245 | -12.726 | 6.957 | 0.262 |
|  |  |  | survey year:region [A] | -0.751 | -3.548 | 3.084 | 0.304 |
|  |  |  | survey year:region [B] | -0.046 | -4.157 | 4.124 | 0.491 |
|  |  |  | survey year:region [KB] | 0.946 | -3.261 | 4.733 | 0.678 |
|  |  |  | survey year:region [KM] | 0.894 | -3.771 | 5.428 | 0.646 |
|  |  |  | survey year:region [metro] | 0.757 | -4.389 | 5.702 | 0.621 |
|  |  |  | survey year:region [SB] | 0.763 | -1.997 | 2.896 | 0.781 |
|  |  |  | survey year:region [ST] | 0.535 | -2.672 | 3.919 | 0.654 |
|  |  |  | survey year:region [A] - smoother | 0.724 | 0.035 | 3.191 | 1 |
|  |  |  | survey year:region [B] - smoother | 1.008 | 0.038 | 4.945 | 1 |
|  |  |  | survey year:region [KB] - smoother | 0.931 | 0.046 | 4.273 | 1 |
|  |  |  | survey year:region [KM] - smoother | 1.755 | 0.081 | 7.244 | 1 |
|  |  |  | survey year:region [metro] - smoother | 2.798 | 0.362 | 9.564 | 1 |
|  |  |  | survey year:region [SB] - smoother | 0.364 | 0.015 | 2.043 | 1 |
|  |  |  | survey year:region [ST] - smoother | 0.612 | 0.027 | 3.198 | 1 |
|  |  |  | ID intercept | 1.887 | 1.412 | 2.524 | 1 |
|  |  | binomial (zi) | zi - intercept | -5.285 | -9.284 | -2.870 | 0 |
|  |  |  | zi - PC1 - linear | -0.411 | -5.053 | 1.154 | 0.304 |
|  |  |  | zi - PC2 - linear | -6.121 | -10.167 | -0.167 | 0.021 |
|  |  |  | zi - PC3 - linear | -1.051 | -3.783 | 2.620 | 0.214 |
| Long-tailed Tit | Aegithalos caudatus | pois | intercept [negative, A] | -0.060 | -0.584 | 0.446 | 0.406 |
|  |  |  | region [B] | -0.692 | -1.222 | -0.193 | 0.002 |
|  |  |  | region [KB] | -0.566 | -1.236 | 0.108 | 0.051 |
|  |  |  | region [KM] | -0.343 | -0.899 | 0.230 | 0.114 |
|  |  |  | region [metro] | -0.150 | -0.643 | 0.323 | 0.264 |
|  |  |  | region [SB] | -0.193 | -0.839 | 0.438 | 0.273 |
|  |  |  | region [ST] | -0.179 | -0.650 | 0.284 | 0.218 |
|  |  |  | observer effect [none] | 0.471 | 0.213 | 0.750 | 1 |
|  |  |  | observer effect [positive] | 1.093 | 0.794 | 1.398 | 1 |
|  |  |  | PC1 - linear | -8.114 | -14.244 | -2.112 | 0.004 |
|  |  |  | PC2 - linear | 7.325 | 3.424 | 11.374 | 1 |
|  |  |  | PC3 - linear | -1.538 | -4.691 | 1.819 | 0.182 |
|  |  |  | PC1 - quadratic | -2.775 | -6.406 | 0.539 | 0.05 |
|  |  |  | PC2 - quadratic | 1.017 | -2.373 | 4.358 | 0.735 |
|  |  |  | PC3 - quadratic | -2.878 | -6.271 | 0.799 | 0.062 |
|  |  |  | survey year:region [A] | -0.568 | -2.950 | 2.345 | 0.285 |
|  |  |  | survey year:region [B] | -1.728 | -4.813 | 2.760 | 0.193 |
|  |  |  | survey year:region [KB] | 0.608 | -3.027 | 4.348 | 0.657 |
|  |  |  | survey year:region [KM] | -0.403 | -3.805 | 3.005 | 0.378 |
|  |  |  | survey year:region [metro] | -0.706 | -3.506 | 2.687 | 0.279 |
|  |  |  | survey year:region [SB] | -1.260 | -4.621 | 1.857 | 0.166 |
|  |  |  | survey year:region [ST] | -0.293 | -2.545 | 2.814 | 0.403 |
|  |  |  | survey year:region [A] - smoother | 0.405 | 0.015 | 2.369 | 1 |
|  |  |  | survey year:region [B] - smoother | 0.811 | 0.035 | 3.943 | 1 |
|  |  |  | survey year:region [KB] - smoother | 1.037 | 0.059 | 4.152 | 1 |
|  |  |  | survey year:region [KM] - smoother | 0.644 | 0.031 | 2.959 | 1 |
|  |  |  | survey year:region [metro] - smoother | 0.618 | 0.027 | 3.066 | 1 |
|  |  |  | survey year:region [SB] - smoother | 0.830 | 0.084 | 3.009 | 1 |
|  |  |  | survey year:region [ST] - smoother | 0.568 | 0.021 | 2.715 | 1 |
|  |  |  | ID intercept | 0.493 | 0.381 | 0.638 | 1 |
|  |  | binomial (zi) | zi - intercept | -0.858 | -1.255 | -0.574 | 0 |
|  |  |  | zi - PC1 - linear | 0.197 | -0.258 | 0.517 | 0.829 |
|  |  |  | zi - PC2 - linear | -0.769 | -1.058 | -0.511 | 0 |
|  |  |  | zi - PC3 - linear | 0.541 | 0.229 | 0.903 | 1 |
| Common Blackbird | Turdus merula | nb | intercept [negative, A] | 2.887 | 2.623 | 3.150 | 1 |
|  |  |  | region [B] | -0.060 | -0.349 | 0.229 | 0.337 |
|  |  |  | region [KB] | 0.000 | -0.371 | 0.371 | 0.5 |
|  |  |  | region [KM] | 0.245 | -0.106 | 0.577 | 0.915 |
|  |  |  | region [metro] | -0.109 | -0.421 | 0.203 | 0.238 |
|  |  |  | region [SB] | -0.043 | -0.414 | 0.336 | 0.414 |
|  |  |  | region [ST] | 0.178 | -0.105 | 0.459 | 0.893 |
|  |  |  | observer effect [none] | 0.462 | 0.378 | 0.547 | 1 |
|  |  |  | observer effect [positive] | 0.815 | 0.719 | 0.914 | 1 |
|  |  |  | PC1 - linear | -2.998 | -5.986 | 0.066 | 0.028 |
|  |  |  | PC2 - linear | 8.501 | 6.295 | 10.806 | 1 |
|  |  |  | PC3 - linear | 5.458 | 3.593 | 7.188 | 1 |
|  |  |  | PC1 - quadratic | -1.208 | -3.056 | 0.620 | 0.095 |
|  |  |  | PC2 - quadratic | -3.778 | -5.694 | -1.826 | 0 |
|  |  |  | PC3 - quadratic | -0.479 | -2.371 | 1.425 | 0.312 |
|  |  |  | survey year:region [A] | 0.045 | -1.577 | 1.975 | 0.539 |
|  |  |  | survey year:region [B] | -0.068 | -3.247 | 2.091 | 0.468 |
|  |  |  | survey year:region [KB] | 0.126 | -2.250 | 1.739 | 0.596 |
|  |  |  | survey year:region [KM] | -0.151 | -3.297 | 2.048 | 0.421 |
|  |  |  | survey year:region [metro] | -0.354 | -2.628 | 2.681 | 0.354 |
|  |  |  | survey year:region [SB] | 0.253 | -1.536 | 1.185 | 0.733 |
|  |  |  | survey year:region [ST] | 0.123 | -1.807 | 1.702 | 0.612 |
|  |  |  | survey year:region [A] - smoother | 0.213 | 0.008 | 1.397 | 1 |
|  |  |  | survey year:region [B] - smoother | 0.413 | 0.016 | 1.964 | 1 |
|  |  |  | survey year:region [KB] - smoother | 0.232 | 0.008 | 1.421 | 1 |
|  |  |  | survey year:region [KM] - smoother | 0.426 | 0.014 | 2.121 | 1 |
|  |  |  | survey year:region [metro] - smoother | 0.554 | 0.034 | 2.187 | 1 |
|  |  |  | survey year:region [SB] - smoother | 0.138 | 0.005 | 1.002 | 1 |
|  |  |  | survey year:region [ST] - smoother | 0.256 | 0.010 | 1.891 | 1 |
|  |  |  | ID intercept | 0.410 | 0.363 | 0.465 | 1 |
|  |  | binomial (zi) | zi - intercept | -6.795 | -10.129 | -5.124 | 0 |
| Common Goldcrest | Regulus regulus | nb | intercept [negative, A] | -1.288 | -2.198 | -0.418 | 0.002 |
|  |  |  | region [B] | -0.782 | -1.881 | 0.289 | 0.08 |
|  |  |  | region [KB] | 0.636 | -0.512 | 1.838 | 0.858 |
|  |  |  | region [KM] | 0.496 | -0.691 | 1.682 | 0.797 |
|  |  |  | region [metro] | -0.482 | -1.631 | 0.623 | 0.198 |
|  |  |  | region [SB] | 0.983 | -0.165 | 2.146 | 0.951 |
|  |  |  | region [ST] | 1.527 | 0.568 | 2.490 | 0.999 |
|  |  |  | observer effect [none] | 0.770 | 0.496 | 1.038 | 1 |
|  |  |  | observer effect [positive] | 1.202 | 0.870 | 1.538 | 1 |
|  |  |  | PC1 - linear | 24.352 | 15.228 | 33.602 | 1 |
|  |  |  | PC2 - linear | 15.985 | 8.780 | 23.470 | 1 |
|  |  |  | PC3 - linear | -15.748 | -21.625 | -9.883 | 0 |
|  |  |  | PC1 - quadratic | -3.541 | -9.296 | 2.364 | 0.115 |
|  |  |  | PC2 - quadratic | -3.463 | -9.775 | 2.683 | 0.137 |
|  |  |  | PC3 - quadratic | -0.720 | -7.086 | 5.255 | 0.406 |
|  |  |  | survey year:region [A] | -0.386 | -3.849 | 3.846 | 0.416 |
|  |  |  | survey year:region [B] | 0.466 | -3.865 | 4.108 | 0.601 |
|  |  |  | survey year:region [KB] | -1.240 | -3.790 | 2.301 | 0.187 |
|  |  |  | survey year:region [KM] | -0.665 | -4.079 | 3.147 | 0.338 |
|  |  |  | survey year:region [metro] | -0.389 | -4.754 | 3.899 | 0.425 |
|  |  |  | survey year:region [SB] | -1.277 | -3.886 | 1.424 | 0.107 |
|  |  |  | survey year:region [ST] | 0.703 | -2.421 | 3.927 | 0.709 |
|  |  |  | survey year:region [A] - smoother | 1.173 | 0.065 | 5.443 | 1 |
|  |  |  | survey year:region [B] - smoother | 1.131 | 0.056 | 5.768 | 1 |
|  |  |  | survey year:region [KB] - smoother | 0.494 | 0.021 | 2.870 | 1 |
|  |  |  | survey year:region [KM] - smoother | 0.681 | 0.035 | 4.232 | 1 |
|  |  |  | survey year:region [metro] - smoother | 2.077 | 0.296 | 6.999 | 1 |
|  |  |  | survey year:region [SB] - smoother | 0.429 | 0.019 | 2.707 | 1 |
|  |  |  | survey year:region [ST] - smoother | 0.949 | 0.183 | 3.749 | 1 |
|  |  |  | ID intercept | 1.325 | 1.113 | 1.588 | 1 |
|  |  | binomial (zi) | zi - intercept | -3.343 | -4.604 | -2.576 | 0 |
| Common Whitethroat | Sylvia communis | nb | intercept [negative, A] | 0.838 | 0.300 | 1.362 | 0.998 |
|  |  |  | region [B] | -0.609 | -1.162 | -0.061 | 0.016 |
|  |  |  | region [KB] | -0.196 | -0.883 | 0.517 | 0.283 |
|  |  |  | region [KM] | -0.322 | -0.977 | 0.330 | 0.173 |
|  |  |  | region [metro] | 0.107 | -0.519 | 0.718 | 0.641 |
|  |  |  | region [SB] | -0.285 | -0.996 | 0.459 | 0.216 |
|  |  |  | region [ST] | -0.871 | -1.424 | -0.318 | 0.001 |
|  |  |  | observer effect [none] | 0.204 | -0.009 | 0.429 | 0.97 |
|  |  |  | observer effect [positive] | 0.476 | 0.234 | 0.724 | 1 |
|  |  |  | PC1 - linear | -10.502 | -16.487 | -4.514 | 0 |
|  |  |  | PC2 - linear | -19.023 | -23.886 | -14.406 | 0 |
|  |  |  | PC3 - linear | 11.700 | 7.637 | 15.769 | 1 |
|  |  |  | PC1 - quadratic | 3.177 | -0.769 | 6.847 | 0.946 |
|  |  |  | PC2 - quadratic | -2.924 | -6.969 | 0.927 | 0.068 |
|  |  |  | PC3 - quadratic | -7.577 | -12.274 | -3.177 | 0 |
|  |  |  | survey year:region [A] | 0.960 | -1.984 | 3.822 | 0.779 |
|  |  |  | survey year:region [B] | 0.850 | -1.920 | 3.705 | 0.798 |
|  |  |  | survey year:region [KB] | 1.371 | -1.849 | 4.022 | 0.867 |
|  |  |  | survey year:region [KM] | 0.654 | -2.595 | 3.180 | 0.721 |
|  |  |  | survey year:region [metro] | 0.509 | -2.444 | 3.238 | 0.681 |
|  |  |  | survey year:region [SB] | 1.202 | -3.280 | 3.186 | 0.728 |
|  |  |  | survey year:region [ST] | 0.363 | -2.940 | 2.272 | 0.618 |
|  |  |  | survey year:region [A] - smoother | 0.699 | 0.063 | 2.577 | 1 |
|  |  |  | survey year:region [B] - smoother | 0.428 | 0.018 | 3.130 | 1 |
|  |  |  | survey year:region [KB] - smoother | 0.471 | 0.019 | 2.900 | 1 |
|  |  |  | survey year:region [KM] - smoother | 0.424 | 0.019 | 2.255 | 1 |
|  |  |  | survey year:region [metro] - smoother | 0.530 | 0.021 | 2.665 | 1 |
|  |  |  | survey year:region [SB] - smoother | 0.573 | 0.026 | 2.576 | 1 |
|  |  |  | survey year:region [ST] - smoother | 0.504 | 0.021 | 2.290 | 1 |
|  |  |  | ID intercept | 0.782 | 0.670 | 0.918 | 1 |
|  |  | binomial (zi) | zi - intercept | -2.668 | -3.341 | -2.184 | 0 |
| Dunnock | Prunella modularis | nb | intercept [negative, A] | 1.532 | 1.196 | 1.855 | 1 |
|  |  |  | region [B] | 0.028 | -0.320 | 0.378 | 0.561 |
|  |  |  | region [KB] | 0.340 | -0.100 | 0.784 | 0.94 |
|  |  |  | region [KM] | 0.356 | -0.062 | 0.768 | 0.954 |
|  |  |  | region [metro] | 0.312 | -0.059 | 0.680 | 0.952 |
|  |  |  | region [SB] | 0.368 | -0.070 | 0.831 | 0.948 |
|  |  |  | region [ST] | 0.074 | -0.261 | 0.421 | 0.671 |
|  |  |  | observer effect [none] | 0.545 | 0.417 | 0.674 | 1 |
|  |  |  | observer effect [positive] | 0.871 | 0.721 | 1.018 | 1 |
|  |  |  | PC1 - linear | -5.870 | -9.502 | -2.195 | 0.001 |
|  |  |  | PC2 - linear | 8.499 | 5.688 | 11.362 | 1 |
|  |  |  | PC3 - linear | 6.813 | 4.645 | 9.002 | 1 |
|  |  |  | PC1 - quadratic | 1.604 | -0.589 | 3.791 | 0.926 |
|  |  |  | PC2 - quadratic | -4.286 | -6.501 | -1.977 | 0 |
|  |  |  | PC3 - quadratic | -2.038 | -4.340 | 0.356 | 0.047 |
|  |  |  | survey year:region [A] | -0.123 | -2.517 | 1.951 | 0.421 |
|  |  |  | survey year:region [B] | 0.069 | -2.439 | 2.624 | 0.539 |
|  |  |  | survey year:region [KB] | 0.217 | -3.199 | 2.083 | 0.59 |
|  |  |  | survey year:region [KM] | -0.695 | -3.396 | 2.245 | 0.244 |
|  |  |  | survey year:region [metro] | -0.610 | -3.279 | 2.186 | 0.276 |
|  |  |  | survey year:region [SB] | -0.718 | -4.214 | 1.283 | 0.265 |
|  |  |  | survey year:region [ST] | -0.251 | -4.030 | 1.885 | 0.407 |
|  |  |  | survey year:region [A] - smoother | 0.318 | 0.012 | 1.894 | 1 |
|  |  |  | survey year:region [B] - smoother | 0.359 | 0.014 | 2.068 | 1 |
|  |  |  | survey year:region [KB] - smoother | 0.388 | 0.015 | 2.020 | 1 |
|  |  |  | survey year:region [KM] - smoother | 0.412 | 0.015 | 2.527 | 1 |
|  |  |  | survey year:region [metro] - smoother | 0.592 | 0.064 | 2.049 | 1 |
|  |  |  | survey year:region [SB] - smoother | 0.652 | 0.148 | 2.267 | 1 |
|  |  |  | survey year:region [ST] - smoother | 0.808 | 0.054 | 3.701 | 1 |
|  |  |  | ID intercept | 0.484 | 0.423 | 0.555 | 1 |
|  |  | binomial (zi) | zi - intercept | -4.681 | -5.942 | -3.893 | 0 |
| Eurasian Skylark | Alauda arvensis | nb | intercept [negative, A] | -1.873 | -2.980 | -0.810 | 0 |
|  |  |  | region [B] | 1.711 | 0.517 | 2.923 | 0.997 |
|  |  |  | region [KB] | 1.801 | 0.333 | 3.272 | 0.993 |
|  |  |  | region [KM] | -1.059 | -2.593 | 0.468 | 0.087 |
|  |  |  | region [metro] | -1.071 | -2.686 | 0.475 | 0.085 |
|  |  |  | region [SB] | 0.211 | -1.291 | 1.670 | 0.609 |
|  |  |  | region [ST] | -1.378 | -2.582 | -0.137 | 0.015 |
|  |  |  | observer effect [none] | 0.154 | -0.093 | 0.409 | 0.894 |
|  |  |  | observer effect [positive] | 0.329 | 0.007 | 0.635 | 0.977 |
|  |  |  | PC1 - linear | -18.880 | -31.286 | -6.342 | 0.002 |
|  |  |  | PC2 - linear | -42.248 | -53.985 | -30.623 | 0 |
|  |  |  | PC3 - linear | 7.357 | -2.062 | 16.792 | 0.938 |
|  |  |  | PC1 - quadratic | -3.443 | -12.835 | 5.662 | 0.235 |
|  |  |  | PC2 - quadratic | -5.016 | -14.712 | 4.816 | 0.155 |
|  |  |  | PC3 - quadratic | -5.230 | -15.466 | 5.053 | 0.156 |
|  |  |  | survey year:region [A] | -0.430 | -3.415 | 2.411 | 0.34 |
|  |  |  | survey year:region [B] | -0.118 | -2.413 | 2.527 | 0.437 |
|  |  |  | survey year:region [KB] | 0.274 | -2.553 | 3.963 | 0.581 |
|  |  |  | survey year:region [KM] | -0.935 | -4.567 | 2.337 | 0.237 |
|  |  |  | survey year:region [metro] | -0.780 | -4.679 | 3.711 | 0.357 |
|  |  |  | survey year:region [SB] | 0.727 | -1.570 | 3.749 | 0.789 |
|  |  |  | survey year:region [ST] | -1.963 | -4.786 | 1.275 | 0.076 |
|  |  |  | survey year:region [A] - smoother | 0.509 | 0.022 | 2.994 | 1 |
|  |  |  | survey year:region [B] - smoother | 0.340 | 0.015 | 1.890 | 1 |
|  |  |  | survey year:region [KB] - smoother | 0.830 | 0.113 | 3.546 | 1 |
|  |  |  | survey year:region [KM] - smoother | 0.702 | 0.028 | 3.839 | 1 |
|  |  |  | survey year:region [metro] - smoother | 1.035 | 0.044 | 4.611 | 1 |
|  |  |  | survey year:region [SB] - smoother | 0.501 | 0.022 | 2.552 | 1 |
|  |  |  | survey year:region [ST] - smoother | 0.563 | 0.022 | 3.286 | 1 |
|  |  |  | ID intercept | 1.861 | 1.566 | 2.216 | 1 |
|  |  | binomial (zi) | zi - intercept | -2.789 | -3.869 | -2.106 | 0 |
| Eurasian Tree Sparrow | Passer montanus | nb | intercept [negative, A] | -1.626 | -2.606 | -0.637 | 0.001 |
|  |  |  | region [B] | -0.078 | -1.172 | 0.982 | 0.443 |
|  |  |  | region [KB] | 0.449 | -0.834 | 1.693 | 0.761 |
|  |  |  | region [KM] | 1.028 | -0.203 | 2.243 | 0.952 |
|  |  |  | region [metro] | -0.569 | -1.871 | 0.676 | 0.191 |
|  |  |  | region [SB] | 0.298 | -0.951 | 1.502 | 0.685 |
|  |  |  | region [ST] | 0.891 | -0.098 | 1.897 | 0.962 |
|  |  |  | observer effect [none] | 0.626 | 0.282 | 0.979 | 1 |
|  |  |  | observer effect [positive] | 0.941 | 0.529 | 1.360 | 1 |
|  |  |  | PC1 - linear | -8.223 | -18.451 | 1.864 | 0.053 |
|  |  |  | PC2 - linear | -27.065 | -36.043 | -18.530 | 0 |
|  |  |  | PC3 - linear | 13.765 | 6.143 | 21.216 | 1 |
|  |  |  | PC1 - quadratic | -15.417 | -23.741 | -7.356 | 0 |
|  |  |  | PC2 - quadratic | -6.454 | -13.969 | 1.370 | 0.051 |
|  |  |  | PC3 - quadratic | -11.242 | -19.674 | -3.110 | 0.003 |
|  |  |  | survey year:region [A] | -1.360 | -4.614 | 3.373 | 0.27 |
|  |  |  | survey year:region [B] | -2.206 | -4.910 | 2.110 | 0.134 |
|  |  |  | survey year:region [KB] | -0.691 | -4.481 | 2.295 | 0.28 |
|  |  |  | survey year:region [KM] | 0.090 | -3.295 | 3.006 | 0.53 |
|  |  |  | survey year:region [metro] | -2.449 | -7.214 | 2.743 | 0.183 |
|  |  |  | survey year:region [SB] | -0.196 | -3.623 | 2.867 | 0.441 |
|  |  |  | survey year:region [ST] | -1.516 | -3.955 | 1.624 | 0.116 |
|  |  |  | survey year:region [A] - smoother | 1.008 | 0.049 | 4.386 | 1 |
|  |  |  | survey year:region [B] - smoother | 0.644 | 0.030 | 3.053 | 1 |
|  |  |  | survey year:region [KB] - smoother | 0.703 | 0.028 | 3.787 | 1 |
|  |  |  | survey year:region [KM] - smoother | 0.527 | 0.026 | 2.837 | 1 |
|  |  |  | survey year:region [metro] - smoother | 2.199 | 0.133 | 9.393 | 1 |
|  |  |  | survey year:region [SB] - smoother | 0.620 | 0.023 | 3.333 | 1 |
|  |  |  | survey year:region [ST] - smoother | 0.435 | 0.021 | 2.879 | 1 |
|  |  |  | ID intercept | 1.406 | 1.171 | 1.697 | 1 |
|  |  | binomial (zi) | zi - intercept | -1.441 | -1.928 | -1.048 | 0 |
| European Greenfinch | Chloris chloris | nb | intercept [negative, A] | 0.563 | 0.061 | 1.080 | 0.985 |
|  |  |  | region [B] | 0.498 | -0.035 | 1.040 | 0.967 |
|  |  |  | region [KB] | 0.458 | -0.252 | 1.146 | 0.898 |
|  |  |  | region [KM] | 0.317 | -0.316 | 0.945 | 0.84 |
|  |  |  | region [metro] | -0.168 | -0.757 | 0.426 | 0.281 |
|  |  |  | region [SB] | 0.088 | -0.605 | 0.776 | 0.603 |
|  |  |  | region [ST] | 0.391 | -0.148 | 0.914 | 0.926 |
|  |  |  | observer effect [none] | 0.430 | 0.258 | 0.601 | 1 |
|  |  |  | observer effect [positive] | 0.754 | 0.556 | 0.947 | 1 |
|  |  |  | PC1 - linear | -12.977 | -18.885 | -7.291 | 0 |
|  |  |  | PC2 - linear | 9.885 | 5.644 | 14.029 | 1 |
|  |  |  | PC3 - linear | 21.621 | 17.997 | 25.159 | 1 |
|  |  |  | PC1 - quadratic | -2.979 | -6.718 | 0.611 | 0.054 |
|  |  |  | PC2 - quadratic | -6.122 | -9.626 | -2.658 | 0 |
|  |  |  | PC3 - quadratic | -6.580 | -10.400 | -2.733 | 0 |
|  |  |  | survey year:region [A] | -0.081 | -2.355 | 2.956 | 0.466 |
|  |  |  | survey year:region [B] | -0.722 | -3.198 | 2.098 | 0.225 |
|  |  |  | survey year:region [KB] | 0.072 | -3.147 | 3.101 | 0.525 |
|  |  |  | survey year:region [KM] | -0.652 | -4.274 | 3.346 | 0.356 |
|  |  |  | survey year:region [metro] | -1.494 | -4.161 | 2.316 | 0.182 |
|  |  |  | survey year:region [SB] | -0.844 | -3.417 | 1.637 | 0.184 |
|  |  |  | survey year:region [ST] | -0.497 | -3.385 | 1.261 | 0.215 |
|  |  |  | survey year:region [A] - smoother | 0.460 | 0.016 | 2.508 | 1 |
|  |  |  | survey year:region [B] - smoother | 0.358 | 0.014 | 1.874 | 1 |
|  |  |  | survey year:region [KB] - smoother | 0.643 | 0.032 | 3.519 | 1 |
|  |  |  | survey year:region [KM] - smoother | 1.263 | 0.241 | 4.144 | 1 |
|  |  |  | survey year:region [metro] - smoother | 0.653 | 0.050 | 2.688 | 1 |
|  |  |  | survey year:region [SB] - smoother | 0.477 | 0.027 | 1.889 | 1 |
|  |  |  | survey year:region [ST] - smoother | 0.377 | 0.014 | 2.518 | 1 |
|  |  |  | ID intercept | 0.763 | 0.660 | 0.884 | 1 |
|  |  | binomial (zi) | zi - intercept | -4.183 | -6.659 | -3.146 | 0 |
| European Robin | Erithacus rubecula | nb | intercept [negative, A] | 2.126 | 1.707 | 2.546 | 1 |
|  |  |  | region [B] | -0.609 | -1.066 | -0.146 | 0.003 |
|  |  |  | region [KB] | -0.254 | -0.811 | 0.343 | 0.195 |
|  |  |  | region [KM] | 0.275 | -0.272 | 0.822 | 0.843 |
|  |  |  | region [metro] | 0.383 | -0.093 | 0.880 | 0.94 |
|  |  |  | region [SB] | -0.293 | -0.859 | 0.278 | 0.161 |
|  |  |  | region [ST] | 0.497 | 0.045 | 0.943 | 0.984 |
|  |  |  | observer effect [none] | 0.524 | 0.407 | 0.639 | 1 |
|  |  |  | observer effect [positive] | 0.937 | 0.800 | 1.070 | 1 |
|  |  |  | PC1 - linear | 8.682 | 3.994 | 13.346 | 1 |
|  |  |  | PC2 - linear | 15.862 | 12.327 | 19.513 | 1 |
|  |  |  | PC3 - linear | -5.981 | -8.814 | -3.182 | 0 |
|  |  |  | PC1 - quadratic | -2.876 | -5.686 | -0.090 | 0.021 |
|  |  |  | PC2 - quadratic | -2.621 | -5.790 | 0.377 | 0.042 |
|  |  |  | PC3 - quadratic | -1.354 | -4.345 | 1.658 | 0.185 |
|  |  |  | survey year:region [A] | 0.530 | -2.850 | 3.276 | 0.649 |
|  |  |  | survey year:region [B] | 1.736 | -1.701 | 3.755 | 0.884 |
|  |  |  | survey year:region [KB] | 0.646 | -2.812 | 2.533 | 0.685 |
|  |  |  | survey year:region [KM] | 0.184 | -2.590 | 2.923 | 0.581 |
|  |  |  | survey year:region [metro] | 1.169 | -2.235 | 3.167 | 0.783 |
|  |  |  | survey year:region [SB] | 0.797 | -1.752 | 2.313 | 0.844 |
|  |  |  | survey year:region [ST] | 1.291 | -1.268 | 3.739 | 0.89 |
|  |  |  | survey year:region [A] - smoother | 0.758 | 0.064 | 3.057 | 1 |
|  |  |  | survey year:region [B] - smoother | 0.372 | 0.016 | 2.086 | 1 |
|  |  |  | survey year:region [KB] - smoother | 0.472 | 0.021 | 2.145 | 1 |
|  |  |  | survey year:region [KM] - smoother | 0.508 | 0.032 | 2.047 | 1 |
|  |  |  | survey year:region [metro] - smoother | 0.497 | 0.033 | 1.979 | 1 |
|  |  |  | survey year:region [SB] - smoother | 0.277 | 0.013 | 1.672 | 1 |
|  |  |  | survey year:region [ST] - smoother | 0.703 | 0.211 | 2.435 | 1 |
|  |  |  | ID intercept | 0.656 | 0.577 | 0.747 | 1 |
|  |  | binomial (zi) | zi - intercept | -5.110 | -7.126 | -3.998 | 0 |
| Garden Warbler | Sylvia borin | nb | intercept [negative, A] | 0.649 | 0.217 | 1.072 | 0.998 |
|  |  |  | region [B] | -0.700 | -1.137 | -0.251 | 0.001 |
|  |  |  | region [KB] | -0.259 | -0.802 | 0.290 | 0.168 |
|  |  |  | region [KM] | -0.223 | -0.721 | 0.267 | 0.184 |
|  |  |  | region [metro] | -0.226 | -0.713 | 0.238 | 0.17 |
|  |  |  | region [SB] | 0.047 | -0.507 | 0.602 | 0.569 |
|  |  |  | region [ST] | -0.373 | -0.787 | 0.030 | 0.033 |
|  |  |  | observer effect [none] | 0.632 | 0.413 | 0.845 | 1 |
|  |  |  | observer effect [positive] | 0.925 | 0.664 | 1.186 | 1 |
|  |  |  | PC1 - linear | -4.959 | -9.697 | -0.315 | 0.018 |
|  |  |  | PC2 - linear | -1.727 | -5.243 | 1.718 | 0.158 |
|  |  |  | PC3 - linear | -4.503 | -7.349 | -1.697 | 0.001 |
|  |  |  | PC1 - quadratic | -2.948 | -5.765 | -0.014 | 0.024 |
|  |  |  | PC2 - quadratic | 2.026 | -0.936 | 4.994 | 0.913 |
|  |  |  | PC3 - quadratic | -2.578 | -5.544 | 0.508 | 0.05 |
|  |  |  | survey year:region [A] | -1.037 | -3.317 | 1.855 | 0.162 |
|  |  |  | survey year:region [B] | -0.921 | -3.699 | 2.845 | 0.276 |
|  |  |  | survey year:region [KB] | -1.372 | -3.607 | 1.662 | 0.128 |
|  |  |  | survey year:region [KM] | -1.761 | -4.379 | 1.748 | 0.121 |
|  |  |  | survey year:region [metro] | -1.654 | -3.985 | 2.246 | 0.17 |
|  |  |  | survey year:region [SB] | -0.316 | -3.310 | 1.750 | 0.345 |
|  |  |  | survey year:region [ST] | -0.938 | -3.147 | 1.232 | 0.122 |
|  |  |  | survey year:region [A] - smoother | 0.397 | 0.016 | 2.196 | 1 |
|  |  |  | survey year:region [B] - smoother | 0.663 | 0.033 | 2.983 | 1 |
|  |  |  | survey year:region [KB] - smoother | 0.380 | 0.016 | 2.074 | 1 |
|  |  |  | survey year:region [KM] - smoother | 0.489 | 0.024 | 2.549 | 1 |
|  |  |  | survey year:region [metro] - smoother | 0.582 | 0.027 | 2.517 | 1 |
|  |  |  | survey year:region [SB] - smoother | 0.441 | 0.021 | 2.115 | 1 |
|  |  |  | survey year:region [ST] - smoother | 0.307 | 0.016 | 2.172 | 1 |
|  |  |  | ID intercept | 0.544 | 0.441 | 0.659 | 1 |
|  |  | binomial (zi) | zi - intercept | -2.498 | -3.060 | -2.092 | 0 |
| Great Tit | Parus major | nb | intercept [negative, A] | 2.510 | 2.227 | 2.804 | 1 |
|  |  |  | region [B] | -0.360 | -0.677 | -0.055 | 0.01 |
|  |  |  | region [KB] | -0.080 | -0.479 | 0.312 | 0.352 |
|  |  |  | region [KM] | 0.220 | -0.148 | 0.580 | 0.876 |
|  |  |  | region [metro] | 0.120 | -0.225 | 0.442 | 0.761 |
|  |  |  | region [SB] | -0.082 | -0.488 | 0.323 | 0.343 |
|  |  |  | region [ST] | 0.133 | -0.173 | 0.437 | 0.807 |
|  |  |  | observer effect [none] | 0.486 | 0.386 | 0.583 | 1 |
|  |  |  | observer effect [positive] | 0.901 | 0.783 | 1.016 | 1 |
|  |  |  | PC1 - linear | -2.532 | -5.948 | 0.816 | 0.067 |
|  |  |  | PC2 - linear | 9.812 | 7.290 | 12.335 | 1 |
|  |  |  | PC3 - linear | 1.434 | -0.534 | 3.435 | 0.923 |
|  |  |  | PC1 - quadratic | -1.633 | -3.597 | 0.257 | 0.046 |
|  |  |  | PC2 - quadratic | -2.881 | -4.912 | -0.857 | 0.003 |
|  |  |  | PC3 - quadratic | -0.647 | -2.746 | 1.440 | 0.272 |
|  |  |  | survey year:region [A] | 0.916 | -1.900 | 3.952 | 0.806 |
|  |  |  | survey year:region [B] | 0.230 | -2.087 | 2.593 | 0.639 |
|  |  |  | survey year:region [KB] | -0.017 | -1.904 | 2.611 | 0.49 |
|  |  |  | survey year:region [KM] | 0.369 | -2.521 | 3.045 | 0.661 |
|  |  |  | survey year:region [metro] | 0.061 | -1.925 | 2.910 | 0.529 |
|  |  |  | survey year:region [SB] | 1.270 | -1.601 | 3.846 | 0.843 |
|  |  |  | survey year:region [ST] | 1.546 | -0.383 | 4.215 | 0.954 |
|  |  |  | survey year:region [A] - smoother | 0.803 | 0.061 | 2.863 | 1 |
|  |  |  | survey year:region [B] - smoother | 0.303 | 0.014 | 1.660 | 1 |
|  |  |  | survey year:region [KB] - smoother | 0.302 | 0.012 | 1.795 | 1 |
|  |  |  | survey year:region [KM] - smoother | 0.507 | 0.031 | 2.261 | 1 |
|  |  |  | survey year:region [metro] - smoother | 0.382 | 0.025 | 1.775 | 1 |
|  |  |  | survey year:region [SB] - smoother | 0.858 | 0.183 | 2.456 | 1 |
|  |  |  | survey year:region [ST] - smoother | 0.649 | 0.078 | 2.256 | 1 |
|  |  |  | ID intercept | 0.439 | 0.386 | 0.503 | 1 |
|  |  | binomial (zi) | zi - intercept | -5.181 | -7.196 | -4.204 | 0 |
| Hawfinch | Coccothraustes coccothraustes | nb | intercept [negative, A] | -2.050 | -3.118 | -1.050 | 0 |
|  |  |  | region [B] | -0.260 | -1.377 | 0.835 | 0.317 |
|  |  |  | region [KB] | 1.378 | 0.210 | 2.548 | 0.99 |
|  |  |  | region [KM] | 1.069 | -0.019 | 2.172 | 0.973 |
|  |  |  | region [metro] | 0.888 | -0.136 | 1.945 | 0.958 |
|  |  |  | region [SB] | 0.853 | -0.262 | 2.014 | 0.934 |
|  |  |  | region [ST] | 0.499 | -0.453 | 1.487 | 0.843 |
|  |  |  | observer effect [none] | 0.396 | -0.108 | 0.912 | 0.94 |
|  |  |  | observer effect [positive] | 0.884 | 0.293 | 1.494 | 0.997 |
|  |  |  | PC1 - linear | 3.364 | -5.527 | 12.318 | 0.776 |
|  |  |  | PC2 - linear | 24.110 | 16.900 | 31.389 | 1 |
|  |  |  | PC3 - linear | -17.216 | -23.055 | -11.406 | 0 |
|  |  |  | PC1 - quadratic | -8.812 | -15.031 | -3.032 | 0.001 |
|  |  |  | PC2 - quadratic | -1.441 | -7.715 | 4.796 | 0.321 |
|  |  |  | PC3 - quadratic | -7.291 | -13.358 | -1.135 | 0.01 |
|  |  |  | survey year:region [A] | 1.465 | -2.534 | 5.038 | 0.779 |
|  |  |  | survey year:region [B] | 1.575 | -2.977 | 5.125 | 0.751 |
|  |  |  | survey year:region [KB] | 1.237 | -2.517 | 4.510 | 0.772 |
|  |  |  | survey year:region [KM] | 0.224 | -3.706 | 3.640 | 0.555 |
|  |  |  | survey year:region [metro] | 1.400 | -2.829 | 4.740 | 0.771 |
|  |  |  | survey year:region [SB] | 1.123 | -2.659 | 3.839 | 0.783 |
|  |  |  | survey year:region [ST] | 1.562 | -2.971 | 4.553 | 0.751 |
|  |  |  | survey year:region [A] - smoother | 0.980 | 0.055 | 4.346 | 1 |
|  |  |  | survey year:region [B] - smoother | 0.959 | 0.040 | 4.304 | 1 |
|  |  |  | survey year:region [KB] - smoother | 0.786 | 0.036 | 3.412 | 1 |
|  |  |  | survey year:region [KM] - smoother | 0.700 | 0.025 | 3.368 | 1 |
|  |  |  | survey year:region [metro] - smoother | 0.849 | 0.039 | 3.996 | 1 |
|  |  |  | survey year:region [SB] - smoother | 0.606 | 0.030 | 3.349 | 1 |
|  |  |  | survey year:region [ST] - smoother | 0.994 | 0.059 | 3.888 | 1 |
|  |  |  | ID intercept | 1.026 | 0.768 | 1.321 | 1 |
|  |  | binomial (zi) | zi - intercept | -1.058 | -1.708 | -0.575 | 0 |
| Icterine Warbler | Hippolais icterina | nb | intercept [negative, A] | -1.146 | -2.152 | -0.214 | 0.007 |
|  |  |  | region [B] | -0.629 | -1.536 | 0.265 | 0.079 |
|  |  |  | region [KB] | -0.285 | -1.366 | 0.856 | 0.308 |
|  |  |  | region [KM] | -0.459 | -1.520 | 0.615 | 0.196 |
|  |  |  | region [metro] | -0.518 | -1.726 | 0.584 | 0.181 |
|  |  |  | region [SB] | -0.416 | -1.549 | 0.708 | 0.223 |
|  |  |  | region [ST] | -1.090 | -1.958 | -0.245 | 0.006 |
|  |  |  | observer effect [none] | 0.281 | -0.220 | 0.795 | 0.863 |
|  |  |  | observer effect [positive] | 0.640 | 0.020 | 1.248 | 0.978 |
|  |  |  | PC1 - linear | -19.416 | -29.891 | -9.468 | 0 |
|  |  |  | PC2 - linear | -22.984 | -31.923 | -14.830 | 0 |
|  |  |  | PC3 - linear | 6.838 | -0.493 | 14.186 | 0.964 |
|  |  |  | PC1 - quadratic | -2.012 | -9.525 | 5.114 | 0.292 |
|  |  |  | PC2 - quadratic | -1.958 | -9.291 | 5.288 | 0.296 |
|  |  |  | PC3 - quadratic | -0.188 | -7.613 | 6.876 | 0.483 |
|  |  |  | survey year:region [A] | -0.388 | -3.755 | 3.052 | 0.394 |
|  |  |  | survey year:region [B] | 0.461 | -3.324 | 3.775 | 0.614 |
|  |  |  | survey year:region [KB] | 0.366 | -3.525 | 3.779 | 0.589 |
|  |  |  | survey year:region [KM] | -0.537 | -4.352 | 3.053 | 0.378 |
|  |  |  | survey year:region [metro] | 0.294 | -3.558 | 4.518 | 0.559 |
|  |  |  | survey year:region [SB] | -0.603 | -3.777 | 2.991 | 0.343 |
|  |  |  | survey year:region [ST] | -0.766 | -3.484 | 3.095 | 0.307 |
|  |  |  | survey year:region [A] - smoother | 0.782 | 0.044 | 3.338 | 1 |
|  |  |  | survey year:region [B] - smoother | 0.697 | 0.035 | 3.419 | 1 |
|  |  |  | survey year:region [KB] - smoother | 0.765 | 0.036 | 4.030 | 1 |
|  |  |  | survey year:region [KM] - smoother | 0.875 | 0.039 | 5.096 | 1 |
|  |  |  | survey year:region [metro] - smoother | 1.097 | 0.051 | 4.898 | 1 |
|  |  |  | survey year:region [SB] - smoother | 0.649 | 0.027 | 3.461 | 1 |
|  |  |  | survey year:region [ST] - smoother | 0.681 | 0.028 | 3.865 | 1 |
|  |  |  | ID intercept | 0.993 | 0.688 | 1.341 | 1 |
|  |  | binomial (zi) | zi - intercept | -0.237 | -0.742 | 0.170 | 0.14 |
| White Wagtail | Motacilla alba | nb | intercept [negative, A] | 0.591 | 0.133 | 1.034 | 0.994 |
|  |  |  | region [B] | -0.352 | -0.826 | 0.127 | 0.073 |
|  |  |  | region [KB] | -0.295 | -0.911 | 0.305 | 0.158 |
|  |  |  | region [KM] | -0.245 | -0.819 | 0.319 | 0.192 |
|  |  |  | region [metro] | -0.500 | -1.023 | 0.031 | 0.032 |
|  |  |  | region [SB] | -0.054 | -0.631 | 0.530 | 0.426 |
|  |  |  | region [ST] | 0.133 | -0.326 | 0.606 | 0.709 |
|  |  |  | observer effect [none] | 0.369 | 0.219 | 0.524 | 1 |
|  |  |  | observer effect [positive] | 0.738 | 0.568 | 0.912 | 1 |
|  |  |  | PC1 - linear | -2.991 | -7.803 | 1.776 | 0.109 |
|  |  |  | PC2 - linear | -4.260 | -7.929 | -0.586 | 0.012 |
|  |  |  | PC3 - linear | 13.733 | 10.518 | 16.993 | 1 |
|  |  |  | PC1 - quadratic | -2.938 | -6.014 | -0.020 | 0.024 |
|  |  |  | PC2 - quadratic | -3.870 | -6.999 | -0.757 | 0.009 |
|  |  |  | PC3 - quadratic | -7.215 | -10.764 | -3.899 | 0 |
|  |  |  | survey year:region [A] | 0.106 | -2.020 | 2.715 | 0.562 |
|  |  |  | survey year:region [B] | 0.055 | -2.884 | 2.819 | 0.524 |
|  |  |  | survey year:region [KB] | 0.167 | -2.215 | 2.918 | 0.592 |
|  |  |  | survey year:region [KM] | 0.371 | -2.649 | 3.361 | 0.634 |
|  |  |  | survey year:region [metro] | 0.272 | -2.406 | 3.701 | 0.598 |
|  |  |  | survey year:region [SB] | -0.348 | -3.407 | 1.918 | 0.372 |
|  |  |  | survey year:region [ST] | -0.473 | -1.990 | 1.296 | 0.188 |
|  |  |  | survey year:region [A] - smoother | 0.344 | 0.016 | 2.081 | 1 |
|  |  |  | survey year:region [B] - smoother | 0.507 | 0.034 | 2.024 | 1 |
|  |  |  | survey year:region [KB] - smoother | 0.353 | 0.014 | 2.130 | 1 |
|  |  |  | survey year:region [KM] - smoother | 0.622 | 0.038 | 2.757 | 1 |
|  |  |  | survey year:region [metro] - smoother | 0.477 | 0.021 | 2.469 | 1 |
|  |  |  | survey year:region [SB] - smoother | 0.606 | 0.112 | 2.591 | 1 |
|  |  |  | survey year:region [ST] - smoother | 0.225 | 0.009 | 1.426 | 1 |
|  |  |  | ID intercept | 0.608 | 0.510 | 0.721 | 1 |
|  |  | binomial (zi) | zi - intercept | -2.877 | -3.482 | -2.398 | 0 |
| Barn Swallow | Hirundo rustica | nb | intercept [negative, A] | -1.304 | -2.556 | -0.053 | 0.021 |
|  |  |  | region [B] | 0.277 | -1.145 | 1.703 | 0.648 |
|  |  |  | region [KB] | -0.235 | -1.980 | 1.470 | 0.394 |
|  |  |  | region [KM] | 0.879 | -0.865 | 2.568 | 0.85 |
|  |  |  | region [metro] | -0.404 | -2.143 | 1.426 | 0.334 |
|  |  |  | region [SB] | 0.802 | -0.826 | 2.450 | 0.83 |
|  |  |  | region [ST] | 1.340 | -0.076 | 2.682 | 0.969 |
|  |  |  | observer effect [none] | -0.007 | -0.268 | 0.254 | 0.481 |
|  |  |  | observer effect [positive] | 0.218 | -0.076 | 0.525 | 0.924 |
|  |  |  | PC1 - linear | -6.714 | -19.374 | 6.681 | 0.162 |
|  |  |  | PC2 - linear | -27.810 | -39.569 | -16.364 | 0 |
|  |  |  | PC3 - linear | 8.327 | -2.518 | 18.697 | 0.937 |
|  |  |  | PC1 - quadratic | -7.759 | -17.931 | 1.641 | 0.052 |
|  |  |  | PC2 - quadratic | -15.542 | -26.605 | -5.008 | 0.002 |
|  |  |  | PC3 - quadratic | -20.408 | -31.962 | -9.335 | 0 |
|  |  |  | survey year:region [A] | 0.214 | -2.545 | 2.909 | 0.58 |
|  |  |  | survey year:region [B] | -0.579 | -3.229 | 2.803 | 0.312 |
|  |  |  | survey year:region [KB] | -0.891 | -3.566 | 3.403 | 0.312 |
|  |  |  | survey year:region [KM] | -0.809 | -3.687 | 2.947 | 0.304 |
|  |  |  | survey year:region [metro] | -0.549 | -4.342 | 3.374 | 0.378 |
|  |  |  | survey year:region [SB] | 0.201 | -2.532 | 2.734 | 0.595 |
|  |  |  | survey year:region [ST] | 0.511 | -1.636 | 2.628 | 0.757 |
|  |  |  | survey year:region [A] - smoother | 0.468 | 0.023 | 2.455 | 1 |
|  |  |  | survey year:region [B] - smoother | 0.479 | 0.017 | 2.446 | 1 |
|  |  |  | survey year:region [KB] - smoother | 0.746 | 0.030 | 3.469 | 1 |
|  |  |  | survey year:region [KM] - smoother | 0.663 | 0.035 | 2.890 | 1 |
|  |  |  | survey year:region [metro] - smoother | 0.900 | 0.037 | 4.353 | 1 |
|  |  |  | survey year:region [SB] - smoother | 0.440 | 0.022 | 2.229 | 1 |
|  |  |  | survey year:region [ST] - smoother | 0.328 | 0.013 | 1.791 | 1 |
|  |  |  | ID intercept | 2.248 | 1.896 | 2.676 | 1 |
|  |  | binomial (zi) | zi - intercept [A] | -2.115 | -3.858 | -0.779 | 0 |
|  |  |  | zi - region [B] | 0.221 | -2.062 | 2.196 | 0.594 |
|  |  |  | zi - region [KB] | -1.997 | -5.547 | 0.668 | 0.077 |
|  |  |  | zi - region [KM] | 0.056 | -2.506 | 1.974 | 0.521 |
|  |  |  | zi - region [metro] | 1.025 | -2.133 | 3.143 | 0.802 |
|  |  |  | zi - region [SB] | -0.018 | -1.669 | 1.855 | 0.492 |
|  |  |  | zi - region [ST] | -0.703 | -2.645 | 1.174 | 0.226 |
| Eurasian Collared Dove | Streptopelia decaocto | nb | intercept [negative, A] | -1.766 | -2.759 | -0.794 | 0 |
|  |  |  | region [B] | 0.396 | -0.610 | 1.449 | 0.775 |
|  |  |  | region [KB] | 0.012 | -1.415 | 1.436 | 0.507 |
|  |  |  | region [KM] | 0.224 | -0.977 | 1.440 | 0.649 |
|  |  |  | region [metro] | -0.957 | -2.131 | 0.217 | 0.056 |
|  |  |  | region [SB] | -0.600 | -2.025 | 0.791 | 0.199 |
|  |  |  | region [ST] | -0.017 | -1.054 | 1.013 | 0.486 |
|  |  |  | observer effect [none] | 0.524 | 0.101 | 0.947 | 0.993 |
|  |  |  | observer effect [positive] | 0.957 | 0.513 | 1.413 | 1 |
|  |  |  | PC1 - linear | -25.991 | -37.684 | -14.522 | 0 |
|  |  |  | PC2 - linear | 12.626 | 4.114 | 20.794 | 0.999 |
|  |  |  | PC3 - linear | 33.152 | 25.276 | 41.349 | 1 |
|  |  |  | PC1 - quadratic | -2.304 | -11.269 | 5.831 | 0.296 |
|  |  |  | PC2 - quadratic | -7.972 | -15.120 | -0.687 | 0.015 |
|  |  |  | PC3 - quadratic | -6.105 | -13.906 | 1.989 | 0.07 |
|  |  |  | survey year:region [A] | -1.138 | -4.090 | 2.616 | 0.229 |
|  |  |  | survey year:region [B] | -0.523 | -3.219 | 2.612 | 0.327 |
|  |  |  | survey year:region [KB] | -0.320 | -3.884 | 3.156 | 0.418 |
|  |  |  | survey year:region [KM] | 0.007 | -3.303 | 3.389 | 0.502 |
|  |  |  | survey year:region [metro] | -0.379 | -4.225 | 3.861 | 0.419 |
|  |  |  | survey year:region [SB] | 0.482 | -2.656 | 3.828 | 0.645 |
|  |  |  | survey year:region [ST] | 0.168 | -2.495 | 2.781 | 0.577 |
|  |  |  | survey year:region [A] - smoother | 0.707 | 0.035 | 3.305 | 1 |
|  |  |  | survey year:region [B] - smoother | 0.469 | 0.018 | 2.484 | 1 |
|  |  |  | survey year:region [KB] - smoother | 0.771 | 0.034 | 3.771 | 1 |
|  |  |  | survey year:region [KM] - smoother | 0.618 | 0.030 | 3.080 | 1 |
|  |  |  | survey year:region [metro] - smoother | 1.106 | 0.072 | 4.078 | 1 |
|  |  |  | survey year:region [SB] - smoother | 0.543 | 0.022 | 3.140 | 1 |
|  |  |  | survey year:region [ST] - smoother | 0.430 | 0.015 | 2.767 | 1 |
|  |  |  | ID intercept | 1.408 | 1.145 | 1.729 | 1 |
|  |  | binomial (zi) | zi - intercept [A] | -2.777 | -5.058 | -1.173 | 0 |
|  |  |  | zi - region [B] | -1.199 | -4.942 | 1.491 | 0.202 |
|  |  |  | zi - region [KB] | 1.747 | -1.177 | 4.121 | 0.912 |
|  |  |  | zi - region [KM] | -1.424 | -5.335 | 1.466 | 0.177 |
|  |  |  | zi - region [metro] | 1.809 | -0.084 | 4.107 | 0.97 |
|  |  |  | zi - region [SB] | 1.837 | -1.216 | 4.242 | 0.916 |
|  |  |  | zi - region [ST] | -1.146 | -4.950 | 1.611 | 0.224 |
| European Serin | Serinus serinus | nb | intercept [negative, A] | -3.464 | -5.184 | -1.867 | 0 |
|  |  |  | region [B] | 0.637 | -1.020 | 2.326 | 0.769 |
|  |  |  | region [KB] | -0.041 | -1.998 | 1.940 | 0.486 |
|  |  |  | region [KM] | -1.155 | -4.454 | 3.106 | 0.278 |
|  |  |  | region [metro] | 1.665 | -1.468 | 4.414 | 0.861 |
|  |  |  | region [SB] | 0.099 | -1.735 | 2.014 | 0.543 |
|  |  |  | region [ST] | -1.141 | -3.014 | 0.672 | 0.115 |
|  |  |  | observer effect [none] | 0.060 | -0.519 | 0.625 | 0.587 |
|  |  |  | observer effect [positive] | -0.058 | -0.777 | 0.625 | 0.437 |
|  |  |  | PC1 - linear | 4.987 | -9.071 | 19.043 | 0.76 |
|  |  |  | PC2 - linear | 1.097 | -10.286 | 12.730 | 0.573 |
|  |  |  | PC3 - linear | 41.739 | 30.334 | 53.427 | 1 |
|  |  |  | PC1 - quadratic | -8.319 | -19.278 | 2.515 | 0.065 |
|  |  |  | PC2 - quadratic | -13.398 | -24.070 | -1.815 | 0.01 |
|  |  |  | PC3 - quadratic | 2.275 | -8.917 | 13.532 | 0.653 |
|  |  |  | survey year:region [A] | -1.366 | -5.226 | 3.236 | 0.266 |
|  |  |  | survey year:region [B] | -0.885 | -4.074 | 3.033 | 0.296 |
|  |  |  | survey year:region [KB] | -0.443 | -3.733 | 3.212 | 0.39 |
|  |  |  | survey year:region [KM] | -0.785 | -5.500 | 4.040 | 0.374 |
|  |  |  | survey year:region [metro] | -0.170 | -5.040 | 4.661 | 0.474 |
|  |  |  | survey year:region [SB] | -1.627 | -5.050 | 2.822 | 0.226 |
|  |  |  | survey year:region [ST] | -0.772 | -4.746 | 3.766 | 0.37 |
|  |  |  | survey year:region [A] - smoother | 1.299 | 0.061 | 5.066 | 1 |
|  |  |  | survey year:region [B] - smoother | 0.657 | 0.032 | 3.501 | 1 |
|  |  |  | survey year:region [KB] - smoother | 0.662 | 0.027 | 3.299 | 1 |
|  |  |  | survey year:region [KM] - smoother | 1.924 | 0.083 | 8.393 | 1 |
|  |  |  | survey year:region [metro] - smoother | 2.105 | 0.084 | 9.907 | 1 |
|  |  |  | survey year:region [SB] - smoother | 1.250 | 0.178 | 3.784 | 1 |
|  |  |  | survey year:region [ST] - smoother | 1.792 | 0.263 | 5.349 | 1 |
|  |  |  | ID intercept | 2.006 | 1.499 | 2.726 | 1 |
|  |  | binomial (zi) | zi - intercept [A] | -2.048 | -4.928 | 0.142 | 0.035 |
|  |  |  | zi - region [B] | -1.458 | -5.224 | 1.735 | 0.187 |
|  |  |  | zi - region [KB] | -1.322 | -5.155 | 1.877 | 0.211 |
|  |  |  | zi - region [KM] | 2.388 | -3.689 | 6.016 | 0.793 |
|  |  |  | zi - region [metro] | 3.837 | 1.469 | 6.768 | 0.999 |
|  |  |  | zi - region [SB] | -1.670 | -5.198 | 1.401 | 0.147 |
|  |  |  | zi - region [ST] | -0.725 | -4.687 | 2.530 | 0.34 |
| Marsh Warbler | Acrocephalus palustris | nb | intercept [negative, A] | 0.273 | -0.648 | 1.129 | 0.723 |
|  |  |  | region [B] | -1.170 | -2.114 | -0.245 | 0.008 |
|  |  |  | region [KB] | -1.348 | -2.586 | -0.117 | 0.015 |
|  |  |  | region [KM] | -1.203 | -2.374 | 0.007 | 0.026 |
|  |  |  | region [metro] | -0.468 | -1.579 | 0.647 | 0.194 |
|  |  |  | region [SB] | -0.856 | -2.032 | 0.286 | 0.073 |
|  |  |  | region [ST] | -1.865 | -2.902 | -0.824 | 0.001 |
|  |  |  | observer effect [none] | 0.196 | -0.202 | 0.584 | 0.838 |
|  |  |  | observer effect [positive] | 0.358 | -0.102 | 0.803 | 0.939 |
|  |  |  | PC1 - linear | -11.061 | -21.237 | -0.793 | 0.018 |
|  |  |  | PC2 - linear | -12.408 | -20.583 | -4.424 | 0.001 |
|  |  |  | PC3 - linear | 10.487 | 2.869 | 18.255 | 0.997 |
|  |  |  | PC1 - quadratic | -1.772 | -9.246 | 5.472 | 0.317 |
|  |  |  | PC2 - quadratic | -0.220 | -7.198 | 6.654 | 0.478 |
|  |  |  | PC3 - quadratic | -9.689 | -17.644 | -1.914 | 0.006 |
|  |  |  | survey year:region [A] | -1.446 | -3.814 | 1.483 | 0.111 |
|  |  |  | survey year:region [B] | -0.241 | -3.776 | 3.496 | 0.436 |
|  |  |  | survey year:region [KB] | 0.023 | -3.824 | 3.498 | 0.505 |
|  |  |  | survey year:region [KM] | -0.498 | -4.200 | 2.955 | 0.368 |
|  |  |  | survey year:region [metro] | 0.515 | -3.181 | 4.277 | 0.622 |
|  |  |  | survey year:region [SB] | -0.023 | -2.376 | 2.973 | 0.491 |
|  |  |  | survey year:region [ST] | -0.397 | -3.615 | 3.286 | 0.394 |
|  |  |  | survey year:region [A] - smoother | 0.391 | 0.017 | 2.223 | 1 |
|  |  |  | survey year:region [B] - smoother | 0.878 | 0.079 | 3.084 | 1 |
|  |  |  | survey year:region [KB] - smoother | 0.706 | 0.030 | 3.590 | 1 |
|  |  |  | survey year:region [KM] - smoother | 0.789 | 0.039 | 3.828 | 1 |
|  |  |  | survey year:region [metro] - smoother | 0.886 | 0.041 | 3.533 | 1 |
|  |  |  | survey year:region [SB] - smoother | 0.438 | 0.019 | 2.444 | 1 |
|  |  |  | survey year:region [ST] - smoother | 0.930 | 0.058 | 3.562 | 1 |
|  |  |  | ID intercept | 1.272 | 0.958 | 1.611 | 1 |
|  |  | binomial (zi) | zi - intercept [A] | -1.663 | -2.878 | -0.776 | 0 |
|  |  |  | zi - region [B] | 0.176 | -1.409 | 1.608 | 0.594 |
|  |  |  | zi - region [KB] | 2.137 | 0.851 | 3.515 | 0.999 |
|  |  |  | zi - region [KM] | 0.962 | -0.539 | 2.401 | 0.901 |
|  |  |  | zi - region [metro] | 1.249 | 0.129 | 2.561 | 0.986 |
|  |  |  | zi - region [SB] | 1.084 | -0.166 | 2.390 | 0.957 |
|  |  |  | zi - region [ST] | 1.656 | 0.440 | 2.973 | 0.995 |
| Carrion Crow | Corvus corone | nb | intercept [negative, A] | 0.862 | 0.486 | 1.239 | 1 |
|  |  |  | region [B] | -0.452 | -0.843 | -0.073 | 0.012 |
|  |  |  | region [KB] | -0.269 | -0.782 | 0.215 | 0.14 |
|  |  |  | region [KM] | -0.266 | -0.740 | 0.191 | 0.129 |
|  |  |  | region [metro] | -0.467 | -0.945 | -0.029 | 0.019 |
|  |  |  | region [SB] | -0.183 | -0.675 | 0.323 | 0.245 |
|  |  |  | region [ST] | -0.525 | -0.896 | -0.155 | 0.002 |
|  |  |  | observer effect [none] | 0.351 | 0.139 | 0.569 | 0.999 |
|  |  |  | observer effect [positive] | 0.575 | 0.320 | 0.831 | 1 |
|  |  |  | PC1 - linear | -6.545 | -11.358 | -1.955 | 0.003 |
|  |  |  | PC2 - linear | 4.645 | 1.416 | 7.752 | 0.998 |
|  |  |  | PC3 - linear | 2.566 | -0.354 | 5.606 | 0.956 |
|  |  |  | PC1 - quadratic | 1.392 | -1.560 | 4.345 | 0.827 |
|  |  |  | PC2 - quadratic | -0.165 | -2.660 | 2.412 | 0.448 |
|  |  |  | PC3 - quadratic | -4.406 | -7.237 | -1.549 | 0.002 |
|  |  |  | survey year:region [A] | 1.013 | -1.376 | 4.305 | 0.863 |
|  |  |  | survey year:region [B] | 0.865 | -2.669 | 3.550 | 0.75 |
|  |  |  | survey year:region [KB] | 0.513 | -3.027 | 3.308 | 0.665 |
|  |  |  | survey year:region [KM] | -0.354 | -3.710 | 3.403 | 0.41 |
|  |  |  | survey year:region [metro] | 0.363 | -2.622 | 3.397 | 0.627 |
|  |  |  | survey year:region [SB] | 0.054 | -2.115 | 2.610 | 0.527 |
|  |  |  | survey year:region [ST] | 0.401 | -2.256 | 2.079 | 0.687 |
|  |  |  | survey year:region [A] - smoother | 0.543 | 0.021 | 2.841 | 1 |
|  |  |  | survey year:region [B] - smoother | 0.577 | 0.032 | 2.516 | 1 |
|  |  |  | survey year:region [KB] - smoother | 0.569 | 0.026 | 2.702 | 1 |
|  |  |  | survey year:region [KM] - smoother | 0.959 | 0.097 | 3.098 | 1 |
|  |  |  | survey year:region [metro] - smoother | 0.509 | 0.027 | 2.489 | 1 |
|  |  |  | survey year:region [SB] - smoother | 0.419 | 0.026 | 1.934 | 1 |
|  |  |  | survey year:region [ST] - smoother | 0.341 | 0.015 | 1.895 | 1 |
|  |  |  | ID intercept | 0.479 | 0.390 | 0.578 | 1 |
|  |  | binomial (zi) | zi - intercept | -2.365 | -3.303 | -1.838 | 0 |
|  |  |  | zi - PC1 - linear | 0.634 | 0.281 | 1.073 | 0.999 |
|  |  |  | zi - PC2 - linear | -0.131 | -0.861 | 0.364 | 0.332 |
|  |  |  | zi - PC3 - linear | -0.999 | -1.890 | -0.448 | 0 |
| Common Chaffinch | Fringilla coelebs | nb | intercept [negative, A] | 2.922 | 2.602 | 3.243 | 1 |
|  |  |  | region [B] | -0.242 | -0.601 | 0.121 | 0.101 |
|  |  |  | region [KB] | 0.067 | -0.368 | 0.515 | 0.612 |
|  |  |  | region [KM] | 0.505 | 0.088 | 0.911 | 0.993 |
|  |  |  | region [metro] | -0.101 | -0.473 | 0.276 | 0.304 |
|  |  |  | region [SB] | -0.086 | -0.551 | 0.367 | 0.357 |
|  |  |  | region [ST] | 0.616 | 0.273 | 0.963 | 1 |
|  |  |  | observer effect [none] | 0.445 | 0.370 | 0.521 | 1 |
|  |  |  | observer effect [positive] | 0.739 | 0.654 | 0.826 | 1 |
|  |  |  | PC1 - linear | 4.838 | 1.085 | 8.482 | 0.994 |
|  |  |  | PC2 - linear | 5.737 | 2.852 | 8.486 | 1 |
|  |  |  | PC3 - linear | -3.658 | -5.881 | -1.420 | 0 |
|  |  |  | PC1 - quadratic | -0.510 | -2.755 | 1.784 | 0.325 |
|  |  |  | PC2 - quadratic | -3.702 | -6.148 | -1.426 | 0.001 |
|  |  |  | PC3 - quadratic | -0.655 | -3.067 | 1.725 | 0.294 |
|  |  |  | survey year:region [A] | -0.412 | -3.386 | 1.403 | 0.264 |
|  |  |  | survey year:region [B] | -0.335 | -2.980 | 1.753 | 0.268 |
|  |  |  | survey year:region [KB] | -0.181 | -2.725 | 2.467 | 0.384 |
|  |  |  | survey year:region [KM] | -0.109 | -1.903 | 1.739 | 0.409 |
|  |  |  | survey year:region [metro] | -0.296 | -2.825 | 2.085 | 0.353 |
|  |  |  | survey year:region [SB] | -0.384 | -1.428 | 0.574 | 0.122 |
|  |  |  | survey year:region [ST] | 0.209 | -0.764 | 1.787 | 0.719 |
|  |  |  | survey year:region [A] - smoother | 0.461 | 0.017 | 2.297 | 1 |
|  |  |  | survey year:region [B] - smoother | 0.277 | 0.013 | 1.926 | 1 |
|  |  |  | survey year:region [KB] - smoother | 0.486 | 0.015 | 2.536 | 1 |
|  |  |  | survey year:region [KM] - smoother | 0.207 | 0.008 | 1.240 | 1 |
|  |  |  | survey year:region [metro] - smoother | 0.455 | 0.021 | 1.797 | 1 |
|  |  |  | survey year:region [SB] - smoother | 0.114 | 0.005 | 0.768 | 1 |
|  |  |  | survey year:region [ST] - smoother | 0.213 | 0.012 | 1.068 | 1 |
|  |  |  | ID intercept | 0.526 | 0.471 | 0.592 | 1 |
|  |  | binomial (zi) | zi - intercept | -7.812 | -12.166 | -5.560 | 0 |
|  |  |  | zi - PC1 - linear | -0.998 | -4.743 | 1.105 | 0.206 |
|  |  |  | zi - PC2 - linear | -1.598 | -4.620 | 0.362 | 0.057 |
|  |  |  | zi - PC3 - linear | 0.343 | -2.079 | 2.731 | 0.617 |
| Eurasian Jay | Garrulus glandarius | nb | intercept [negative, A] | 0.106 | -0.295 | 0.492 | 0.692 |
|  |  |  | region [B] | -0.302 | -0.736 | 0.134 | 0.083 |
|  |  |  | region [KB] | -0.273 | -0.805 | 0.240 | 0.144 |
|  |  |  | region [KM] | 0.211 | -0.249 | 0.676 | 0.815 |
|  |  |  | region [metro] | 0.372 | -0.034 | 0.808 | 0.964 |
|  |  |  | region [SB] | -0.080 | -0.561 | 0.413 | 0.377 |
|  |  |  | region [ST] | 0.277 | -0.109 | 0.661 | 0.919 |
|  |  |  | observer effect [none] | 0.438 | 0.246 | 0.636 | 1 |
|  |  |  | observer effect [positive] | 0.715 | 0.491 | 0.934 | 1 |
|  |  |  | PC1 - linear | 8.461 | 4.355 | 12.657 | 1 |
|  |  |  | PC2 - linear | 9.244 | 6.164 | 12.447 | 1 |
|  |  |  | PC3 - linear | -5.663 | -8.350 | -2.998 | 0 |
|  |  |  | PC1 - quadratic | -2.684 | -5.002 | -0.361 | 0.011 |
|  |  |  | PC2 - quadratic | 0.138 | -2.501 | 2.833 | 0.539 |
|  |  |  | PC3 - quadratic | -2.713 | -5.454 | -0.076 | 0.021 |
|  |  |  | survey year:region [A] | 1.256 | -1.284 | 4.144 | 0.883 |
|  |  |  | survey year:region [B] | 0.367 | -2.704 | 3.553 | 0.621 |
|  |  |  | survey year:region [KB] | -0.500 | -3.034 | 2.583 | 0.321 |
|  |  |  | survey year:region [KM] | 0.164 | -2.926 | 3.009 | 0.559 |
|  |  |  | survey year:region [metro] | 0.114 | -3.093 | 3.237 | 0.538 |
|  |  |  | survey year:region [SB] | 0.209 | -1.466 | 2.433 | 0.657 |
|  |  |  | survey year:region [ST] | 1.253 | -0.809 | 4.323 | 0.904 |
|  |  |  | survey year:region [A] - smoother | 0.486 | 0.020 | 2.595 | 1 |
|  |  |  | survey year:region [B] - smoother | 0.595 | 0.025 | 2.761 | 1 |
|  |  |  | survey year:region [KB] - smoother | 0.465 | 0.020 | 2.327 | 1 |
|  |  |  | survey year:region [KM] - smoother | 0.474 | 0.022 | 2.876 | 1 |
|  |  |  | survey year:region [metro] - smoother | 0.664 | 0.055 | 2.709 | 1 |
|  |  |  | survey year:region [SB] - smoother | 0.272 | 0.010 | 1.582 | 1 |
|  |  |  | survey year:region [ST] - smoother | 0.649 | 0.044 | 2.716 | 1 |
|  |  |  | ID intercept | 0.426 | 0.340 | 0.529 | 1 |
|  |  | binomial (zi) | zi - intercept | -2.904 | -4.046 | -2.228 | 0 |
|  |  |  | zi - PC1 - linear | -0.804 | -1.719 | -0.255 | 0.001 |
|  |  |  | zi - PC2 - linear | -1.432 | -2.044 | -0.965 | 0 |
|  |  |  | zi - PC3 - linear | 1.552 | 0.914 | 2.418 | 1 |
| European Crested Tit | Lophophanes cristatus | nb | intercept [negative, A] | -1.378 | -2.536 | -0.291 | 0.007 |
|  |  |  | region [B] | -0.129 | -1.479 | 1.224 | 0.427 |
|  |  |  | region [KB] | -0.397 | -1.726 | 0.999 | 0.283 |
|  |  |  | region [KM] | 0.663 | -0.683 | 2.078 | 0.832 |
|  |  |  | region [metro] | -0.093 | -1.559 | 1.331 | 0.45 |
|  |  |  | region [SB] | 0.351 | -0.915 | 1.715 | 0.697 |
|  |  |  | region [ST] | 2.087 | 0.970 | 3.250 | 1 |
|  |  |  | observer effect [none] | 0.533 | 0.188 | 0.878 | 0.999 |
|  |  |  | observer effect [positive] | 0.829 | 0.377 | 1.268 | 1 |
|  |  |  | PC1 - linear | 22.320 | 11.632 | 32.777 | 1 |
|  |  |  | PC2 - linear | 12.814 | 4.678 | 21.004 | 0.999 |
|  |  |  | PC3 - linear | -6.894 | -13.962 | 0.271 | 0.029 |
|  |  |  | PC1 - quadratic | -4.051 | -10.181 | 2.257 | 0.105 |
|  |  |  | PC2 - quadratic | -1.602 | -9.073 | 5.728 | 0.331 |
|  |  |  | PC3 - quadratic | 0.510 | -6.033 | 7.288 | 0.559 |
|  |  |  | survey year:region [A] | -0.865 | -4.164 | 3.126 | 0.325 |
|  |  |  | survey year:region [B] | -0.865 | -5.038 | 3.449 | 0.346 |
|  |  |  | survey year:region [KB] | 0.132 | -3.518 | 3.431 | 0.535 |
|  |  |  | survey year:region [KM] | -0.092 | -3.581 | 3.555 | 0.478 |
|  |  |  | survey year:region [metro] | 1.506 | -3.187 | 5.513 | 0.747 |
|  |  |  | survey year:region [SB] | -0.785 | -2.960 | 1.750 | 0.182 |
|  |  |  | survey year:region [ST] | 0.277 | -2.625 | 3.628 | 0.582 |
|  |  |  | survey year:region [A] - smoother | 0.810 | 0.036 | 3.646 | 1 |
|  |  |  | survey year:region [B] - smoother | 1.029 | 0.044 | 4.935 | 1 |
|  |  |  | survey year:region [KB] - smoother | 0.784 | 0.042 | 3.317 | 1 |
|  |  |  | survey year:region [KM] - smoother | 0.693 | 0.031 | 3.355 | 1 |
|  |  |  | survey year:region [metro] - smoother | 1.218 | 0.064 | 4.462 | 1 |
|  |  |  | survey year:region [SB] - smoother | 0.317 | 0.014 | 2.054 | 1 |
|  |  |  | survey year:region [ST] - smoother | 0.847 | 0.046 | 3.662 | 1 |
|  |  |  | ID intercept | 1.057 | 0.813 | 1.343 | 1 |
|  |  | binomial (zi) | zi - intercept | -1.833 | -3.448 | -0.647 | 0 |
|  |  |  | zi - PC1 - linear | -3.496 | -5.646 | -1.970 | 0 |
|  |  |  | zi - PC2 - linear | -1.455 | -2.294 | -0.753 | 0 |
|  |  |  | zi - PC3 - linear | 2.119 | 1.329 | 3.048 | 1 |
| Fieldfare | Turdus pilaris | nb | intercept [negative, A] | -0.116 | -1.483 | 1.122 | 0.432 |
|  |  |  | region [B] | -1.046 | -2.555 | 0.502 | 0.094 |
|  |  |  | region [KB] | 0.058 | -1.553 | 1.671 | 0.53 |
|  |  |  | region [KM] | -2.616 | -5.057 | -0.577 | 0.005 |
|  |  |  | region [metro] | -1.535 | -3.597 | 0.708 | 0.086 |
|  |  |  | region [SB] | -0.004 | -1.552 | 1.537 | 0.497 |
|  |  |  | region [ST] | -1.135 | -2.580 | 0.306 | 0.062 |
|  |  |  | observer effect [none] | 0.168 | -0.255 | 0.572 | 0.789 |
|  |  |  | observer effect [positive] | 0.356 | -0.112 | 0.822 | 0.931 |
|  |  |  | PC1 - linear | 2.423 | -10.740 | 16.301 | 0.639 |
|  |  |  | PC2 - linear | -7.156 | -17.342 | 3.201 | 0.093 |
|  |  |  | PC3 - linear | 15.774 | 6.162 | 25.279 | 0.999 |
|  |  |  | PC1 - quadratic | -4.330 | -12.782 | 3.822 | 0.155 |
|  |  |  | PC2 - quadratic | -3.072 | -10.929 | 4.743 | 0.22 |
|  |  |  | PC3 - quadratic | -4.602 | -12.840 | 3.793 | 0.146 |
|  |  |  | survey year:region [A] | -0.741 | -3.780 | 2.811 | 0.303 |
|  |  |  | survey year:region [B] | -1.237 | -4.408 | 3.283 | 0.254 |
|  |  |  | survey year:region [KB] | -0.881 | -5.074 | 3.141 | 0.302 |
|  |  |  | survey year:region [KM] | -1.006 | -5.841 | 4.038 | 0.343 |
|  |  |  | survey year:region [metro] | 0.215 | -4.551 | 4.870 | 0.531 |
|  |  |  | survey year:region [SB] | -0.158 | -3.471 | 2.522 | 0.441 |
|  |  |  | survey year:region [ST] | -1.364 | -4.610 | 2.867 | 0.242 |
|  |  |  | survey year:region [A] - smoother | 0.686 | 0.040 | 3.206 | 1 |
|  |  |  | survey year:region [B] - smoother | 0.817 | 0.035 | 3.737 | 1 |
|  |  |  | survey year:region [KB] - smoother | 1.753 | 0.080 | 6.512 | 1 |
|  |  |  | survey year:region [KM] - smoother | 2.387 | 0.130 | 8.601 | 1 |
|  |  |  | survey year:region [metro] - smoother | 1.250 | 0.054 | 5.947 | 1 |
|  |  |  | survey year:region [SB] - smoother | 0.639 | 0.032 | 2.852 | 1 |
|  |  |  | survey year:region [ST] - smoother | 0.838 | 0.043 | 3.968 | 1 |
|  |  |  | ID intercept | 1.304 | 0.956 | 1.797 | 1 |
|  |  | binomial (zi) | zi - intercept | 0.301 | -0.307 | 0.746 | 0.857 |
|  |  |  | zi - PC1 - linear | -1.354 | -1.837 | -0.904 | 0 |
|  |  |  | zi - PC2 - linear | 0.415 | -0.059 | 0.842 | 0.96 |
|  |  |  | zi - PC3 - linear | -0.594 | -1.095 | -0.110 | 0.01 |
| Short-toed Treecreeper | Certhia brachydactyla | nb | intercept [negative, A] | 0.789 | 0.326 | 1.271 | 0.999 |
|  |  |  | region [B] | -0.862 | -1.378 | -0.351 | 0 |
|  |  |  | region [KB] | -0.593 | -1.269 | 0.045 | 0.035 |
|  |  |  | region [KM] | 0.003 | -0.601 | 0.575 | 0.503 |
|  |  |  | region [metro] | 0.002 | -0.529 | 0.539 | 0.503 |
|  |  |  | region [SB] | -0.626 | -1.285 | 0.036 | 0.032 |
|  |  |  | region [ST] | 0.276 | -0.226 | 0.765 | 0.859 |
|  |  |  | observer effect [none] | 0.507 | 0.317 | 0.699 | 1 |
|  |  |  | observer effect [positive] | 0.898 | 0.692 | 1.110 | 1 |
|  |  |  | PC1 - linear | -2.348 | -8.445 | 3.503 | 0.222 |
|  |  |  | PC2 - linear | 11.867 | 7.608 | 16.188 | 1 |
|  |  |  | PC3 - linear | -10.072 | -13.315 | -6.686 | 0 |
|  |  |  | PC1 - quadratic | -7.631 | -11.700 | -3.485 | 0 |
|  |  |  | PC2 - quadratic | -0.766 | -4.320 | 2.857 | 0.335 |
|  |  |  | PC3 - quadratic | -3.029 | -6.650 | 0.657 | 0.053 |
|  |  |  | survey year:region [A] | 0.918 | -1.703 | 3.011 | 0.838 |
|  |  |  | survey year:region [B] | 1.118 | -2.242 | 3.816 | 0.807 |
|  |  |  | survey year:region [KB] | 1.285 | -2.412 | 3.790 | 0.796 |
|  |  |  | survey year:region [KM] | 0.457 | -2.968 | 3.183 | 0.643 |
|  |  |  | survey year:region [metro] | 0.896 | -2.572 | 3.589 | 0.734 |
|  |  |  | survey year:region [SB] | 0.230 | -3.918 | 2.697 | 0.563 |
|  |  |  | survey year:region [ST] | 1.179 | -1.007 | 2.914 | 0.92 |
|  |  |  | survey year:region [A] - smoother | 0.342 | 0.017 | 1.937 | 1 |
|  |  |  | survey year:region [B] - smoother | 0.521 | 0.024 | 2.390 | 1 |
|  |  |  | survey year:region [KB] - smoother | 0.585 | 0.026 | 3.192 | 1 |
|  |  |  | survey year:region [KM] - smoother | 0.621 | 0.035 | 2.597 | 1 |
|  |  |  | survey year:region [metro] - smoother | 0.666 | 0.051 | 2.387 | 1 |
|  |  |  | survey year:region [SB] - smoother | 0.986 | 0.026 | 3.665 | 1 |
|  |  |  | survey year:region [ST] - smoother | 0.285 | 0.011 | 2.007 | 1 |
|  |  |  | ID intercept | 0.710 | 0.603 | 0.833 | 1 |
|  |  | binomial (zi) | zi - intercept | -4.691 | -6.163 | -3.580 | 0 |
|  |  |  | zi - PC1 - linear | 2.555 | 1.880 | 3.395 | 1 |
|  |  |  | zi - PC2 - linear | -1.910 | -2.876 | -1.087 | 0 |
|  |  |  | zi - PC3 - linear | -0.254 | -0.980 | 0.405 | 0.229 |
| Common Buzzard | Buteo buteo | pois | intercept [negative, A] | -0.721 | -1.217 | -0.226 | 0.001 |
|  |  |  | region [B] | 0.183 | -0.233 | 0.606 | 0.806 |
|  |  |  | region [KB] | -0.056 | -0.647 | 0.516 | 0.424 |
|  |  |  | region [KM] | -0.037 | -0.524 | 0.444 | 0.44 |
|  |  |  | region [metro] | -0.047 | -0.615 | 0.466 | 0.434 |
|  |  |  | region [SB] | 0.411 | -0.101 | 0.934 | 0.938 |
|  |  |  | region [ST] | -0.143 | -0.560 | 0.266 | 0.249 |
|  |  |  | observer effect [none] | 0.321 | -0.015 | 0.682 | 0.969 |
|  |  |  | observer effect [positive] | 0.461 | 0.034 | 0.890 | 0.982 |
|  |  |  | PC1 - linear | -5.189 | -9.990 | -0.365 | 0.017 |
|  |  |  | PC2 - linear | -0.456 | -4.175 | 3.135 | 0.403 |
|  |  |  | PC3 - linear | -7.276 | -11.675 | -2.955 | 0.001 |
|  |  |  | PC1 - quadratic | -2.672 | -6.324 | 1.106 | 0.079 |
|  |  |  | PC2 - quadratic | 0.013 | -3.555 | 3.566 | 0.501 |
|  |  |  | PC3 - quadratic | -6.299 | -10.317 | -2.434 | 0.001 |
|  |  |  | survey year:region [A] | 0.564 | -2.314 | 3.633 | 0.694 |
|  |  |  | survey year:region [B] | -0.191 | -3.211 | 3.323 | 0.449 |
|  |  |  | survey year:region [KB] | -0.477 | -3.782 | 3.202 | 0.375 |
|  |  |  | survey year:region [KM] | -0.154 | -3.549 | 3.149 | 0.46 |
|  |  |  | survey year:region [metro] | -0.019 | -3.219 | 3.629 | 0.496 |
|  |  |  | survey year:region [SB] | -0.134 | -2.653 | 3.189 | 0.453 |
|  |  |  | survey year:region [ST] | -0.103 | -2.406 | 3.506 | 0.466 |
|  |  |  | survey year:region [A] - smoother | 0.521 | 0.022 | 2.619 | 1 |
|  |  |  | survey year:region [B] - smoother | 0.566 | 0.026 | 2.912 | 1 |
|  |  |  | survey year:region [KB] - smoother | 0.663 | 0.030 | 3.350 | 1 |
|  |  |  | survey year:region [KM] - smoother | 0.620 | 0.030 | 3.072 | 1 |
|  |  |  | survey year:region [metro] - smoother | 0.633 | 0.030 | 3.104 | 1 |
|  |  |  | survey year:region [SB] - smoother | 0.576 | 0.024 | 2.792 | 1 |
|  |  |  | survey year:region [ST] - smoother | 0.650 | 0.028 | 3.133 | 1 |
|  |  |  | ID intercept | 0.107 | 0.006 | 0.306 | 1 |
|  |  | binomial (zi) | zi - intercept | -0.529 | -1.475 | 0.139 | 0.064 |
|  |  |  | zi - ID intercept | 3.339 | 2.288 | 5.073 | 1 |
| Common House Martin | Delichon urbicum | nb | intercept [negative, A] | 1.540 | 0.385 | 2.601 | 0.994 |
|  |  |  | region [B] | -0.710 | -1.907 | 0.510 | 0.126 |
|  |  |  | region [KB] | -0.455 | -1.965 | 1.038 | 0.271 |
|  |  |  | region [KM] | -0.146 | -1.728 | 1.427 | 0.424 |
|  |  |  | region [metro] | -0.549 | -2.368 | 1.231 | 0.266 |
|  |  |  | region [SB] | -0.744 | -2.102 | 0.641 | 0.144 |
|  |  |  | region [ST] | -0.892 | -2.139 | 0.368 | 0.076 |
|  |  |  | observer effect [none] | 0.000 | -0.454 | 0.476 | 0.5 |
|  |  |  | observer effect [positive] | 0.523 | -0.005 | 1.063 | 0.974 |
|  |  |  | PC1 - linear | -2.327 | -15.004 | 10.151 | 0.361 |
|  |  |  | PC2 - linear | -3.098 | -12.433 | 5.536 | 0.247 |
|  |  |  | PC3 - linear | 22.628 | 9.833 | 33.694 | 1 |
|  |  |  | PC1 - quadratic | -0.563 | -9.747 | 8.203 | 0.451 |
|  |  |  | PC2 - quadratic | 2.227 | -6.138 | 10.573 | 0.701 |
|  |  |  | PC3 - quadratic | -11.168 | -20.788 | -1.298 | 0.014 |
|  |  |  | survey year:region [A] | -0.252 | -3.277 | 3.261 | 0.432 |
|  |  |  | survey year:region [B] | -0.579 | -4.447 | 3.322 | 0.36 |
|  |  |  | survey year:region [KB] | -0.631 | -4.012 | 2.931 | 0.337 |
|  |  |  | survey year:region [KM] | 1.146 | -3.092 | 4.378 | 0.729 |
|  |  |  | survey year:region [metro] | -0.441 | -4.752 | 4.043 | 0.421 |
|  |  |  | survey year:region [SB] | 0.177 | -2.968 | 3.530 | 0.553 |
|  |  |  | survey year:region [ST] | -0.053 | -2.809 | 3.380 | 0.483 |
|  |  |  | survey year:region [A] - smoother | 0.722 | 0.032 | 3.345 | 1 |
|  |  |  | survey year:region [B] - smoother | 1.044 | 0.058 | 4.669 | 1 |
|  |  |  | survey year:region [KB] - smoother | 0.630 | 0.028 | 3.037 | 1 |
|  |  |  | survey year:region [KM] - smoother | 0.820 | 0.032 | 3.605 | 1 |
|  |  |  | survey year:region [metro] - smoother | 1.121 | 0.049 | 5.289 | 1 |
|  |  |  | survey year:region [SB] - smoother | 0.711 | 0.036 | 3.135 | 1 |
|  |  |  | survey year:region [ST] - smoother | 0.683 | 0.036 | 3.021 | 1 |
|  |  |  | ID intercept | 1.191 | 0.900 | 1.585 | 1 |
|  |  | binomial (zi) | zi - intercept | 1.470 | 0.467 | 2.566 | 0.998 |
|  |  |  | zi - ID intercept | 4.671 | 3.342 | 6.780 | 1 |
| Common Linnet | Carduelis cannabina | pois | intercept [negative, A] | -0.580 | -1.380 | 0.201 | 0.075 |
|  |  |  | region [B] | 0.377 | -0.406 | 1.210 | 0.829 |
|  |  |  | region [KB] | 0.761 | -0.200 | 1.750 | 0.938 |
|  |  |  | region [KM] | 0.112 | -0.979 | 1.167 | 0.587 |
|  |  |  | region [metro] | -0.674 | -1.846 | 0.488 | 0.124 |
|  |  |  | region [SB] | 0.366 | -0.617 | 1.357 | 0.767 |
|  |  |  | region [ST] | 0.118 | -0.733 | 0.954 | 0.615 |
|  |  |  | observer effect [none] | 0.317 | 0.043 | 0.584 | 0.989 |
|  |  |  | observer effect [positive] | 0.788 | 0.468 | 1.116 | 1 |
|  |  |  | PC1 - linear | -3.301 | -11.288 | 4.465 | 0.198 |
|  |  |  | PC2 - linear | -3.739 | -10.087 | 2.394 | 0.119 |
|  |  |  | PC3 - linear | 13.249 | 7.462 | 19.338 | 1 |
|  |  |  | PC1 - quadratic | 5.867 | 0.312 | 11.614 | 0.981 |
|  |  |  | PC2 - quadratic | -6.291 | -11.528 | -1.039 | 0.01 |
|  |  |  | PC3 - quadratic | -6.258 | -12.179 | -0.397 | 0.018 |
|  |  |  | survey year:region [A] | -0.097 | -2.803 | 3.524 | 0.472 |
|  |  |  | survey year:region [B] | 1.141 | -1.809 | 3.589 | 0.85 |
|  |  |  | survey year:region [KB] | 0.126 | -2.566 | 3.819 | 0.542 |
|  |  |  | survey year:region [KM] | -0.104 | -3.715 | 3.267 | 0.475 |
|  |  |  | survey year:region [metro] | 0.211 | -3.866 | 4.055 | 0.549 |
|  |  |  | survey year:region [SB] | 0.304 | -2.669 | 2.546 | 0.631 |
|  |  |  | survey year:region [ST] | -0.243 | -3.605 | 1.712 | 0.387 |
|  |  |  | survey year:region [A] - smoother | 0.626 | 0.031 | 3.332 | 1 |
|  |  |  | survey year:region [B] - smoother | 0.361 | 0.015 | 2.051 | 1 |
|  |  |  | survey year:region [KB] - smoother | 0.639 | 0.032 | 2.899 | 1 |
|  |  |  | survey year:region [KM] - smoother | 0.657 | 0.029 | 3.783 | 1 |
|  |  |  | survey year:region [metro] - smoother | 0.823 | 0.040 | 3.886 | 1 |
|  |  |  | survey year:region [SB] - smoother | 0.454 | 0.020 | 2.133 | 1 |
|  |  |  | survey year:region [ST] - smoother | 0.455 | 0.019 | 2.923 | 1 |
|  |  |  | ID intercept | 0.860 | 0.684 | 1.092 | 1 |
|  |  | binomial (zi) | zi - intercept | -0.565 | -1.380 | 0.091 | 0.046 |
|  |  |  | zi - ID intercept | 2.756 | 1.893 | 4.087 | 1 |
| Eurasian Magpie | Pica pica | nb | intercept [negative, A] | 0.188 | -0.282 | 0.628 | 0.788 |
|  |  |  | region [B] | -0.020 | -0.505 | 0.442 | 0.467 |
|  |  |  | region [KB] | 0.197 | -0.435 | 0.847 | 0.731 |
|  |  |  | region [KM] | -0.018 | -0.661 | 0.604 | 0.477 |
|  |  |  | region [metro] | 0.067 | -0.449 | 0.574 | 0.6 |
|  |  |  | region [SB] | -0.046 | -0.652 | 0.561 | 0.44 |
|  |  |  | region [ST] | -0.150 | -0.630 | 0.342 | 0.277 |
|  |  |  | observer effect [none] | 0.371 | 0.158 | 0.588 | 1 |
|  |  |  | observer effect [positive] | 0.452 | 0.212 | 0.689 | 1 |
|  |  |  | PC1 - linear | -9.112 | -14.935 | -3.589 | 0 |
|  |  |  | PC2 - linear | 8.166 | 4.408 | 11.867 | 1 |
|  |  |  | PC3 - linear | 15.686 | 10.217 | 21.043 | 1 |
|  |  |  | PC1 - quadratic | 4.574 | 0.901 | 8.167 | 0.993 |
|  |  |  | PC2 - quadratic | -1.654 | -4.978 | 1.383 | 0.138 |
|  |  |  | PC3 - quadratic | -4.130 | -8.630 | 0.449 | 0.039 |
|  |  |  | survey year:region [A] | 1.278 | -1.973 | 4.431 | 0.842 |
|  |  |  | survey year:region [B] | 0.316 | -2.681 | 3.282 | 0.627 |
|  |  |  | survey year:region [KB] | 0.157 | -3.684 | 2.834 | 0.546 |
|  |  |  | survey year:region [KM] | 0.538 | -2.766 | 3.263 | 0.67 |
|  |  |  | survey year:region [metro] | 0.499 | -1.982 | 4.309 | 0.636 |
|  |  |  | survey year:region [SB] | 0.508 | -1.611 | 2.826 | 0.75 |
|  |  |  | survey year:region [ST] | 0.298 | -1.915 | 2.554 | 0.665 |
|  |  |  | survey year:region [A] - smoother | 0.715 | 0.033 | 3.394 | 1 |
|  |  |  | survey year:region [B] - smoother | 0.508 | 0.024 | 2.444 | 1 |
|  |  |  | survey year:region [KB] - smoother | 0.593 | 0.024 | 2.806 | 1 |
|  |  |  | survey year:region [KM] - smoother | 0.501 | 0.024 | 2.536 | 1 |
|  |  |  | survey year:region [metro] - smoother | 0.721 | 0.045 | 2.853 | 1 |
|  |  |  | survey year:region [SB] - smoother | 0.311 | 0.012 | 1.904 | 1 |
|  |  |  | survey year:region [ST] - smoother | 0.353 | 0.015 | 2.092 | 1 |
|  |  |  | ID intercept | 0.525 | 0.425 | 0.650 | 1 |
|  |  | binomial (zi) | zi - intercept | -5.444 | -9.157 | -3.218 | 0 |
|  |  |  | zi - ID intercept | 8.720 | 5.593 | 14.369 | 1 |
| European Goldfinch | Carduelis carduelis | nb | intercept [negative, A] | 0.260 | -0.326 | 0.836 | 0.815 |
|  |  |  | region [B] | -0.217 | -0.809 | 0.370 | 0.233 |
|  |  |  | region [KB] | -0.117 | -0.882 | 0.663 | 0.382 |
|  |  |  | region [KM] | -0.460 | -1.165 | 0.266 | 0.111 |
|  |  |  | region [metro] | -0.520 | -1.141 | 0.097 | 0.049 |
|  |  |  | region [SB] | -0.164 | -0.939 | 0.608 | 0.339 |
|  |  |  | region [ST] | -0.641 | -1.232 | -0.056 | 0.015 |
|  |  |  | observer effect [none] | 0.174 | -0.107 | 0.454 | 0.889 |
|  |  |  | observer effect [positive] | 0.685 | 0.389 | 0.992 | 1 |
|  |  |  | PC1 - linear | -11.155 | -18.364 | -3.990 | 0.001 |
|  |  |  | PC2 - linear | 1.334 | -3.191 | 6.001 | 0.72 |
|  |  |  | PC3 - linear | 13.755 | 8.920 | 18.623 | 1 |
|  |  |  | PC1 - quadratic | 2.745 | -1.586 | 6.970 | 0.897 |
|  |  |  | PC2 - quadratic | -1.742 | -5.733 | 1.974 | 0.187 |
|  |  |  | PC3 - quadratic | -6.098 | -10.515 | -1.888 | 0.002 |
|  |  |  | survey year:region [A] | 0.430 | -2.106 | 4.007 | 0.638 |
|  |  |  | survey year:region [B] | 1.575 | -1.957 | 4.639 | 0.85 |
|  |  |  | survey year:region [KB] | 0.850 | -2.604 | 4.693 | 0.73 |
|  |  |  | survey year:region [KM] | 0.034 | -3.246 | 3.183 | 0.511 |
|  |  |  | survey year:region [metro] | 1.696 | -1.519 | 3.997 | 0.895 |
|  |  |  | survey year:region [SB] | 0.665 | -2.803 | 3.952 | 0.675 |
|  |  |  | survey year:region [ST] | 0.937 | -2.219 | 3.664 | 0.754 |
|  |  |  | survey year:region [A] - smoother | 0.640 | 0.033 | 3.332 | 1 |
|  |  |  | survey year:region [B] - smoother | 0.701 | 0.050 | 2.940 | 1 |
|  |  |  | survey year:region [KB] - smoother | 0.930 | 0.039 | 4.912 | 1 |
|  |  |  | survey year:region [KM] - smoother | 0.589 | 0.024 | 3.002 | 1 |
|  |  |  | survey year:region [metro] - smoother | 0.378 | 0.014 | 2.034 | 1 |
|  |  |  | survey year:region [SB] - smoother | 0.960 | 0.134 | 3.128 | 1 |
|  |  |  | survey year:region [ST] - smoother | 0.900 | 0.204 | 2.953 | 1 |
|  |  |  | ID intercept | 0.678 | 0.553 | 0.836 | 1 |
|  |  | binomial (zi) | zi - intercept | -1.436 | -2.261 | -0.862 | 0 |
|  |  |  | zi - ID intercept | 2.312 | 1.531 | 3.395 | 1 |
| European Pied Flycatcher | Ficedula hypoleuca | pois | intercept [negative, A] | -0.822 | -2.210 | 0.358 | 0.089 |
|  |  |  | region [B] | -4.126 | -7.433 | -1.798 | 0 |
|  |  |  | region [KB] | -1.922 | -3.531 | -0.331 | 0.01 |
|  |  |  | region [KM] | -0.073 | -1.468 | 1.415 | 0.459 |
|  |  |  | region [metro] | -1.023 | -2.566 | 0.559 | 0.098 |
|  |  |  | region [SB] | -0.812 | -2.165 | 0.683 | 0.137 |
|  |  |  | region [ST] | 1.337 | 0.276 | 2.615 | 0.993 |
|  |  |  | observer effect [none] | 0.137 | -0.319 | 0.597 | 0.713 |
|  |  |  | observer effect [positive] | 0.470 | -0.023 | 1.004 | 0.969 |
|  |  |  | PC1 - linear | 21.317 | 8.781 | 33.489 | 0.999 |
|  |  |  | PC2 - linear | 5.092 | -2.470 | 13.094 | 0.908 |
|  |  |  | PC3 - linear | -10.341 | -18.881 | -1.927 | 0.011 |
|  |  |  | PC1 - quadratic | -6.600 | -12.476 | -0.406 | 0.019 |
|  |  |  | PC2 - quadratic | -4.698 | -12.648 | 4.020 | 0.14 |
|  |  |  | PC3 - quadratic | 5.357 | -1.656 | 12.468 | 0.935 |
|  |  |  | survey year:region [A] | -1.610 | -5.068 | 2.577 | 0.202 |
|  |  |  | survey year:region [B] | -0.108 | -4.980 | 4.774 | 0.484 |
|  |  |  | survey year:region [KB] | -1.363 | -5.275 | 3.488 | 0.278 |
|  |  |  | survey year:region [KM] | 0.899 | -3.920 | 5.195 | 0.647 |
|  |  |  | survey year:region [metro] | 1.072 | -3.624 | 5.366 | 0.684 |
|  |  |  | survey year:region [SB] | 0.250 | -2.930 | 3.194 | 0.593 |
|  |  |  | survey year:region [ST] | 1.384 | -1.787 | 4.342 | 0.839 |
|  |  |  | survey year:region [A] - smoother | 0.918 | 0.042 | 4.295 | 1 |
|  |  |  | survey year:region [B] - smoother | 1.487 | 0.062 | 6.585 | 1 |
|  |  |  | survey year:region [KB] - smoother | 1.176 | 0.061 | 5.123 | 1 |
|  |  |  | survey year:region [KM] - smoother | 2.347 | 0.161 | 8.784 | 1 |
|  |  |  | survey year:region [metro] - smoother | 1.749 | 0.089 | 7.385 | 1 |
|  |  |  | survey year:region [SB] - smoother | 0.517 | 0.023 | 3.081 | 1 |
|  |  |  | survey year:region [ST] - smoother | 0.922 | 0.202 | 3.203 | 1 |
|  |  |  | ID intercept | 0.629 | 0.366 | 1.053 | 1 |
|  |  | binomial (zi) | zi - intercept | 1.364 | 0.337 | 2.512 | 0.993 |
|  |  |  | zi - ID intercept | 3.172 | 2.050 | 5.116 | 1 |
| Firecrest | Regulus ignicapilla | nb | intercept [negative, A] | -0.110 | -0.822 | 0.618 | 0.385 |
|  |  |  | region [B] | -0.855 | -1.862 | 0.055 | 0.033 |
|  |  |  | region [KB] | 0.839 | -0.125 | 1.796 | 0.959 |
|  |  |  | region [KM] | 0.431 | -0.495 | 1.356 | 0.819 |
|  |  |  | region [metro] | 0.365 | -0.654 | 1.341 | 0.77 |
|  |  |  | region [SB] | 0.947 | 0.031 | 1.859 | 0.979 |
|  |  |  | region [ST] | 0.367 | -0.424 | 1.143 | 0.827 |
|  |  |  | observer effect [none] | 0.546 | 0.305 | 0.781 | 1 |
|  |  |  | observer effect [positive] | 0.980 | 0.668 | 1.280 | 1 |
|  |  |  | PC1 - linear | 15.725 | 8.081 | 23.334 | 1 |
|  |  |  | PC2 - linear | 9.291 | 3.926 | 15.427 | 1 |
|  |  |  | PC3 - linear | -7.615 | -12.701 | -2.938 | 0.001 |
|  |  |  | PC1 - quadratic | -0.359 | -4.800 | 3.901 | 0.432 |
|  |  |  | PC2 - quadratic | -3.114 | -8.444 | 1.997 | 0.111 |
|  |  |  | PC3 - quadratic | 1.750 | -3.215 | 6.290 | 0.766 |
|  |  |  | survey year:region [A] | -0.857 | -4.101 | 2.879 | 0.294 |
|  |  |  | survey year:region [B] | 0.580 | -4.275 | 4.624 | 0.597 |
|  |  |  | survey year:region [KB] | 0.157 | -3.022 | 2.949 | 0.558 |
|  |  |  | survey year:region [KM] | -0.334 | -3.980 | 3.171 | 0.413 |
|  |  |  | survey year:region [metro] | 1.383 | -3.271 | 4.932 | 0.728 |
|  |  |  | survey year:region [SB] | -0.953 | -4.561 | 1.664 | 0.187 |
|  |  |  | survey year:region [ST] | 0.246 | -2.640 | 3.788 | 0.579 |
|  |  |  | survey year:region [A] - smoother | 0.723 | 0.030 | 4.687 | 1 |
|  |  |  | survey year:region [B] - smoother | 2.022 | 0.117 | 9.982 | 1 |
|  |  |  | survey year:region [KB] - smoother | 0.510 | 0.024 | 2.555 | 1 |
|  |  |  | survey year:region [KM] - smoother | 0.814 | 0.036 | 4.801 | 1 |
|  |  |  | survey year:region [metro] - smoother | 1.224 | 0.079 | 4.825 | 1 |
|  |  |  | survey year:region [SB] - smoother | 0.948 | 0.037 | 3.280 | 1 |
|  |  |  | survey year:region [ST] - smoother | 0.854 | 0.072 | 3.961 | 1 |
|  |  |  | ID intercept | 0.719 | 0.570 | 0.927 | 1 |
|  |  | binomial (zi) | zi - intercept | -1.474 | -2.997 | -0.414 | 0.003 |
|  |  |  | zi - ID intercept | 5.617 | 3.892 | 8.556 | 1 |
| Great Spotted Woodpecker | Dendrocopos major | nb | intercept [negative, A] | 0.769 | 0.342 | 1.180 | 1 |
|  |  |  | region [B] | -0.144 | -0.655 | 0.364 | 0.286 |
|  |  |  | region [KB] | -0.577 | -1.169 | 0.012 | 0.027 |
|  |  |  | region [KM] | 0.118 | -0.388 | 0.628 | 0.676 |
|  |  |  | region [metro] | 0.019 | -0.445 | 0.492 | 0.534 |
|  |  |  | region [SB] | -0.660 | -1.235 | -0.087 | 0.013 |
|  |  |  | region [ST] | -0.047 | -0.462 | 0.368 | 0.417 |
|  |  |  | observer effect [none] | 0.404 | 0.234 | 0.576 | 1 |
|  |  |  | observer effect [positive] | 0.704 | 0.506 | 0.899 | 1 |
|  |  |  | PC1 - linear | 3.896 | -0.987 | 8.982 | 0.941 |
|  |  |  | PC2 - linear | 12.918 | 9.467 | 16.633 | 1 |
|  |  |  | PC3 - linear | -12.439 | -15.608 | -9.332 | 0 |
|  |  |  | PC1 - quadratic | -5.239 | -8.155 | -2.259 | 0 |
|  |  |  | PC2 - quadratic | 0.147 | -2.784 | 3.307 | 0.537 |
|  |  |  | PC3 - quadratic | -3.356 | -6.425 | -0.318 | 0.017 |
|  |  |  | survey year:region [A] | 1.576 | -1.135 | 4.937 | 0.91 |
|  |  |  | survey year:region [B] | 1.054 | -2.476 | 4.434 | 0.76 |
|  |  |  | survey year:region [KB] | 1.205 | -2.353 | 4.534 | 0.782 |
|  |  |  | survey year:region [KM] | 0.637 | -2.269 | 3.281 | 0.741 |
|  |  |  | survey year:region [metro] | 0.811 | -2.314 | 3.146 | 0.769 |
|  |  |  | survey year:region [SB] | 0.699 | -1.105 | 2.603 | 0.866 |
|  |  |  | survey year:region [ST] | 0.937 | -0.581 | 3.563 | 0.927 |
|  |  |  | survey year:region [A] - smoother | 0.755 | 0.048 | 3.136 | 1 |
|  |  |  | survey year:region [B] - smoother | 0.854 | 0.056 | 3.494 | 1 |
|  |  |  | survey year:region [KB] - smoother | 0.911 | 0.134 | 3.036 | 1 |
|  |  |  | survey year:region [KM] - smoother | 0.419 | 0.018 | 2.618 | 1 |
|  |  |  | survey year:region [metro] - smoother | 0.430 | 0.023 | 1.935 | 1 |
|  |  |  | survey year:region [SB] - smoother | 0.257 | 0.013 | 1.488 | 1 |
|  |  |  | survey year:region [ST] - smoother | 0.382 | 0.015 | 2.019 | 1 |
|  |  |  | ID intercept | 0.568 | 0.481 | 0.671 | 1 |
|  |  | binomial (zi) | zi - intercept | -6.482 | -10.015 | -4.414 | 0 |
|  |  |  | zi - ID intercept | 5.341 | 3.565 | 8.379 | 1 |
| House Sparrow | Passer domesticus | nb | intercept [negative, A] | 1.910 | 1.388 | 2.419 | 1 |
|  |  |  | region [B] | 0.445 | -0.145 | 1.039 | 0.934 |
|  |  |  | region [KB] | 0.556 | -0.168 | 1.299 | 0.933 |
|  |  |  | region [KM] | 0.144 | -0.539 | 0.819 | 0.655 |
|  |  |  | region [metro] | -0.195 | -0.805 | 0.418 | 0.268 |
|  |  |  | region [SB] | 0.788 | 0.093 | 1.503 | 0.988 |
|  |  |  | region [ST] | 0.478 | -0.074 | 1.039 | 0.955 |
|  |  |  | observer effect [none] | 0.509 | 0.354 | 0.668 | 1 |
|  |  |  | observer effect [positive] | 0.921 | 0.741 | 1.099 | 1 |
|  |  |  | PC1 - linear | -14.644 | -21.399 | -7.813 | 0 |
|  |  |  | PC2 - linear | 1.796 | -2.761 | 6.575 | 0.779 |
|  |  |  | PC3 - linear | 12.518 | 7.353 | 17.889 | 1 |
|  |  |  | PC1 - quadratic | 0.693 | -3.762 | 5.031 | 0.621 |
|  |  |  | PC2 - quadratic | -3.090 | -6.870 | 0.786 | 0.058 |
|  |  |  | PC3 - quadratic | -0.762 | -5.950 | 4.161 | 0.38 |
|  |  |  | survey year:region [A] | -0.350 | -3.534 | 1.867 | 0.346 |
|  |  |  | survey year:region [B] | 0.317 | -2.138 | 3.252 | 0.654 |
|  |  |  | survey year:region [KB] | 0.075 | -2.267 | 3.355 | 0.531 |
|  |  |  | survey year:region [KM] | 0.853 | -2.016 | 3.275 | 0.802 |
|  |  |  | survey year:region [metro] | -0.010 | -2.833 | 2.470 | 0.497 |
|  |  |  | survey year:region [SB] | 0.310 | -3.790 | 1.649 | 0.612 |
|  |  |  | survey year:region [ST] | 0.353 | -1.861 | 1.745 | 0.698 |
|  |  |  | survey year:region [A] - smoother | 0.523 | 0.027 | 2.415 | 1 |
|  |  |  | survey year:region [B] - smoother | 0.418 | 0.023 | 2.058 | 1 |
|  |  |  | survey year:region [KB] - smoother | 0.445 | 0.020 | 2.287 | 1 |
|  |  |  | survey year:region [KM] - smoother | 0.374 | 0.016 | 2.045 | 1 |
|  |  |  | survey year:region [metro] - smoother | 0.366 | 0.015 | 2.050 | 1 |
|  |  |  | survey year:region [SB] - smoother | 0.342 | 0.013 | 3.137 | 1 |
|  |  |  | survey year:region [ST] - smoother | 0.270 | 0.011 | 1.606 | 1 |
|  |  |  | ID intercept | 0.778 | 0.678 | 0.896 | 1 |
|  |  | binomial (zi) | zi - intercept | -6.579 | -10.086 | -4.383 | 0 |
|  |  |  | zi - ID intercept | 9.642 | 6.592 | 14.598 | 1 |
| Marsh Tit | Poecile palustris | nb | intercept [negative, A] | 0.240 | -0.317 | 0.775 | 0.8 |
|  |  |  | region [B] | -0.187 | -0.939 | 0.537 | 0.313 |
|  |  |  | region [KB] | -0.266 | -0.985 | 0.417 | 0.22 |
|  |  |  | region [KM] | 0.344 | -0.264 | 0.947 | 0.864 |
|  |  |  | region [metro] | 0.311 | -0.298 | 0.917 | 0.853 |
|  |  |  | region [SB] | 0.082 | -0.629 | 0.790 | 0.594 |
|  |  |  | region [ST] | 0.617 | 0.121 | 1.140 | 0.992 |
|  |  |  | observer effect [none] | 0.306 | 0.045 | 0.551 | 0.99 |
|  |  |  | observer effect [positive] | 0.436 | 0.132 | 0.735 | 0.997 |
|  |  |  | PC1 - linear | 5.669 | -0.004 | 11.542 | 0.975 |
|  |  |  | PC2 - linear | 2.378 | -1.670 | 6.142 | 0.875 |
|  |  |  | PC3 - linear | -10.339 | -13.584 | -7.132 | 0 |
|  |  |  | PC1 - quadratic | -3.633 | -7.017 | -0.356 | 0.015 |
|  |  |  | PC2 - quadratic | 1.163 | -2.220 | 4.695 | 0.753 |
|  |  |  | PC3 - quadratic | 0.792 | -2.391 | 4.013 | 0.684 |
|  |  |  | survey year:region [A] | 0.299 | -2.777 | 3.598 | 0.596 |
|  |  |  | survey year:region [B] | 0.815 | -3.726 | 4.119 | 0.661 |
|  |  |  | survey year:region [KB] | 0.709 | -2.407 | 3.269 | 0.73 |
|  |  |  | survey year:region [KM] | -0.379 | -3.845 | 2.750 | 0.385 |
|  |  |  | survey year:region [metro] | 0.218 | -3.037 | 2.970 | 0.57 |
|  |  |  | survey year:region [SB] | 0.724 | -1.764 | 2.845 | 0.817 |
|  |  |  | survey year:region [ST] | 0.800 | -1.839 | 2.688 | 0.811 |
|  |  |  | survey year:region [A] - smoother | 0.607 | 0.025 | 3.316 | 1 |
|  |  |  | survey year:region [B] - smoother | 0.897 | 0.040 | 4.232 | 1 |
|  |  |  | survey year:region [KB] - smoother | 0.451 | 0.018 | 2.537 | 1 |
|  |  |  | survey year:region [KM] - smoother | 0.683 | 0.037 | 2.886 | 1 |
|  |  |  | survey year:region [metro] - smoother | 0.468 | 0.021 | 2.360 | 1 |
|  |  |  | survey year:region [SB] - smoother | 0.313 | 0.013 | 2.097 | 1 |
|  |  |  | survey year:region [ST] - smoother | 0.342 | 0.015 | 2.208 | 1 |
|  |  |  | ID intercept | 0.550 | 0.435 | 0.692 | 1 |
|  |  | binomial (zi) | zi - intercept | -1.922 | -2.978 | -1.216 | 0 |
|  |  |  | zi - ID intercept | 3.194 | 2.243 | 4.623 | 1 |
| Spotted Flycatcher | Muscicapa striata | pois | intercept [negative, A] | -0.240 | -0.754 | 0.239 | 0.171 |
|  |  |  | region [B] | -0.278 | -0.738 | 0.176 | 0.114 |
|  |  |  | region [KB] | -0.124 | -0.692 | 0.441 | 0.339 |
|  |  |  | region [KM] | 0.080 | -0.446 | 0.616 | 0.611 |
|  |  |  | region [metro] | -0.509 | -1.187 | 0.068 | 0.041 |
|  |  |  | region [SB] | -0.219 | -0.791 | 0.368 | 0.234 |
|  |  |  | region [ST] | 0.217 | -0.168 | 0.648 | 0.862 |
|  |  |  | observer effect [none] | 0.754 | 0.432 | 1.100 | 1 |
|  |  |  | observer effect [positive] | 0.925 | 0.562 | 1.291 | 1 |
|  |  |  | PC1 - linear | -1.623 | -7.529 | 4.030 | 0.288 |
|  |  |  | PC2 - linear | 2.174 | -1.481 | 5.929 | 0.878 |
|  |  |  | PC3 - linear | 0.639 | -2.240 | 3.499 | 0.675 |
|  |  |  | PC1 - quadratic | -0.378 | -4.155 | 3.280 | 0.418 |
|  |  |  | PC2 - quadratic | -2.808 | -6.269 | 0.558 | 0.051 |
|  |  |  | PC3 - quadratic | -0.741 | -3.596 | 2.095 | 0.312 |
|  |  |  | survey year:region [A] | -0.630 | -3.012 | 2.649 | 0.285 |
|  |  |  | survey year:region [B] | -1.009 | -4.157 | 2.445 | 0.233 |
|  |  |  | survey year:region [KB] | 0.224 | -3.186 | 3.494 | 0.564 |
|  |  |  | survey year:region [KM] | -1.449 | -4.825 | 2.716 | 0.208 |
|  |  |  | survey year:region [metro] | 0.928 | -3.004 | 4.380 | 0.693 |
|  |  |  | survey year:region [SB] | -1.064 | -4.821 | 2.044 | 0.208 |
|  |  |  | survey year:region [ST] | -0.631 | -2.841 | 1.364 | 0.174 |
|  |  |  | survey year:region [A] - smoother | 0.485 | 0.019 | 2.473 | 1 |
|  |  |  | survey year:region [B] - smoother | 0.596 | 0.027 | 3.048 | 1 |
|  |  |  | survey year:region [KB] - smoother | 0.671 | 0.034 | 3.229 | 1 |
|  |  |  | survey year:region [KM] - smoother | 0.860 | 0.036 | 5.151 | 1 |
|  |  |  | survey year:region [metro] - smoother | 0.879 | 0.046 | 3.851 | 1 |
|  |  |  | survey year:region [SB] - smoother | 0.927 | 0.055 | 3.802 | 1 |
|  |  |  | survey year:region [ST] - smoother | 0.295 | 0.012 | 2.125 | 1 |
|  |  |  | ID intercept | 0.421 | 0.313 | 0.551 | 1 |
|  |  | binomial (zi) | zi - intercept | -0.335 | -0.812 | 0.075 | 0.056 |
|  |  |  | zi - ID intercept | 2.155 | 1.580 | 2.951 | 1 |
| Tawny Owl | Strix aluco | pois | intercept [negative, A] | -0.369 | -0.980 | 0.192 | 0.098 |
|  |  |  | region [B] | -0.363 | -1.089 | 0.289 | 0.141 |
|  |  |  | region [KB] | -0.400 | -1.155 | 0.334 | 0.134 |
|  |  |  | region [KM] | -0.446 | -1.157 | 0.225 | 0.091 |
|  |  |  | region [metro] | -0.359 | -1.066 | 0.307 | 0.148 |
|  |  |  | region [SB] | -0.139 | -0.812 | 0.516 | 0.347 |
|  |  |  | region [ST] | 0.034 | -0.452 | 0.534 | 0.556 |
|  |  |  | observer effect [none] | 0.191 | -0.168 | 0.567 | 0.845 |
|  |  |  | observer effect [positive] | 0.287 | -0.189 | 0.752 | 0.884 |
|  |  |  | PC1 - linear | -0.328 | -6.559 | 6.263 | 0.461 |
|  |  |  | PC2 - linear | 2.952 | -1.652 | 7.726 | 0.901 |
|  |  |  | PC3 - linear | -7.425 | -13.079 | -1.564 | 0.008 |
|  |  |  | PC1 - quadratic | -3.390 | -7.429 | 0.238 | 0.032 |
|  |  |  | PC2 - quadratic | -2.822 | -7.151 | 1.559 | 0.096 |
|  |  |  | PC3 - quadratic | -3.986 | -8.611 | 0.487 | 0.041 |
|  |  |  | survey year:region [A] | -0.362 | -3.245 | 3.079 | 0.388 |
|  |  |  | survey year:region [B] | 0.940 | -3.301 | 4.391 | 0.684 |
|  |  |  | survey year:region [KB] | -0.972 | -4.059 | 2.757 | 0.262 |
|  |  |  | survey year:region [KM] | -0.773 | -4.570 | 3.199 | 0.33 |
|  |  |  | survey year:region [metro] | 0.664 | -3.648 | 4.366 | 0.635 |
|  |  |  | survey year:region [SB] | 0.057 | -2.339 | 3.390 | 0.521 |
|  |  |  | survey year:region [ST] | -0.471 | -2.741 | 2.274 | 0.306 |
|  |  |  | survey year:region [A] - smoother | 0.570 | 0.026 | 2.738 | 1 |
|  |  |  | survey year:region [B] - smoother | 0.801 | 0.035 | 3.852 | 1 |
|  |  |  | survey year:region [KB] - smoother | 0.633 | 0.027 | 3.152 | 1 |
|  |  |  | survey year:region [KM] - smoother | 0.873 | 0.045 | 4.585 | 1 |
|  |  |  | survey year:region [metro] - smoother | 0.834 | 0.043 | 4.440 | 1 |
|  |  |  | survey year:region [SB] - smoother | 0.470 | 0.019 | 2.593 | 1 |
|  |  |  | survey year:region [ST] - smoother | 0.405 | 0.014 | 2.173 | 1 |
|  |  |  | ID intercept | 0.224 | 0.020 | 0.453 | 1 |
|  |  | binomial (zi) | zi - intercept | 0.231 | -0.667 | 0.992 | 0.714 |
|  |  |  | zi - ID intercept | 3.425 | 2.341 | 5.369 | 1 |

### 7 Species list

Table 7.1: Species list with proportion of zeros, number of surveys, as well as model characteristics.

|  | | prop. of zeros [%] | | |  | | |
| --- | --- | --- | --- | --- | --- | --- | --- |
| species | scientific | total | atlantic | continental | # of surveys | model family | binomial model coefficients |
| Greylag Goose | Anser anser | 96.74 | 94.91 | 99.59 | 614 |  |  |
| Canada Goose | Branta canadensis | 91.52 | 89.78 | 94.19 | 613 |  |  |
| Mute Swan | Cygnus olor | 95.77 | 94.37 | 97.93 | 614 |  |  |
| Egyptian Goose | Alopochen aegyptiaca | 90.39 | 86.06 | 97.10 | 614 |  |  |
| Ruddy Shelduck | Tadorna ferruginea | 99.35 | 99.20 | 99.59 | 614 |  |  |
| Common Shelduck | Tadorna tadorna | 99.67 | 99.46 | 100.00 | 614 |  |  |
| Muscovy Duck | Cairina moschata f. domestica | 99.84 | 99.73 | 100.00 | 614 |  |  |
| Wood Duck | Aix sponsa | 99.84 | 100.00 | 99.59 | 614 |  |  |
| Mandarin Duck | Aix galericulata | 99.67 | 99.73 | 99.59 | 614 |  |  |
| Garganey | Anas querquedula | 99.35 | 98.93 | 100.00 | 614 |  |  |
| Northern Shoveler | Anas clypeata | 99.35 | 98.93 | 100.00 | 614 |  |  |
| Gadwall | Anas strepera | 99.02 | 98.39 | 100.00 | 614 |  |  |
| Mallard | Anas platyrhynchos | 59.45 | 52.82 | 69.71 | 614 | zip | ~ 1 |
| Domestic Duck | Anas platyrhynchos f. domestica | 99.67 | 99.46 | 100.00 | 614 |  |  |
| Common Teal | Anas crecca | 99.19 | 98.66 | 100.00 | 614 |  |  |
| Common Pochard | Aythya ferina | 99.19 | 99.73 | 98.34 | 614 |  |  |
| Tufted Duck | Aythya fuligula | 94.62 | 93.28 | 96.68 | 613 |  |  |
| Grey Partridge | Perdix perdix | 83.36 | 75.54 | 95.44 | 613 |  |  |
| Common Quail | Coturnix coturnix | 94.12 | 93.80 | 94.61 | 612 |  |  |
| Common Pheasant | Phasianus colchicus | 53.34 | 30.91 | 87.97 | 613 | nb | ~ 1 |
| Little Grebe | Tachybaptus ruficollis | 98.21 | 97.58 | 99.17 | 613 |  |  |
| Great Crested Grebe | Podiceps cristatus | 95.77 | 93.30 | 99.59 | 614 |  |  |
| Black Stork | Ciconia nigra | 99.67 | 100.00 | 99.17 | 614 |  |  |
| White Stork | Ciconia ciconia | 99.84 | 99.73 | 100.00 | 614 |  |  |
| Grey Heron | Ardea cinerea | 98.86 | 98.93 | 98.76 | 614 |  |  |
| European Honey Buzzard | Pernis apivorus | 99.67 | 99.73 | 99.59 | 614 |  |  |
| Red Kite | Milvus milvus | 97.39 | 98.93 | 95.02 | 614 |  |  |
| Black Kite | Milvus migrans | 99.51 | 99.46 | 99.59 | 614 |  |  |
| Western Marsh Harrier | Circus aeruginosus | 99.35 | 98.93 | 100.00 | 614 |  |  |
| Eurasian Sparrowhawk | Accipiter nisus | 92.51 | 91.42 | 94.19 | 614 |  |  |
| Northern Goshawk | Accipiter gentilis | 91.69 | 90.88 | 92.95 | 614 |  |  |
| Common Buzzard | Buteo buteo | 57.59 | 59.95 | 53.94 | 613 | zip | ~ (1\|ID) |
| Common Kestrel | Falco tinnunculus | 77.45 | 78.71 | 75.52 | 612 | zip | ~ L |
| Eurasian Hobby | Falco subbuteo | 97.56 | 97.86 | 97.10 | 614 |  |  |
| Peregrine Falcon | Falco peregrinus | 99.67 | 99.46 | 100.00 | 614 |  |  |
| Corn Crake | Crex crex | 99.84 | 100.00 | 99.59 | 614 |  |  |
| Water Rail | Rallus aquaticus | 98.86 | 98.12 | 100.00 | 614 |  |  |
| Eurasian Moorhen | Gallinula chloropus | 87.44 | 81.99 | 95.85 | 613 |  |  |
| Eurasian Coot | Fulica atra | 89.89 | 86.56 | 95.02 | 613 |  |  |
| Northern Lapwing | Vanellus vanellus | 81.11 | 69.97 | 98.34 | 614 |  |  |
| Little Ringed Plover | Charadrius dubius | 97.88 | 96.51 | 100.00 | 614 |  |  |
| Eurasian Oystercatcher | Haematopus ostralegus | 98.53 | 97.59 | 100.00 | 614 |  |  |
| Eurasian Curlew | Numenius arquata | 98.70 | 97.86 | 100.00 | 614 |  |  |
| Eurasian Woodcock | Scolopax rusticola | 86.21 | 89.40 | 81.33 | 609 | zip | ~ 1 |
| Mew Gull | Larus canus | 99.35 | 98.93 | 100.00 | 614 |  |  |
| Herring Gull | Larus argentatus | 99.51 | 99.20 | 100.00 | 614 |  |  |
| Common Pigeon | Columba livia f. domestica | 92.83 | 88.74 | 99.17 | 614 |  |  |
| Stock Dove | Columba oenas | 65.58 | 59.14 | 75.52 | 613 | pois | ~ 1 |
| Common Wood Pigeon | Columba palumbus | 1.14 | 1.34 | 0.83 | 613 | nb | ~ 1 |
| European Turtle Dove | Streptopelia turtur | 90.38 | 89.78 | 91.29 | 613 |  |  |
| Eurasian Collared Dove | Streptopelia decaocto | 63.40 | 49.33 | 85.06 | 612 | zinb | ~ L |
| Rose-ringed Parakeet | Psittacula krameri | 97.72 | 96.25 | 100.00 | 614 |  |  |
| Common Cuckoo | Cuculus canorus | 85.13 | 79.25 | 94.19 | 612 |  |  |
| Barn Owl | Tyto alba | 90.38 | 84.95 | 98.76 | 613 |  |  |
| Eurasian Eagle-Owl | Bubo bubo | 98.05 | 99.46 | 95.85 | 614 |  |  |
| Eurasian Pygmy Owl | Glaucidium passerinum | 98.70 | 99.73 | 97.10 | 614 |  |  |
| Little Owl | Athene noctua | 83.17 | 74.12 | 97.10 | 612 |  |  |
| Tawny Owl | Strix aluco | 64.05 | 69.27 | 56.02 | 612 | zip | ~ (1\|ID) |
| Long-eared Owl | Asio otus | 87.62 | 90.88 | 82.57 | 614 |  |  |
| Boreal Owl | Aegolius funereus | 98.37 | 100.00 | 95.85 | 614 |  |  |
| European Nightjar | Caprimulgus europaeus | 99.84 | 100.00 | 99.59 | 614 |  |  |
| Common Swift | Apus apus | 81.01 | 79.78 | 82.92 | 611 | zip | ~ 1 |
| Common Kingfisher | Alcedo atthis | 93.16 | 93.03 | 93.36 | 614 |  |  |
| Eurasian Wryneck | Jynx torquilla | 99.51 | 99.73 | 99.17 | 614 |  |  |
| Middle Spotted Woodpecker | Dendrocoptes medius | 91.35 | 90.05 | 93.36 | 613 |  |  |
| Lesser Spotted Woodpecker | Dendrocopos minor | 90.05 | 87.90 | 93.36 | 613 |  |  |
| Great Spotted Woodpecker | Dendrocopos major | 20.42 | 21.83 | 18.26 | 612 | zinb | ~ (1\|ID) |
| Black Woodpecker | Dryocopus martius | 86.81 | 92.23 | 78.42 | 614 |  |  |
| European Green Woodpecker | Picus viridis | 59.80 | 50.67 | 73.86 | 612 | pois | ~ 1 |
| Grey-headed Woodpecker | Picus canus | 95.77 | 100.00 | 89.21 | 614 |  |  |
| Red-backed Shrike | Lanius collurio | 86.48 | 95.17 | 73.03 | 614 |  |  |
| Great Grey Shrike | Lanius excubitor | 99.67 | 100.00 | 99.17 | 614 |  |  |
| Eurasian Golden Oriole | Oriolus oriolus | 94.30 | 90.88 | 99.59 | 614 |  |  |
| Eurasian Jay | Garrulus glandarius | 23.37 | 27.22 | 17.43 | 612 | zinb | ~ PCs |
| Eurasian Magpie | Pica pica | 35.78 | 28.84 | 46.47 | 612 | zinb | ~ (1\|ID) |
| Spotted Nutcracker | Nucifraga caryocatactes | 98.53 | 100.00 | 96.27 | 614 |  |  |
| Western Jackdaw | Corvus monedula | 70.75 | 54.45 | 95.85 | 612 |  |  |
| Rook | Corvus frugilegus | 98.86 | 98.12 | 100.00 | 614 |  |  |
| Carrion Crow | Corvus corone | 19.25 | 14.25 | 26.97 | 613 | zinb | ~ PCs |
| Northern Raven | Corvus corax | 98.37 | 99.46 | 96.68 | 614 |  |  |
| Eurasian Skylark | Alauda arvensis | 58.79 | 54.69 | 65.15 | 614 | zinb | ~ 1 |
| Woodlark | Lullula arborea | 96.91 | 95.71 | 98.76 | 614 |  |  |
| Sand Martin | Riparia riparia | 99.84 | 99.73 | 100.00 | 614 |  |  |
| Barn Swallow | Hirundo rustica | 51.71 | 51.08 | 52.70 | 613 | zinb | ~ L |
| Common House Martin | Delichon urbicum | 68.19 | 71.77 | 62.66 | 613 | zinb | ~ (1\|ID) |
| Marsh Tit | Poecile palustris | 36.99 | 43.51 | 26.97 | 611 | zinb | ~ (1\|ID) |
| Willow Tit | Poecile montanus | 62.70 | 76.94 | 40.66 | 614 | zip | ~ 1 |
| Coal Tit | Periparus ater | 54.56 | 78.02 | 18.26 | 614 | nb | ~ 1 |
| European Crested Tit | Lophophanes cristatus | 62.05 | 79.36 | 35.27 | 614 | zinb | ~ PCs |
| Great Tit | Parus major | 0.65 | 0.54 | 0.83 | 611 | zinb | ~ 1 |
| Eurasian Blue Tit | Cyanistes caeruleus | 1.47 | 1.08 | 2.07 | 611 | nb | ~ 1 |
| Eurasian Penduline Tit | Remiz pendulinus | 99.51 | 99.20 | 100.00 | 614 |  |  |
| Long-tailed Tit | Aegithalos caudatus | 44.12 | 38.27 | 53.11 | 612 | zip | ~ PCs |
| Eurasian Nuthatch | Sitta europaea | 24.55 | 30.54 | 15.35 | 611 | pois | ~ 1 |
| Eurasian Treecreeper | Certhia familiaris | 73.62 | 93.83 | 42.32 | 614 |  |  |
| Short-toed Treecreeper | Certhia brachydactyla | 22.09 | 15.41 | 32.37 | 611 | zinb | ~ PCs |
| Eurasian Wren | Troglodytes troglodytes | 1.96 | 2.96 | 0.41 | 612 | nb | ~ 1 |
| White-throated Dipper | Cinclus cinclus | 94.79 | 99.20 | 87.97 | 614 |  |  |
| Common Goldcrest | Regulus regulus | 42.55 | 59.84 | 15.83 | 611 | zinb | ~ 1 |
| Firecrest | Regulus ignicapilla | 48.20 | 67.47 | 18.33 | 612 | zinb | ~ (1\|ID) |
| Willow Warbler | Phylloscopus trochilus | 26.63 | 33.60 | 15.83 | 612 | nb | ~ 1 |
| Common Chiffchaff | Phylloscopus collybita | 0.49 | 0.81 | 0.00 | 611 | nb | ~ 1 |
| Wood Warbler | Phylloscopus sibilatrix | 79.15 | 93.57 | 56.85 | 614 |  |  |
| Greenish Warbler | Phylloscopus trochiloides | 99.84 | 100.00 | 99.59 | 614 |  |  |
| Melodious Warbler | Hippolais polyglotta | 99.84 | 99.73 | 100.00 | 614 |  |  |
| Icterine Warbler | Hippolais icterina | 74.18 | 67.92 | 83.82 | 612 | zinb | ~ 1 |
| Sedge Warbler | Acrocephalus schoenobaenus | 99.84 | 99.73 | 100.00 | 614 |  |  |
| Eurasian Reed Warbler | Acrocephalus scirpaceus | 95.11 | 92.49 | 99.17 | 614 |  |  |
| Marsh Warbler | Acrocephalus palustris | 62.25 | 57.68 | 69.29 | 612 | zinb | ~ L |
| Common Grasshopper Warbler | Locustella naevia | 93.16 | 93.03 | 93.36 | 614 |  |  |
| Eurasian Blackcap | Sylvia atricapilla | 0.49 | 0.54 | 0.41 | 610 | nb | ~ 1 |
| Garden Warbler | Sylvia borin | 14.78 | 14.63 | 15.00 | 609 | zinb | ~ 1 |
| Lesser Whitethroat | Sylvia curruca | 61.76 | 63.34 | 59.34 | 612 | zip | ~ 1 |
| Common Whitethroat | Sylvia communis | 25.16 | 17.79 | 36.51 | 612 | zinb | ~ 1 |
| Spotted Flycatcher | Muscicapa striata | 51.06 | 45.41 | 59.75 | 611 | zip | ~ (1\|ID) |
| European Robin | Erithacus rubecula | 3.11 | 4.86 | 0.41 | 611 | zinb | ~ 1 |
| Common Nightingale | Luscinia megarhynchos | 83.66 | 75.74 | 95.85 | 612 |  |  |
| European Pied Flycatcher | Ficedula hypoleuca | 82.19 | 83.02 | 80.91 | 612 | zip | ~ (1\|ID) |
| Common Redstart | Phoenicurus phoenicurus | 85.43 | 81.08 | 92.12 | 611 |  |  |
| Black Redstart | Phoenicurus ochruros | 20.59 | 18.33 | 24.07 | 612 | zip | ~ 1 |
| Whinchat | Saxicola rubetra | 99.67 | 99.46 | 100.00 | 614 |  |  |
| European Stonechat | Saxicola rubicola | 94.95 | 92.23 | 99.17 | 614 |  |  |
| Common Blackbird | Turdus merula | 0.00 | 0.00 | 0.00 | 611 | zinb | ~ 1 |
| Fieldfare | Turdus pilaris | 71.52 | 87.03 | 47.72 | 611 | zinb | ~ PCs |
| Song Thrush | Turdus philomelos | 2.45 | 4.05 | 0.00 | 611 | nb | ~ 1 |
| Mistle Thrush | Turdus viscivorus | 41.24 | 48.65 | 29.88 | 611 | zip | ~ 1 |
| Common Starling | Sturnus vulgaris | 21.80 | 15.14 | 32.08 | 610 | nb | ~ 1 |
| Dunnock | Prunella modularis | 1.47 | 1.08 | 2.07 | 612 | zinb | ~ 1 |
| Western Yellow Wagtail | Motacilla flava | 75.86 | 65.86 | 91.29 | 613 |  |  |
| Grey Wagtail | Motacilla cinerea | 80.75 | 87.37 | 70.54 | 613 | zip | ~ PCs |
| White Wagtail | Motacilla alba | 16.64 | 13.44 | 21.58 | 613 | zinb | ~ 1 |
| Meadow Pipit | Anthus pratensis | 96.09 | 95.17 | 97.51 | 614 |  |  |
| Tree Pipit | Anthus trivialis | 75.86 | 81.18 | 67.63 | 613 | nb | ~ 1 |
| Yellowhammer | Emberiza citrinella | 25.33 | 30.46 | 17.43 | 612 | nb | ~ 1 |
| Common Reed Bunting | Emberiza schoeniclus | 95.11 | 91.96 | 100.00 | 614 |  |  |
| Corn Bunting | Emberiza calandra | 98.86 | 98.12 | 100.00 | 614 |  |  |
| Common Chaffinch | Fringilla coelebs | 0.16 | 0.27 | 0.00 | 611 | zinb | ~ PCs |
| European Greenfinch | Chloris chloris | 15.66 | 10.48 | 23.65 | 613 | zinb | ~ 1 |
| Red Crossbill | Loxia curvirostra | 90.54 | 99.19 | 77.18 | 613 |  |  |
| Lesser Redpoll | Acanthis cabaret | 94.95 | 97.32 | 91.29 | 614 |  |  |
| Eurasian Siskin | Spinus spinus | 95.77 | 99.46 | 90.04 | 614 |  |  |
| Common Linnet | Carduelis cannabina | 57.19 | 60.92 | 51.45 | 612 | zip | ~ (1\|ID) |
| European Goldfinch | Carduelis carduelis | 45.19 | 37.37 | 57.26 | 613 | zinb | ~ (1\|ID) |
| European Serin | Serinus serinus | 83.03 | 89.25 | 73.44 | 613 | nb | ~ 1 |
| Eurasian Bullfinch | Pyrrhula pyrrhula | 52.53 | 68.01 | 28.63 | 613 | zip | ~ PCs |
| Hawfinch | Coccothraustes coccothraustes | 64.76 | 71.85 | 53.75 | 613 | zinb | ~ 1 |
| House Sparrow | Passer domesticus | 23.40 | 16.22 | 34.44 | 611 | zinb | ~ (1\|ID) |
| Eurasian Tree Sparrow | Passer montanus | 56.79 | 55.41 | 58.92 | 611 | zinb | ~ 1 |
